# InStrain enables population genomic analysis from metagenomic data and rigorous detection of identical microbial strains

**DOI:** 10.1101/2020.01.22.915579

**Authors:** Matthew R. Olm, Alexander Crits-Christoph, Keith Bouma-Gregson, Brian Firek, Michael J. Morowitz, Jillian F. Banfield

## Abstract

Coexisting microbial cells of the same species often exhibit genetic differences that can affect phenotypes ranging from nutrient preference to pathogenicity. Here we present inStrain, a program that utilizes metagenomic paired reads to profile intra-population genetic diversity (microdiversity) across whole genomes and compare populations in a microdiversity-aware manner, dramatically increasing genomic comparison accuracy when benchmarked against existing methods. We use inStrain to profile >1,000 fecal metagenomes from newborn premature infants and find that siblings share significantly more strains than unrelated infants, although identical twins share no more strains than fraternal siblings. Infants born via cesarean section harbored *Klebsiella* with significantly higher nucleotide diversity than infants delivered vaginally, potentially reflecting acquisition from hospital versus maternal microbiomes. Genomic loci showing diversity within an infant included variants found in other infants, possibly reflecting inoculation from diverse hospital-associated sources. InStrain can be applied to any metagenomic dataset for microdiversity analysis and rigorous strain comparison.

## Main

Cells in microbial populations are not all identical to one another. Genetic polymorphisms rapidly arise through *de novo* mutation, and these variants can spread because they confer a fitness advantage or by lateral gene transfer (if the variant confers an advantage or is linked to a fitness-conferring variant). It is estimated that billions to trillions of bacterial genetic mutations are generated *de novo* every day in the microbiome of an individual adult human ^1^, and these differences can be clinically relevant. For example, just three point mutations can confer antibiotic resistance in *Enterobacteriaceae* ^2^. Studying genetic variation in microbial populations has historically involved isolating a multitude of cells from the same population and performing phenotypic analysis and/or genome sequencing. Genome-resolved metagenomic analysis, which involves extracting and sequencing DNA directly from the environment and using computational tools to assemble and bin the resulting DNA sequences into genomes *in silico*, presents an attractive high-throughput alternative to this process. This technique allows simultaneous analysis of microbial communities, the species populations that comprise them, and heterogeneity within these populations, and has been used to reveal fine-scale evolutionary mechanisms ^3–5^, dynamics ^6–12^, and strain level metabolic variation that could contribute to strain selection ^1, 13^.

Many fundamental questions in human microbiome research relate to the transmission of microbial populations between individuals, including how we are seeded by microbes early in life ^14–16^. However, strain diversity presents challenges for such analyses. Sequence comparisons are usually performed by aligning consensus genomes assembled from different samples ^1, 17^ or by modifying a reference genome using mapped reads and comparing it to the same sequence that has been modified by reads from another sample ^18–20^ **(Supplemental Figure S1)**. These methods represent each population based on the most common alleles, which can lead to erroneous results. For example, if sample 1 contains a single nucleotide variant (SNV) A at 20% frequency and T (the consensus choice) at 80% frequency, and sample 2 has A at 100% frequency, comparing the consensus genome of both samples will fail to identify the variant shared by both populations. Further, alleles at intermediate frequencies (e.g. 30% - 70%) can be stochastically detected above or below 50% due to random sampling, resulting in chimeric consensus sequences. As natural microbial populations can have many polymorphic sites, genomic comparison methods that consider the genetic diversity are needed, as are standardized methods that are easy to use and that are applicable to all metagenomic studies.

Here we present inStrain, a program that profiles population microdiversity from metagenomic short read alignments and performs microdiversity-aware genomic comparisons. This includes calculating nucleotide diversity and linkage disequilibrium, identifying SNVs (including non-synonymous and synonymous variants), and reporting accurate coverage depth and breadth. We demonstrate that inStrain performs strain-level comparisons with higher accuracy and sensitivity than leading tools. To demonstrate the value of inStrain for microbiome studies, we apply inStrain to a large collection of previously sequenced infant fecal microbiomes to reveal patterns of microbiome microdiversity and strain sharing among infants born in the same neonatal intensive care unit (NICU) over a period of five years. inStrain is available as an open-source python program on GitHub (https://github.com/MrOlm/inStrain) and documentation is available both in the supplemental materials (**Supplemental Document S1)** and online at https://instrain.readthedocs.io/en/latest/.

## Results

### inStrain measures population-level diversity from metagenomic data

InStrain profiles the microdiversity of any DNA sequence dataset that consists of paired short reads that are mapped to a genome assembled from a metagenome or from a cultured isolate. Functionality can be broken into three major steps:

*Step 1) Read filtering*. To increase the likelihood that mapped read pairs originate from organisms belonging to the same population a series of filters are applied. For each read pair aligned to the reference genome (*de novo* assembled from the same sample or a genome from another source) the mapQ score, average nucleotide identity (ANI) of the pair to the reference genome, and the insert size between aligned reads are calculated. Read pairs that don’t pass adjustable quality cutoffs are removed, as are all unpaired reads. The exclusive use of pairs doubles the number of bases used to calculate the read ANI and mapQ score, increasing their accuracy and substantially increasing the span of genome analyzed. This reduces mismapping at repeat regions or regions conserved in multiple genomes. Other software tools, such as StrainPhlAn and MetaPhlAn ^18, 21^, treat pairs of reads as separate observations and can assign each read pair to a different population, contrary to the strong expectation from Illumina sequencing protocols that a pair originates from a single DNA molecule.

*Step 2) Calculation of nucleotide diversity, SNVs, and linkage*. For each gene, scaffold, and/or genome, inStrain calculates the mean, median, and standard deviation of the depth of coverage (number of reads per base-pair), breadth of coverage (percentage of reference base pairs covered by at least one read), expected breadth of coverage (given the average depth of coverage, the breadth of coverage that would be expected if reads were evenly spread across the genome), and average nucleotide diversity (π; ^22^) of all base-pairs with at least 5x coverage (**Figure 1a**). Both bialllelic and multiallelic SNVs and their frequencies are identified and annotated at positions where phred30 quality filtered reads differ from the reference genome and at positions where multiple bases are simultaneous detected at levels above the expected sequencing error rate. SNVs are classified as synonymous, non-synonymous, or intergenic based on gene annotations, and linkage disequilibrium is calculated between SNVs that are connected by at least twenty read-pairs.

**Figure 1.**
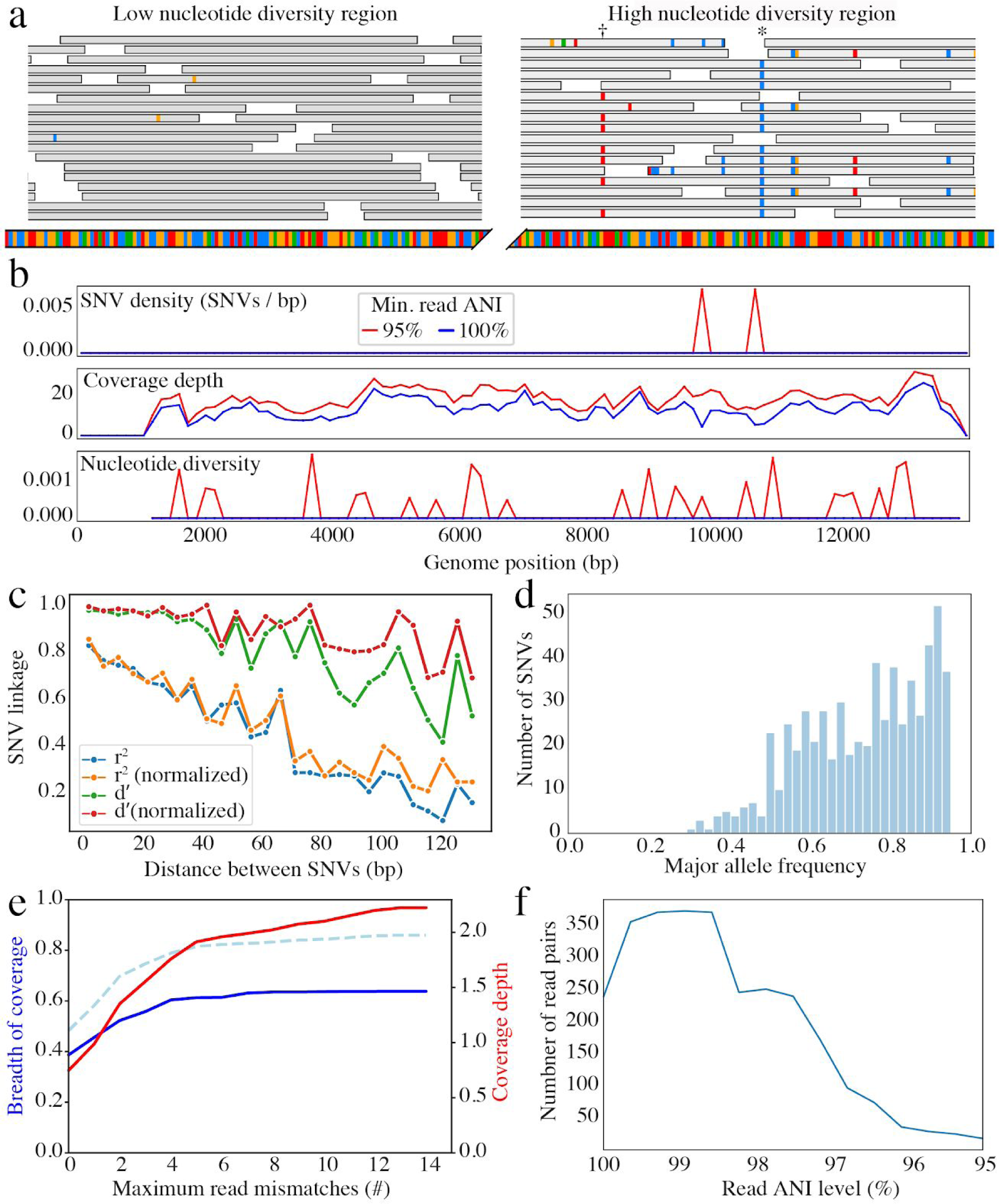
InStrain measures population-level diversity from metagenomic data. a) Examples of metagenomic reads (grey boxes) mapping to genomic regions with low and high nucleotide diversity. Mismatches to the reference genome are represented by small colored marks on the reads, and the reference genome is represented below the reads. b-f) Examples of figures automatically generated by inStrain. b) SNV density, coverage, and nucleotide diversity across a bacteriophage genome. Spikes in nucleotide diversity and SNV density do not correspond with increased coverage, indicating that the signals are not due to read mis-mapping. Positions with nucleotide diversity and no SNV-density are those where diversity exists but is not high enough to call a SNV c) Metrics of SNV linkage vs. distance between SNVs; linkage decay (as shown here) is a common signal of recombination. d) Distribution of the major allele frequencies of bi-allelic SNVs (the Site Frequency Spectrum). Alleles with major frequencies below 50% are the result of multiallelic sites. The lack of distinct puncta suggest that more than a few distinct strains are present. e) Breadth of coverage (blue line), coverage depth (red line), and expected breadth of coverage given the depth of coverage (dotted blue line) versus the minimum ANI of mapped reads. Coverage depth continues to increase while breadth plateaus, suggesting that all regions of the reference genome are not present in the reads being mapped. f) Distribution of read pair ANI levels when mapped to a reference genome; this plot suggests that the reference genome is >1% different than the mapped reads.

*Step 3) Generation of tables and figures*. Tables are generated that describe how many reads were removed by each filter described in *Step 1* and enumerate all metrics described in *Step 2*. Figures are generated for each genome to document SNV allele frequencies, genome-wide nucleotide diversity, patterns of linkage disequilibrium, and to report other findings (**Figure 1b-f**). All data generated during an inStrain run is stored in a space-efficient manner and can be used to quickly re-generate plots and tables with different parameters.

### Microdiversity-aware ANI calculations (PopANI) increase accuracy of strain discrimination

Most existing strain-comparison pipelines compare microbes in different samples based on their consensus genomes. In contrast, inStrain considers both major and minor alleles during genomic comparison. This new microdiversity-aware ANI metric is referred to as “PopANI” (population-level ANI), and it is reported alongside consensus-based ANI (“ConANI”). Both metrics are calculated in a pair-wise manner for samples that have been profiled using the methods described above. First, all positions of the genome that have at least 5x coverage in both samples are identified. Only these positions are considered in the PopANI and ConANI calculations. Second, the number of positions with ≥5x coverage that differ in allelic composition between the samples is enumerated. For ConANI, if the consensus base differs between the two samples a substitution is called. For PopANI, a substitution is called at a site only if both samples share no alleles (either major or minor) **(Figure 2a**).

**Figure 2.**
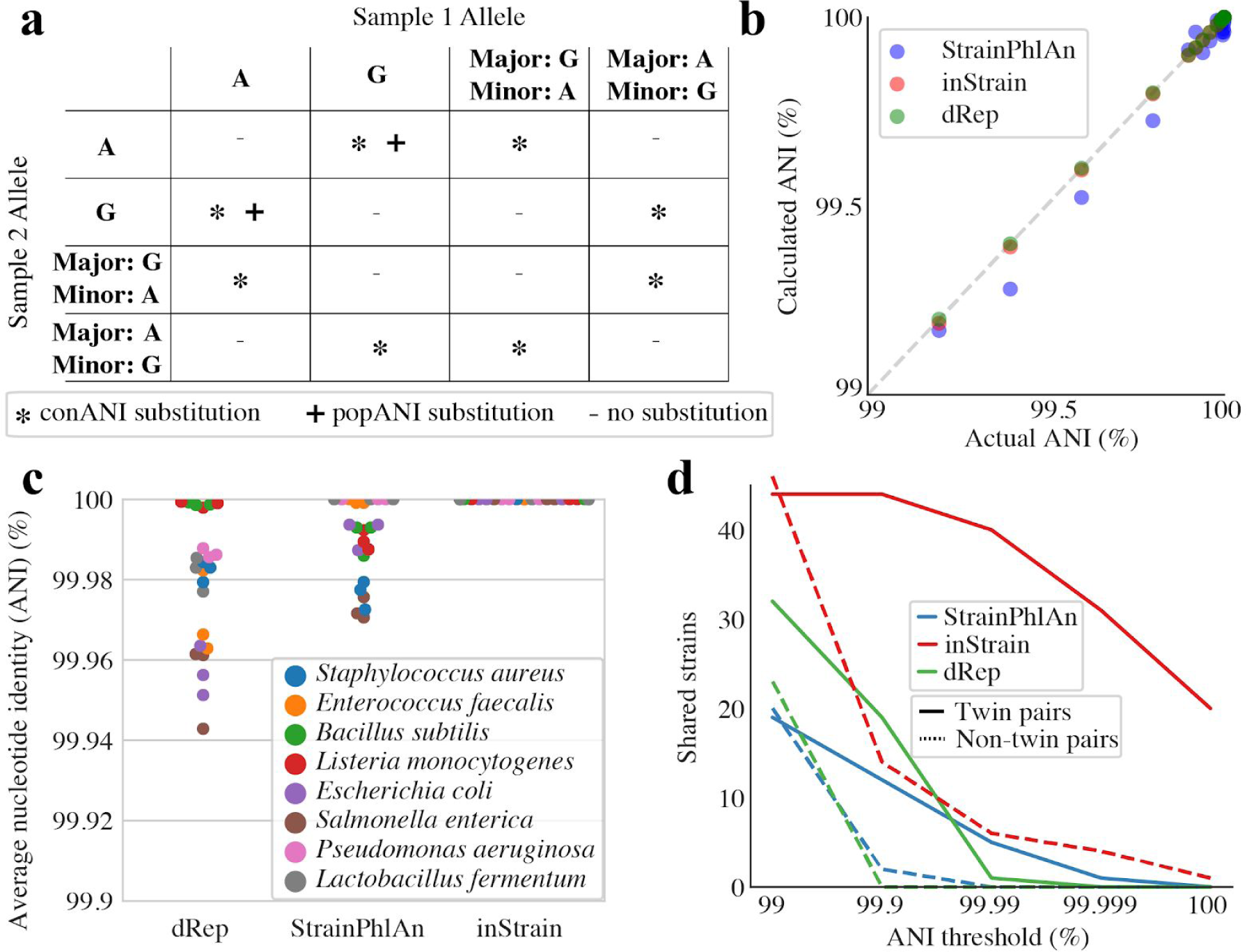
InStrain accurately discriminates between closely related strains. a) Table demonstrating the circumstances under which conANI and popANI substitutions will be called. ConANI substitutions are called whenever the consensus base differs, and popANI substitutions are only called when there is no allelic overlap between samples. b) Synthetic mutations were introduced to a reference genome of *E. coli* obtained from RefSeq to generate variant genomes with specific ANI differences from the reference genome, and three tools were used to compare the variant genomes to the reference genome. dRep and inStrain consistently reported accurate ANI values, while StrainPhlAn was inaccurate by a median of 0.03% ANI. c) A mock community of bacterial cells was sequenced in biological triplicate and compared using three tools. InStrain performed best in correctly identifying that the genomes were identical in all three samples. d) The fecal microbiomes of three sets of twins were compared using each of the three tools, and the number of bacterial genomes with ANI values above a range of thresholds is plotted for pairs of twins (which are expected to share more strains) and pairs of unrelated infants. InStrain remained sensitive at higher ANI thresholds than the other two tools.

We benchmarked inStrain’s strain comparison method against two existing common tools: dRep, which calculates genome-wide ANI ^17^, and StrainPhlAn ^18^, which aligns short reads to a marker gene database (0.3% of the genome in the case of *Escherichia coli*) and compares the consensus maker genes in multiple samples. We first compared the ability of each method to report the ANI between genomes with a known number of *in silico* mutations **(Figure 2b)**. All three methods performed well on this test, which does not consider microdiversity, though dRep and inStrain had lower errors in the ANI calculation than StrainPhlAn overall (0.00001%, 0.002%, and 0.03%, respectively; average discrepancy between the true and calculated ANI). This is likely because dRep and inStrain compare positions from across the entire genome (99.9% and 99.7% of the genome, respectively) and StrainPhlAn does not.

We next used each tool to compare metagenomes derived from defined bacterial communities. The ZymoBIOMICS Microbial Community Standard, which contains cells from eight bacterial species at defined abundances, was divided into three aliquots and subjected to DNA extraction, library preparation, and metagenomic sequencing. Each strain comparison tool was then used to compare bacterial species in each sample to each other in a pairwise manner **(Figure 2c).** As all genomic comparisons originate from the same defined community of microbes, each tool should report 100% ANI for all genomic comparisons. Deviations from this ideal either represent errors in sequence alignment or the presence of microdiversity that is likely present because cultures have been maintained in the laboratory. dRep and StrainPhlAn reported average ANI values of 99.98%, 99.99% whereas inStrain reported average popANI values of 100% for 23 of the 24 comparisons and 99.99996% for one comparison. The difference in performance arises because the Zymo cultures contain non-fixed nucleotide variants that inStrain uses to confirm population overlap but that confuse the consensus sequences reported by dRep and StrainPhlAn.

We used the Zymo data to establish a threshold for the detection of “same” versus “different” strains. The thresholds for dRep, StrainPhlAn and inStrain, calculated based on the lowest average ANI across all 24 sequence comparisons, were 99.94% ANI, 99.97% ANI, and 99.99996% ANI, respectively. Thus, inStrain can be used for detection of identical microbial strains with a stringency that is substantially higher than either other tool. Using the previously reported rate of 0.9 SNPs accumulated per genome per year in the gut microbiome of healthy human adults ^1^, in this test dRep is able to discriminate between strains that have diverged for at least 2,528 years, StrainPhlAn for 1,307 years, and inStrain for 2.2 years (**Supplemental Table S1**). Stringent thresholds are useful for strain tracking, as strains that have diverged for hundreds to thousands of years are clearly not linked by a recent transmission event.

To compare the ability of the three methods to detect strains shared by twin premature infants, the microbiomes of six infants were processed according to the best recommended practice for each of the three tools. We then compared the number of strains found to be shared by twins and non-twins over a range of ANI thresholds. All methods identified significantly more strain sharing among twin pairs than pairs of unrelated infants, as expected, but inStrain and dRep identified substantially more shared strains than StrainPhlAn and inStrain remained sensitive at substantially higher ANI thresholds than either of the other tools **(Figure 2d)**. We attribute the reduced ability of StrainPhlAn to identify shared strains to: (1) StrainPhlAn’s relies on a database of species-specific reference marker genes whereas inStrain and dRep use reference genomes assembled from the samples themselves. This can lead to failure to detect strains that are not sufficiently closely related to those in the reference database. For example, although inStrain identified 55 bacteriophage strains and 20 plasmid strains that were shared between at least two infants, StrainPhlAn detected 0, likely reflecting their poor coverage in reference databases. **(Supplemental Table S1)**. (2) Erroneous read mapping due to failure to consider paired read information. Reads that can be mapped to two genomes equally well, such as those coming from conserved regions of the genome, are randomly assigned to one genome and can corrupt the consensus sequence ^23^. InStrain’s use of paired read information substantially reduces this problem. (3) StrainPhlAn is able to detect only one strain per species in any sample, yet we know that microbiomes can contain multiple coexisting strains. When two or more strains of a species are in a sample at similar abundance levels, this can lead to pileups of reads from multiple strains and chimeric sequences. In combination, the reduced ability to detect truly shared strains and the limitations at high ANI thresholds needed to distinguish “same” from “different” limit the utility of the previously available tools for strain tracking.

### Siblings share significantly more microbial strains at birth than unrelated infant pairs

We next applied inStrain to 1,163 fecal metagenomes from 160 premature infants born into the same neonatal intensive care unit ^24^. The dataset includes samples from six individual sampling campaigns, involved the enrollment of 6 sets of monozygotic twins (MZ; identical), 20 sets of dizygotic twins (DZ; fraternal) and 3 sets of trizygotic (TZ) triplets, and over eight thousand *de novo* genomes from bacteria, bacteriophage, and plasmid colonists. Organisms that may have been introduced through contamination were removed based on their presence in sequenced negative controls, each genome set was de-replicated at 98% ANI to form “sub-species” groups, and representative genomes from each sub-species were combined into a single mapping database consisting of 2,266 genomes in order to reduce multi-mapped reads **(Supplemental Figure S2)**. All metagenomes were mapped to this dereplicated genome set and inStrain was used to profile the microdiversity of each mapping. In all cases where a sub-species was detected in multiple infants with over 50% breadth of coverage, inStrain was used to compare strains.

A threshold of 99.999% popANI was chosen as the threshold to define bacterial, bacteriophage, and plasmid strains as being the same “strain” based on the Zymo experiment (**Figure 2c**) and comparisons between subspecies present in the same infant over time (based on the assumption that strain genotypes from samples collected within days or weeks of each other typically represent the same strain) **(Supplemental Figure S3)**. Thus, to be classified as the same strain, two populations must have no fixed differences within this margin of error. Of the 109,731 comparisons made, 4,103 (grey lines in **Figure 3a**) indicated that infants shared bacterial strains. Of these, 268 cases revealed sharing between pairs of siblings (despite sibling pair comparisons comprising only 0.3% of all comparisons; red lines in **Figure 3a)**. Further, the majority of bacterial strains that were identified in two and only two infants were shared between sibling pairs **(Figure 3b,c)**. Similar patterns were identified for bacteriophage and plasmid colonists **(Supplemental Table S2)**.

**Figure 3.**
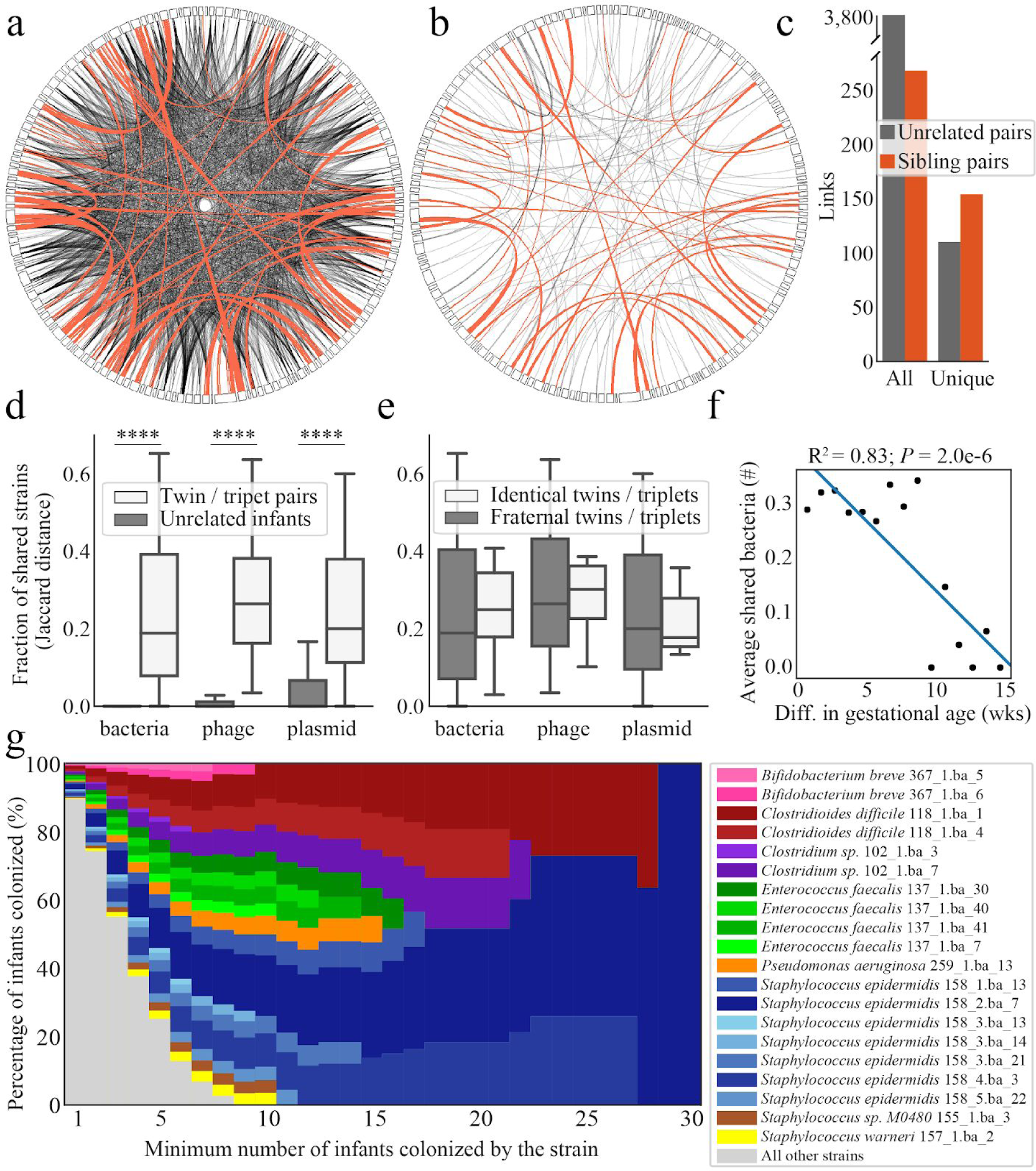
Siblings share significantly more microbial strains at birth than unrelated infants. a,b) A link is drawn for each strain shared between pairs of infants (represented by rectangles along the circumferences). Links between sibling pairs are drawn in red, links between unrelated infants are drawn in grey. Diagrams are made displaying all strains (a) and only strains that are uniquely in two and only two infants (b). c) Enumeration of links drawn in (a) and (b). d) Twin pairs share significantly more strains of all domains than unrelated pairs (**** = *p* < 1e-15). e) Identical twin pairs do not share significantly more strains than fraternal twin pairs. f) Infants born more closely in gestational age share significantly more bacterial strains. g) Most strains colonize only a single infant, but some strains colonize many more. For each minimum number of infants colonized (x-axis), the percentage of total infant colonizations by strains above that threshold is shown. The top eight microbial species are assigned a color, and highly colonizing strains (≥ 5 infants) of each species are assigned a shade of that color.

The majority of bacterial strains identified in this study were detected in only a single infant (1818 of 3044 strains). The most frequently colonizing strain (*Staphylococcus epidermidis* 158.2.ba_7) was identified in samples from 49 of the 160 infants. Six of the seven other most frequently colonizing species were also Firmicutes, and many are known for their role in nosocomial infections, including *Clostridioides difficile* and *Enterococcus faecalis*. *Pseudomonas aeruginosa*, a Proteobacterium, is also implicated in nosocomial infections. Twelve strains colonized more than ten infants, including five strains of *S. epidermidis*, three strains of *E. faecalis,* two strains of *C. difficile*, and one strain each of *P. aeruginosa* and *Clostridium sp.* **(Figure 3g)**. These frequently encountered strains may have specific adaptations that enable them to survive in the neonatal intensive care unit (NICU). Alternatively, they may be acquired from health care workers that commonly interact with these infants.

Overall, siblings shared significantly more strains of bacteria, bacteriophage, and plasmids than unrelated infant pairs (**Figure 3d)**. However, among siblings, monozygotic (MZ) twins shared no more strains than dizygotic (DZ) twins and trizygotic (TZ) triplets **(Figure 3e)**. Infants born at more chronologically similar times shared significantly more strains of bacteriophages and plasmids, supporting the role of the hospital room environment in shaping initial bacteriophage and plasmid strain acquisition **(Supplemental Figure S4)**. Infants born with similar gestational ages and birth weights also shared significantly more strains of bacteria, bacteriophages, and plasmids than those with different ages and weights (**Figure 3f**; **Supplemental Figure S4)**. In combination, the results point to the role of infant physiology, sibling status, and calendar date of birth (i.e., similar date of residence in the NICU) in strain acquisition.

### Nucleotide diversity of the premature infant microbiome

Over the sampling time-series in this study (generally the first few months of life), hospitalized infants were colonized by an average of 17.8 ± 0.7 sub-species of bacteria, 26.9 ± 1.5 sub-species of bacteriophage, and 7.4 ± 0.3 sub-species of plasmids per infant (mean ± SEM; colonization defined as detection of genome at >5x depth coverage across ≥ 50% of the genome) (**Supplemental Table S3**). As the 160 infants were sampled over six different campaigns, each using a unique combination of library preparation methodology, Illumina machine for sequencing, and institutional sequencing center, we first tested for effects related to sampling campaign. Infants of the same campaign were not more likely to share strains **(Supplemental Figure S4)**, but measured nucleotide diversity among colonists varied significantly between the six different sampling campaigns, primarily driven by differences in library preparation methodology and the DNA sequencing machine used **(Supplemental Figure S5).** We thus analyzed each cohort separately for relationships between microdiversity and infant metadata, allowing us to validate the consistency of inStrain when run using different sequencing methodologies.

Bacteria had significantly higher nucleotide diversity than plasmids and phage in 4/6 campaigns, whereas plasmids had the lowest nucleotide diversity in 4/6 campaigns **(Supplemental Figure S5)**. Relative to other bacteria, Proteobacteria had significantly higher and Firmicutes significantly lower nucleotide diversity in 3/6 and in 4/6 campaigns, respectively **(Supplemental Table S2)**. Approximately 75% of premature infants were born via cesarean section (118/160), and their bacterial colonists had significantly higher nucleotide diversity than vaginally delivered infants in the NIH4 and Sloan2 cohorts and overall **(Figure 4a)**. This effect was particularly striking for *Klebsiella* **(Figure 4b)**, and the difference remained significant even when excluding infants in the NIH4 and Sloan2 cohorts **(Supplemental Figure S4)**.

**Figure 4.**
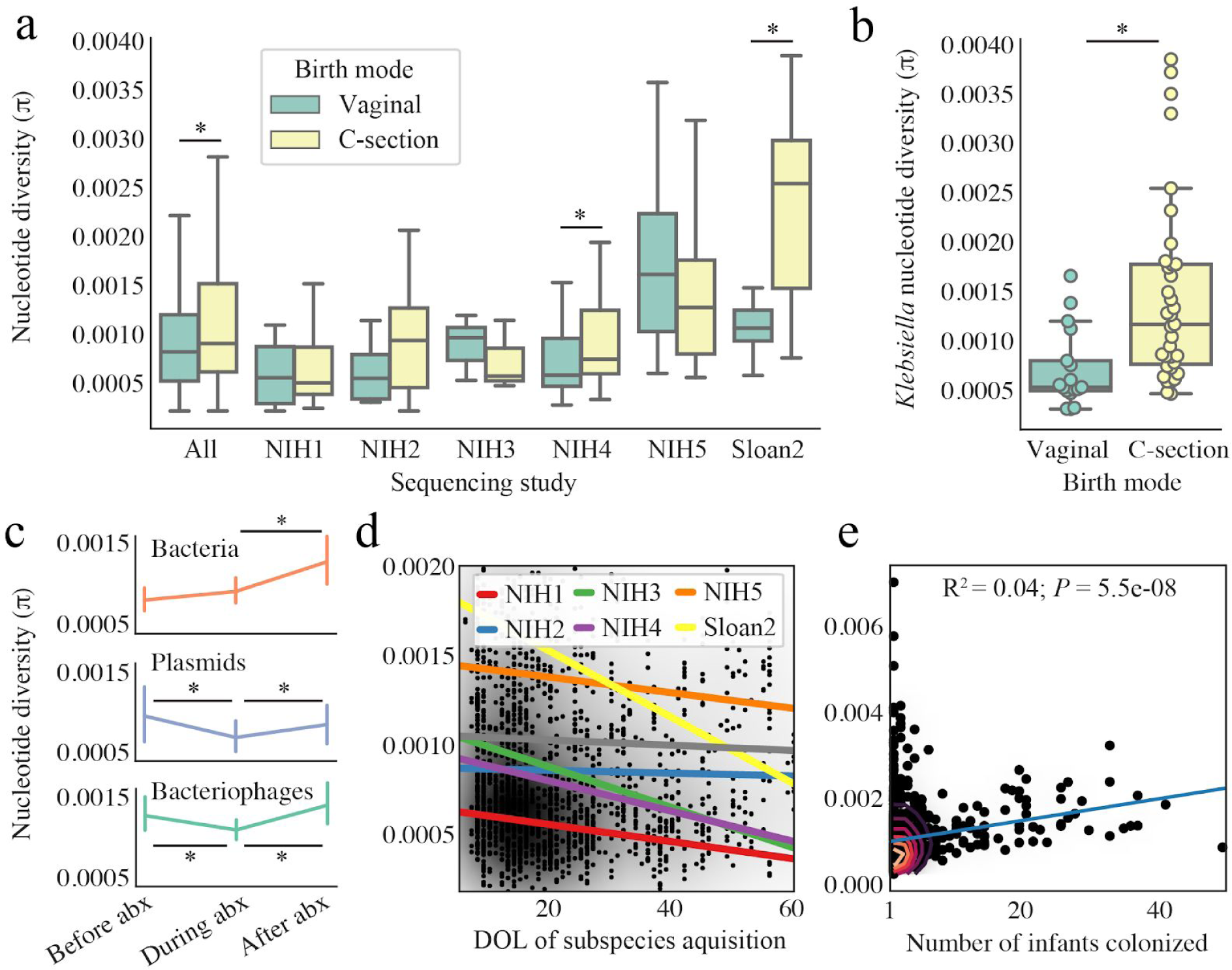
Analysis of the microdiversity of premature infant colonists. a) Overall and among two of the six individual study cohorts, infants born via C-section had host microbes with higher nucleotide diversity than those delivered vaginally (* = *p* < 0.05). b) Organisms of the genus *Klebsiella* have significantly higher nucleotide diversity in infants born via C-section than those delivered vaginally. c) Among organisms present in multiple time-points during antibiotic administration, nucleotide diversity tended to decrease upon administration of antibiotics and increase following cessation of antibiotics. d) Bacterial organisms acquired later in life tended to have lower nucleotide diversity than those acquired earlier in life. e) Bacterial organisms that colonized greater number of infants tended to have higher nucleotide diversity.

The mean bacterial nucleotide diversity within infants did not change over the sampling time. However, bacteriophage and plasmid nucleotide diversity decreased following administration of antibiotics, and bacteria, bacteriophage, and plasmid nucleotide diversity increased after the cessation of antibiotics **(Figure 4c)**. Subspecies detected for the first time later in life had significantly lower nucleotide diversity than those detected earlier **(Figure 4d;** *P* = 0.03), potentially reflecting increasing selective pressures with increasing gut microbiome complexity. Bacterial subspecies that were detected in more infants generally had higher nucleotide diversity **(Figure 4e)**.

Finally, we performed a statistical test to identify genes with significantly different microdiversity than other genes in the genome **(Table 5)**. Genes with significantly lower microdiversity include house-keeping genes like ribosomal protein S16 in bacteria and ParB in bacteriophage (where it is used to maintain circular lysogens ^25^), as well as genes with more interesting functions including a plasmid-encoded polymyxin resistance protein, which is predicted to confer resistance to polymyxin antibiotics ^26^, and bacteriophage lambda head decoration protein D, which stabilizes the expansion of the capsid after genome packaging ^27^. Among the genes with significantly higher microdiversity than the average gene are a bacterial-encoded gene with an immunoglobulin (Ig) domain (which can be involved in cell adhesion and invasion ^28^) and a bacteriophage gene encoding tail fibers (which are often involved in host cell recognition ^29^). Interestingly, both the Ig domain protein and tail fiber protein are involved in host interaction.

**Table 5.** Genes with significantly higher or lower microdiversity than the rest of the genome.

| Type | Taxonomy | Gene ID | q-value | pFam | Description |
| --- | --- | --- | --- | --- | --- |
| **Low microdiversity** |  |  |  |  |  |
| **bacteria** | *E. faecalis* | 23754 | 7.93E-20 | PF00886.18 | Ribosomal protein S16 |
|  | *S. epidermidis* | 16419 | 9.51E-15 | PF02597.19 | ThiS family |
|  | *K. pneumoniae* | 15325 | 4.33E-10 | PF02617.16 | ATP-dependent Clp protease adaptor protein ClpS |
| **phage** | *Escherichia* | 223 | 9.39E-06 | PF08775.9 | ParB family |
|  | *E. coli* | 205 | 0.00011027 | PF02924.13 | Bacteriophage lambda head decoration protein D |
|  | *Phietavirus* | 334 | 0.00018497 | PF00692.18 | dUTPase |
| **plasmid** | *Bacilli* | 0 | 0.00013818 | PF02388.15 | FemAB family |
|  | *K. aerogenes* | 281 | 0.00034062 | PF11183.7 | Polymyxin resistance protein PmrD |
|  | *S. epidermidis* | 290 | 0.00043106 | PF01479.24 | S4 domain |
| **High microdiversity** |  |  |  |  |  |
| **bacteria** | *S. epidermidis* | 15627 | 6.23E-43 | PF05345.11 | Putative Ig domain |
|  | *K.pneumoniae* | 15332 | 2.38E-34 | PF00465.18 | Iron-containing alcohol dehydrogenase |
|  | *E. faecalis* | 23792 | 4.03E-32 | PF13731.5 | WxL domain surface cell wall-binding |
| **phage** | *Escherichia* | 226 | 1.25E-13 | PF03400.12 | IS1 transposase |
|  | *E. coli* | 243 | 6.41E-12 | PF03406.12 | Phage tail fiber repeat |
|  | *E. faecalis* | 293 | 1.18E-08 | PF01183.19 | Glycosyl hydrolases family 25 |
| **plasmid** | Unknown | 358 | 1.95E-28 | PF00665.25 | Integrase core domain |
|  | *Bacilli* | 11 | 4.84E-25 | PF03432.13 | Relaxase/Mobilisation nuclease domain |
|  | *Clostridium* | 374 | 9.71E-14 | PF02782.15 | FGGY family of carbohydrate kinases, C-terminal domain |

### Tracking specific genetic variants within and between populations

To investigate the relationship between the diversity of a population within a single infant (intra-infant diversity) and the diversity of populations of the same subspecies in multiple different infants (inter-infant diversity), we performed a detailed analysis of an *Enterococcus faecalis* bacteriophage (subspecies 482_10.ph) that was present at high coverage depth (>20x) and breadth of coverage (>80%) in 44 infants in our cohort (**Supplemental Table S3**). Genes with a substantial number of intra-infant SNPs had correspondingly more fixed inter-infant substitutions (**Figure 6b)**. We observed that 72% of sites with inter-infant fixed substitutions were also found as intra-infant polymorphisms, indicating that a large fraction of the intra-infant polymorphic variation observed within an infant could be ascribed to mixing of variants that are found alone in other individuals (**Figure 6d**). 34% of intra-infant SNPs were also found to be polymorphic in at least 3 different infants, indicating a substantial overlap in polymorphic variation across infants **(Supplemental Table S4)**.

**Figure 6.**
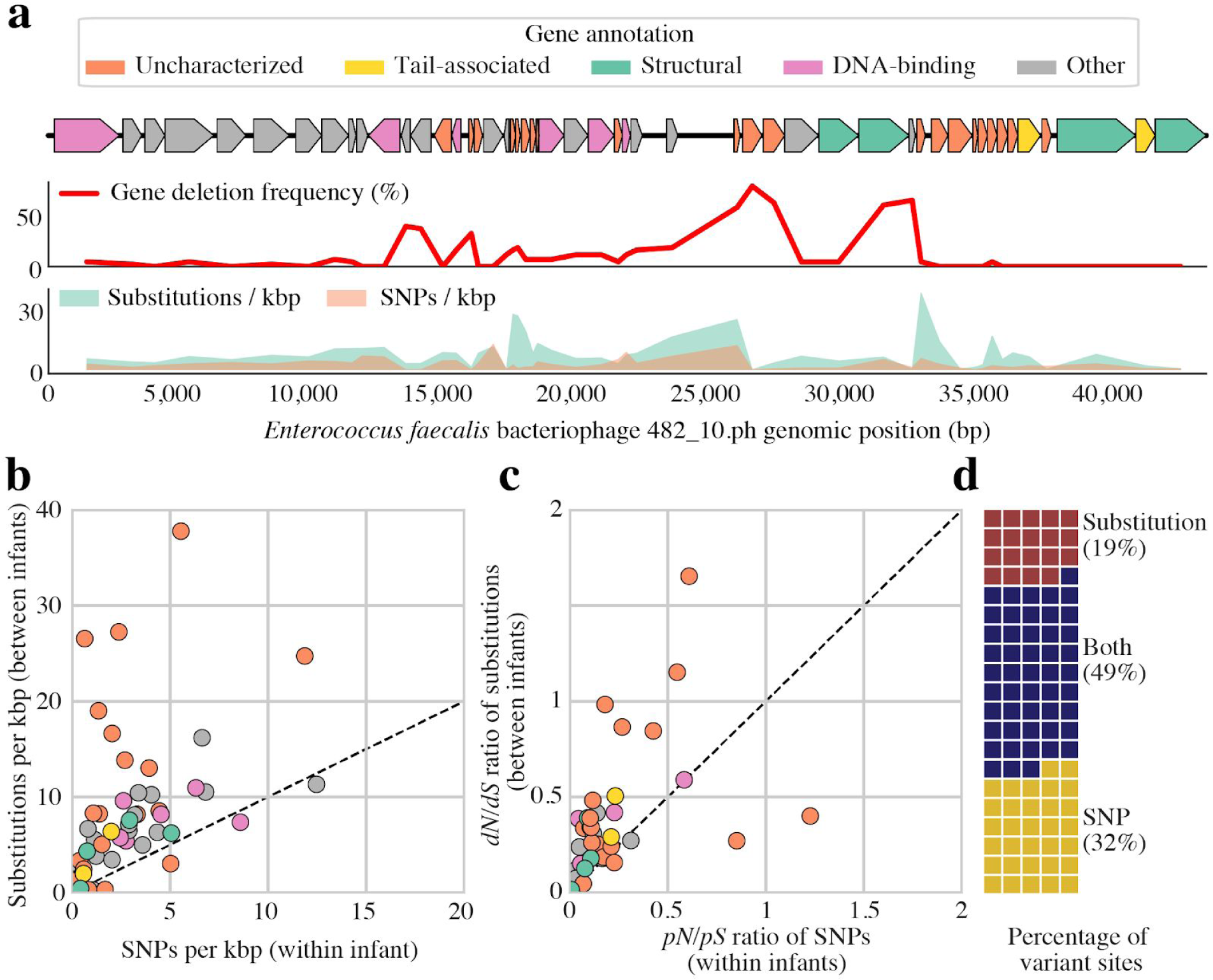
Tracking specific genetic differences within and between populations of an *E. faecalis* bacteriophage. a) Frequencies of gene deletions, substitutions, and SNPs for all genes across an *E. faecalis* bacteriophage genome identified in 44 infants. Genes are colored based on their annotations. b) Frequency of observed substitutions (fixed differences between pairs of infants) in each gene versus frequency of SNPs (positions with multiple alleles in an individual infant). c) Ratios of non-synonymous to synonymous substitutions (*dN/dS*) and ratios of non-synonymous to synonymous population-level variants (*pN/pS*) for each gene. d) Classification of variant sites observed across infants only as substitutions, only as SNPs, and as both.

Seven of the fifty-one genes annotated on the *E. faecalis* bacteriophage genome had *dN/dS* ratios over 0.5, including five proteins of unknown function, a DnaB replication initiation homolog, and a predicted distal tail gene (**Figure 6ab**). The predicted distal tail gene, which might play a role in host specificity, was also found to have a relatively low intra-infant *pN/pS* ratio, possibly indicating selection for variation between but not within individual populations. Multiple small hypothetical proteins also had high *dN/dS* ratios, one of which was only present in ∼50% of infants (**Figure 6c)**. The relaxed purifying selection indicated by high *dN/dS* ratios and the variable presence of these genes perhaps indicates an accessory or vestigial function.

## Discussion

InStrain is an integrated and versatile program for profiling the microdiversity of organisms from metagenomic data. Its ability to perform microdiversity-aware genomic comparisons offers several advantages over existing pipelines, including the consideration of major and minor alleles, thus accounting for the presence of coexisting strains. Because it uses sample-assembled genomes and full paired-read information there is greatly increased confidence that reads are aligned correctly, which improves the high resolution comparisons being made based on entire genomes. Many of these capabilities have been successfully implemented individually in previous studies ^15, 19, 30–33^. However, their simultaneous integration into a well-documented and easy to use pipeline allows substantially more rigorous detection of near-identical strains than the existing commonly used pipelines (**Figure 2**) used in recent high-profile publications to quantify the ecologically critical process of microbiome transmission ^14, 34^. The method substantially increases the stringency of evidence for strain sharing and thus identification of the factors that determine the extent to which this occurs.

Twin studies have previously been used to elucidate relationships between host genetics and human microbiome composition, with the basic premise being that because twins are reared together and share similar environments, increased microbiome similarity between MZ twins compared to DZ twins can be ascribed to genetic effects ^35^. Although studies of adult twins have consistently found some microbial taxa to be more commonly identified in MZ than DZ twins ^36–39^, diet and lifestyle preferences have also been shown to be more similar in MZ twins than DZ twins ^40–42^, presenting significant potential for confounding effects. In contrast to prior studies, all subjects in the current study were housed in the same NICU for the entirety of sampling time. Our findings, based on new and demonstrably more robust methods, indicate that MZ twins shared no more strains of bacteria, bacteriophage, or plasmids than DZ twins. This points to a minimal role of human genetics in early life strain colonization.

Initial colonists are believed to have an outsized role in microbiome development ^43, 44^. The hospitalized premature infants in this study were all given prophylactic antibiotics immediately after birth, housed in isolettes that maintained separation from other infants, and ∼75% were born by cesarean section. These factors likely limited their exposure to microbes from the mother, other family members, and the external home environment. The patterns of strain-sharing among infants in this study suggest the importance of *i) Family-specific sources*. Strains present in two and only two infants were significantly more likely to be shared between siblings **(Figure 3)**, highlighting the role of strain sources such as shared visitors and/or parents in infant colonization. *ii) The hospital environment*. Non-sibling infants born at similar times chronologically shared more strains than those born further apart, indicating that the local hospital microbiome plays a role in strain acquisition. The identification of strains of ESKAPE pathogens (known for their antibiotic resistance and ability to cause nosocomial infections) colonizing large numbers of infants further points to the hospital room as an important source of initial strains. These highly-colonizing strains may have been dispersed in part by healthcare workers that interact with many infants. *iii) Infant physiology*. Infants with similar physiological properties such as gestational age and birth weight shared significantly more strains, potentially due to differences in the development of the human immune system or physical gut environment.. *iv) Unique sources*. The majority of strains identified were found in only a single infant, demonstrating that even in a highly-cleaned environment like the NICU, initial microbiota acquisition is a largely individualized process.

It is difficult to distinguish microbiome diversity that is evolved *in situ* from that introduced by immigration ^1, 10^. In this study of newborn infants we found evidence that initial bacterial microdiversity can be related to mode of acquisition; *Klebsiella* had higher levels of nucleotide diversity in infants born via C-section than those born vaginally, suggesting that there is a more diverse pool of *Klebsiella* strains in the operating room (where *Enterobacteriaceae* have previously been identified ^45^) than in the maternal microbiome. *Firmicutes* may have had lower levels of nucleotide diversity than *Proteobacteria* overall due to their propensity for spore-based transmission ^46^. Patterns of nucleotide diversity in relation to antibiotic administration are consistent with increasing and decreasing selective pressures during and following antibiotic administration, respectively **(Figure 4c)**. The general increase in nucleotide diversity and *dN/dS* ratios of genes involved in cell-cell interactions compared to other functions indicates that these genes are likely under diversifying selection. Identification of house-keeping genes with lower than average nucleotide diversity demonstrates the utility of inStrain for identifying genes under purifying selection **(Table 5)**. The finding that the bacterial strains that colonize the most infants also have the highest nucleotide diversity in infants **(Figure 4e)** is indicative of a relationship between the diversity of the inoculation source (especially the NICU) and the ability of an organism to widely colonize newborns.

By reporting and classifying all gene variants, inStrain enables locus-specific analyses of the genetic differences within and between populations. Further, as inStrain also does not rely upon reference databases or conserved bacterial marker genes, it is capable of tracking genetic variation in bacteriophages and plasmids. For example, applying inStrain to a highly prevalent *E. faecalis* bacteriophage confirmed a relationship between the diversity within individual infants and the subspecies diversity overall, and identified specific genes with divergent *dN/dS* ratios and variable presence **(Figure 6)**. Specifically, we found evidence that nonsynonymous changes in a tail fiber gene are purged within infants (possibly to maintain infectivity), yet selected for between infants (suggestive of variation in bacterial host immunity).

Diversity is a hallmark of stable and healthy human microbiomes ^47–49^. While microbial diversity is typically measured by quantifying the number and evenness of microbial species or genera present in a sample, the detected microbial taxa represent larger populations of cells with within-population genetic heterogeneity. Microdiversity may increase the likelihood of harboring a fit genotype as conditions change. Alternatively, an overall wider gene variant pool may reflect adaptation to spatial variation in local environmental conditions. InStrain allows scientists to easily measure and analyze population microdiversity. In existing and future metagenomic sequencing-based projects, there is the potential to improve our understanding of relationships between microbial population diversity and resilience, stability, population-level phenotypes and to track ecologically relevant processes such as strain migration and *in situ* evolution.

## Methods

### Benchmarking inStrain against other methods

Synthetic comparisons (**Figure 2b**) were performed by using SNP Mutator ^50^ to introduce a known number of mutations into a reference genome (*Escherichia coli* strain SQ88; RefSeq accession number GCF_000988385.1) and comparing the mutated genomes to the original reference genome. For dRep, mutated genomes were compared to the reference genome using dRep on default settings. For inStrain and StrainPhlAn, Illumina reads were simulated for all genomes at 20x coverage using pIRS ^51^. For inStrain, synthetic reads were mapped back to the reference genome using Bowtie 2 ^23^, profiled using “inStrain profile” under default settings, and compared using “inStrain compare” under default settings. For StrainPhlan, synthetic reads profiled with Metaphlan2 ^21^, resulting marker genes were aligned using StrainPhlan, and the ANI of resulting nucleotide alignments was calculated using the class “Bio.Phylo.TreeConstruction.DistanceCalculator(’identity’)” from the BioPython python package^52^. Raw values from this analysis are available in **Supplemental Table S1**.

Isolate-based comparisons (**Figure 2c**) were performed based on the ZymoBIOMICS Microbial Community Standards product (Catalog #D6300). Three samples were prepared from aliquots of this mixture of cells in which DNA extraction, library preparation, and *in silico* sequence trimming and analysis were performed separately. For dRep, reads from each sample were assembled independently using IDBA-UD ^53^, binned into genomes based off of alignment to the provided reference genomes (https://s3.amazonaws.com/zymo-files/BioPool/ZymoBIOMICS.STD.refseq.v2.zip) using nucmer ^54^, and compared using dRep on default settings. For StrainPhlAn, reads from Zymo samples profiled with Metaphlan2, resulting marker genes were aligned using StrainPhlan, and the ANI of resulting nucleotide alignments was calculated as described above. For inStrain, reads from Zymo samples were aligned to the provided reference genomes using Bowtie 2, profiled using “inStrain profile” under default settings, and compared using “inStrain compare” under default settings. “popANI” values were used for inStrain. Eukaryotic genomes were excluded from this analysis, and raw values are available in **Supplemental Table S1**.

Twin-based comparisons **(Figure 2d)** were performed on three randomly chosen sets of twins that were sequenced during a previous study ^24^. For StrainPhlAn, all reads sequenced from each infant were concatenated and profiled using Metaphlan2, compared using StrainPhlAn, and the ANI of resulting nucleotide alignments was calculated as described above. For dRep, all de-replicated bacterial genomes assembled and binned from each infant (available from ^24^) were compared in a pairwise manner using dRep under default settings. For inStain, strain-sharing from these six infants was determined using the methods described below. ANI values from all compared genomes and the number of genomes shared at a number of ANI thresholds are available for all three methods in **Supplemental Table S1**.

### Calling, detection, and profiling of sub-species of bacteria, bacteriophage, and plasmids

Genomes of bacteria, bacteriophage, plasmid, and eukaryotes were previously binned from the infants comprising this study, as described previously ^24^ (genomes are available for download at https://doi.org/10.6084/m9.figshare.c.4740080.v1). To generate a single genome set, all bacterial genomes were compared to each other using dRep version 2.2.0 under default settings, all bacteriophage genomes were compared to each other using the command “dRep dereplicate --S_algorithm ANImf -nc .5 -l 10000 -N50W 0 -sizeW 1 --noQualityFiltering --clusterAlg single”, and all plasmid genomes were compared to each other using the same command as bacteriophages. Genomes with ANI >= 98% were classified as the same subspecies, and the genome with the highest score (as determined by dRep) was chosen as the representative genome from each subspecies. Bacteriophage and plasmid genomes with taxonomic classifications specifying “Eukarya” were marked as “likely human” and excluded from further analysis. Information about sub-species is available in **Supplemental Table S3**.

Reads from each individual fecal sample, reads from each infant concatenated together (referred to as “coReads”), and reads from all negative extraction control samples concatenated together were mapped to all representative sub-species genomes concatenated together using Bowtie 2 with default settings. “InStrain profile” was run on all resulting mapping files with default settings. Detection of a sub-species in a sample was defined as that genome being present with >= 0.5 unmaskedBreadth (meaning that at least half of the bases in the genome were covered by at least 5 reads). Mappings from coReads were used for all analyses unless otherwise specified. Subspecies detected in the negative extraction control sample, and genomes detected significantly more often in one of the six individual sampling campaigns were marked “likely contaminant” and excluded from further analysis. Information on sub-species abundance is available in **Supplemental Table S3**.

### Identification of strains and associations with metadata

Strain-level comparisons were performed between subspecies detected in multiple samples from the same infant over time-series sampling, and strain-level comparisons were performed between subspecies detected in the coReads of multiple infants. For within-infant subspecies comparisons, all subspecies detected in multiple individual samples from an infant (as described above) were compared using “inStrain compare”. Raw values are available in **Supplemental Table S2**. For between-infant subspecies comparisons, subspecies that were detected in coRead samples from multiple infants (or the coRead sample consisting of all negative extraction controls) were compared using “inStrain compare” with default settings. A distance matrix then created for each subspecies based on popANI values, and this matrix was used to cluster subspecies into a number of individual strains using ‘average’ hierarchical clustering with a threshold of 99.999% ANI with the scipy cluster package ^55^. Strains that were present in the reads from the negative extraction control, and strains from subspecies that were filtered out using the methods described above were removed from further analysis. Raw comparison values and strain identities are available in **Supplemental Table S2**.

The number of strains shared between infants was visualized in **Figure 3ab** using Circos ^56^. The strain-level Jaccard distance between infants was calculated according to the formula: Jaccard similarity = number of strains shared by both infants / number of strains present in either infant. P-values for Jaccard similarity are based on the Wilcoxon rank-sum statistic between all twin pairs and all non-twin pairs, as calculated using the python module scipy.stats.ranksums ^55^. Associations between the number of strains shared between infants and their difference in birth day, birth weight, and gestational age was determined by first binning the metadata variable into windows of size 20 (birth weight, gestational age) or 1 (gestational age) and calculating the average number of strains shared between infants within that window. Siblings were excluded from this analysis. P-values and R^2^ values are based on linear regression, as calculated using the python module sklearn.

The visualization in **Figure 3g** was created by first identifying the eight bacterial species with the highest colonizing strain, and then assigning a specific color to each strain within these eight species that colonized at least five infants. For each value on the x-axis, the y-axis displays the proportional count of the total strains detected in infants by strains that colonized at least that value of infants.

### Nucleotide diversity analysis

The coReads inStrain analysis described above resulted in a total of 8,336 subspecies / infant pairs in which the subspecies genome was detected at 5x coverage across at least 50% of the genome. The Wilcoxon rank-sum statistic (as implemented in Scipy ^55^) was used to compare the nucleotide diversity of different sets of genomes and generate p-values.

Time-series sampling information from individual infants was used to analyze nucleotide diversity in relation to antibiotic administration **(Figure 4c)**. Using the same definition of presence as described above, subspecies were identified that were present in at least two of the following three windows: seven days prior to antibiotic administration, during antibiotic administration, seven days after antibiotic administration. To determine whether antibiotic administration changed before as compared to during antibiotic administration, for each organism type (bacteria, plasmid, or bacteriophage), the nucleotide diversities of all subspecies of that type present in both widows were subjected to a two sided dependent t-test (as implemented using the Scipy module “scipy.stats.ttest_rel”). The same procedure was used to test for significant differences between “during” and “after” antibiotic administration, and “before” and “after” antibiotic administration. P-values were corrected for multiple testing using Benjamini-Hochberg p-value correction.

Samples for individual fecal samples were also used to test for differences between subspecies microdiversity and date of subspecies acquisition **(Figure 4d)**. Samples within five days of antibiotic administration were excluded from this analysis, the first day that a bacterial subspecies was detected as present was plotted against the nucleotide diversity of the subspecies in that sample, and a linear regression line of best fit was plotted for infants deriving from each sampling campaign. A binomial t-test (as implemented using the Scipy module scipy.stats.binom_test) was used to determine the p-value for 6/6 campaigns displaying a negative slope.

### Gene-based nucleotide analysis

InStrain was used on default settings to profile genes for all detected subspecies in individual samples and coReads, using gene annotations provided by Prodigal ^57^ run in metagenome mode on original assemblies. Genes with significantly different coverage and/or nucleotide diversity than the rest of genes on the genome were identified using data from coReads profiling of subspecies. For each genome present in at least three infants, the coverage / nucleotide diversity of each gene on the genome across all infants in which the subspecies was present were compared to the coverage / nucleotide diversity of all other genes on the genome across all infants in which the subspecies was present using the Wilcoxon rank-sum statistic (as implemented in Scipy). P-values were corrected to q-values to account for multiple hypothesis testing using Benjamini-Hochberg p-value correction. Genes were annotated based on pFam database HMMs ^58^. For display in **Table 5**, only genes with pFam annotations that did not include the words “Uncharacterized” or “unknown” in the description were retained, all genes with significant differences in coverage (in addition to nucleotide diversity) were excluded, and a maximum of one gene from each taxonomic annotation was allowed for inclusion in each quadrant of high/low microdiversity and organism type.

### Tracking specific nucleotide variants

*Enterococcus faecalis* bacteriophage subspecies 482_10.ph was identified with at least 80% breadth of coverage and 20x coverage depth in the coReads of 44 infants. Open reading frames were called using Prodigal in metagenome mode, and genes were annotated using USEARCH to search against the UniRef100 database. Gene categories (tail-associated, structural, etc.) were assigned based on manual inspection of the resulting database hits. The gene map presented in **Figure 6a** was generated using the python module “dna_features_viewer”.

Bi-allelic SNPs (intra-infant variants) were identified based on the results of “inStrain profile_genes”, where the resulting “SNP_mutation_types” table was subset to SNVs with an allele_count of 2. Substitutions (inter-infant variants) were identified from the “SNVs” table resulting from the operation “inStrain profile”, where the table was subset to SNVs with an allele_count of 1. The number of genomic locations where an SNV was identified in at least one infant, where a substitution was identified in at least one infant, and where both were identified in at least one infant was displayed in a waffle plot using the python module “PyWaffle”.

Synonymous and nonsynonymous variants were identified using inStrain, and the total number of synonymous and nonsynonymous sites in each gene was determined using methods from the script “dnds_from_drep.py” ^59^. *dN/dS* was calculated using the formula [(non-synonymous substitutions / non-synonymous sites) / (synonymous substitutions / synonymous sites)], and *pN/pS* was calculated using the formula [(non-synonymous SNPs / non-synonymous sites) / (synonymous SNPs / synonymous sites)]. The number of substitutions per kbp and the number of SNPs per kbp were calculated by dividing the total number of substitutions / SNPs identified in each gene in all infants by the sum of the length of the gene times the masked breadth (the percentage of the gene with at least 5x coverage; the coverage required to call a SNV) of the gene for each infant the gene was identified in. Genes with a masked breadth ≥ 50% were defined as being present, and the gene deletion frequency was calculated as the percentage of infants where the gene was not present.

## Supplemental Information

Supplemental Figure S1. Graphical comparison of inStrain to other strain-level analysis pipelines.

Supplemental Figure S2. Genome de-replication at 98% ANI reduces multi-mapped reads. **a)** Reads from sample N1_004_0008G1 were mapped to the set of genomes exclusively assembled from infant N1_004 (primary mapping), and reads that mapped to genome dasN1_004_010G1_maxbin2.maxbin.002 in the primary mapping were next mapped to set of genomes consisting of all genomes assembled from all infants de-replicated at 99.8% ANI (secondary mapping). The number of reads mapping to genome dasN1_004_010G1_maxbin2.maxbin.002 is the secondary mapping is shown with an orange dot, and the number of reads mapping to other genomes in the secondary mapping are shown with blue dots. The probability of occurrence of an identical 190bp stretch (the length of read pairs in sample N1_004_0008G1) given an overall genome ANI is drawn with a dotted line using the formula: Probability of identical 190bp fragment = (genome ANI) ^ 190. The fit between the data and the line indicate that read pairs fail to map to the original genome in the secondary mapping due to the presence of identical regions in alternate genomes. **b)** The same process was performed as described in (**a**), but the secondary mapping was filtered to remove all read pairs with MapQ scores less than 2, removing read pairs that map equally well to two locations. In this context, only a miniscule number of reads map to alternate genomes in the secondary mapping, but the number of reads mapping to the original genome is reduced substantially compared to **(a)**. For more information see the online inStrain documentation.

Supplemental Figure S3. Genomic comparisons between subspecies present in the same infant over time. In cases where the same subspecies was detected in multiple time-points over the time-series sampling of an infant, the percentage of comparisons between these subspecies that exceed various popANI **(a)** and conANI **(b)** thresholds is plotted. The use of popANI allows greater stringency than conANI.

Supplemental Figure S4. Metadata associated with strain sharing and nucleotide diversity. **a**) Associations between strain sharing and birth weight, gestational age, and birth study day for bacteria, bacteriophages, and plasmids. **b**) Infants of the same campaign are not more likely to share strains.

Supplemental Figure S5. Metadata associated with nucleotide diversity. **a**) Nucleotide diversity is associated with library preparation methodology and sequencing machine used. **b**) Nucleotide diversity of plasmids, bacteriophages, and bacteria in the six sampling campaigns. Cases where one class has a nucleotide diversity significantly different from the other two within a sampling campaigned are annotated (*P <* 0.05; Wilcoxon rank-sums). **c**) The association between *Klebsiella* and birth mode remains significant when not considering samples from the Sloan2 and NIH4 campaigns.

Supplemental Table S1. Information related to inStrain benchmarking.

Supplemental Table S2. Strain-level comparisons within infant samples, between infant coReads, and strain identities.

Supplemental Table S3. Abundance of subspecies in all infants, individual samples, and controls, and information about subspecies genomes and representatives.

Supplemental Table S4. Detailed SNP information for *Enterococcus faecalis* bacteriophage subspecies 482_10.ph.

**Supplemental Document S1.** InStrain program documentation.

## Acknowledgements

This research was supported by the National Institutes of Health (NIH) under award RAI092531A, the Alfred P. Sloan Foundation under grant APSF-2012-10-05, a National Science Foundation Graduate Research Fellowship to M.O. under Grant No. DGE 1106400, and Chan Zuckerberg Biohub.

