## Supplemental Figure S1 for "InStrain enables population genomic analysis from metagenomic data and rigorous detection of identical microbial strains"

| | | | Graphical example of output for different sample population structures | | |
|---|---|---|---|---|---|
| | | | 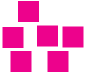 | 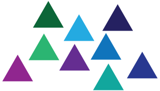 | 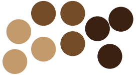 |
| Method | Description | Programs | Clonal | Heterogenous | Distinct |
| Consensus SNP calling | The consenus base at each position of the genome defines the strain in that sample. One possible strain detection per sample. | StrainPhlan<br>MIDAS | 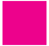 | 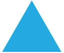 | 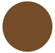 |
| Haplotype phasing | Strains are defined as sets of polymorphic bases that co-vary in frequency accross samples. Each strain is a disntinct genotype. Many possible strain detections per sample. | ConStrain<br>DESMAN | 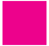 | 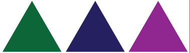 | 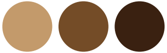 |
| Characterizing the strain cloud | Distict strains are not defined. Polymophic base frequencies, microdiversity, and variations in coverage are calculated for each position along the genome. | inStrain | 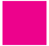 | 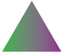 | 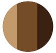 |
