## Supplementary figures and images for "InStrain enables population genomic analysis from metagenomic data and rigorous detection of identical microbial strains"

### Supplemental Figure S2

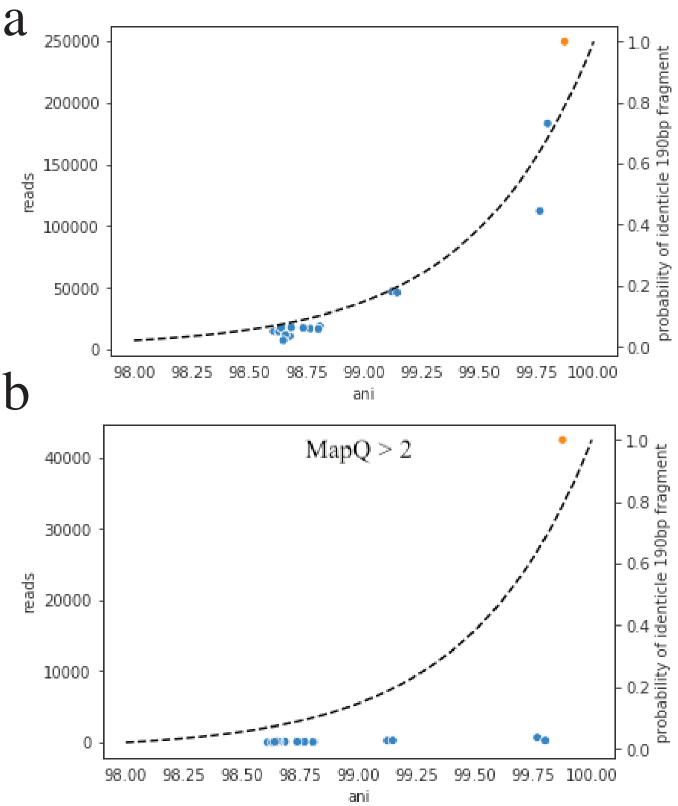

### Supplemental Figure S3

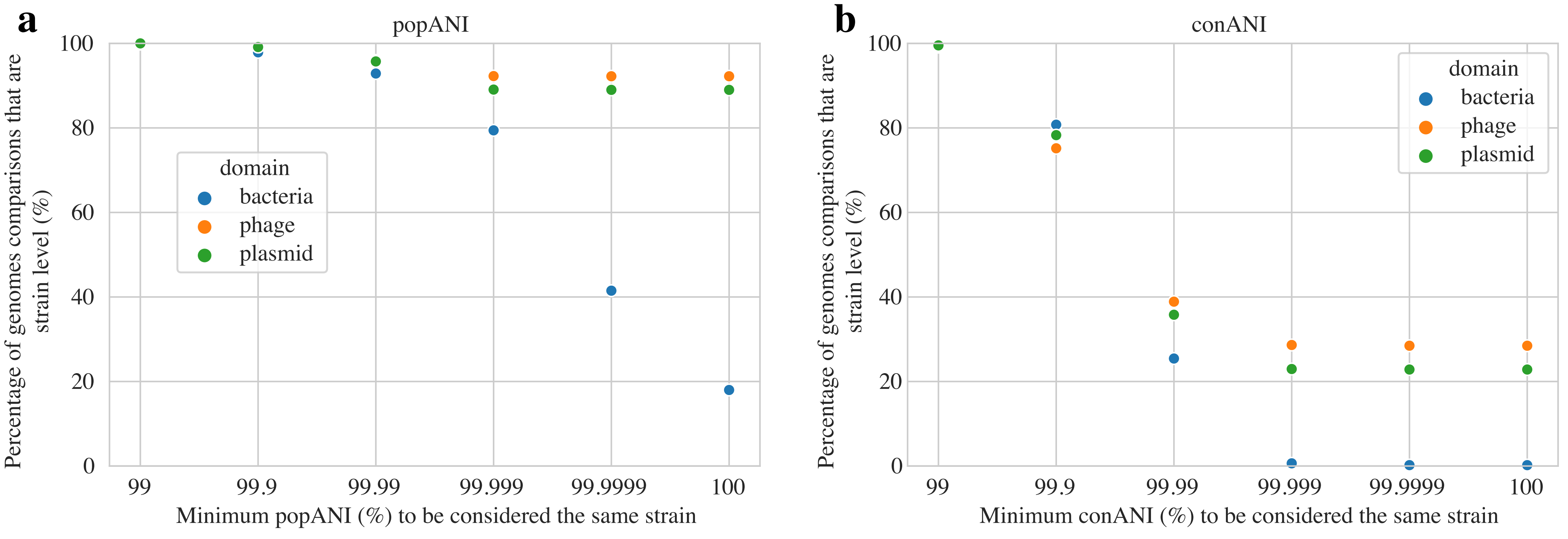

### Supplemental Figure S4

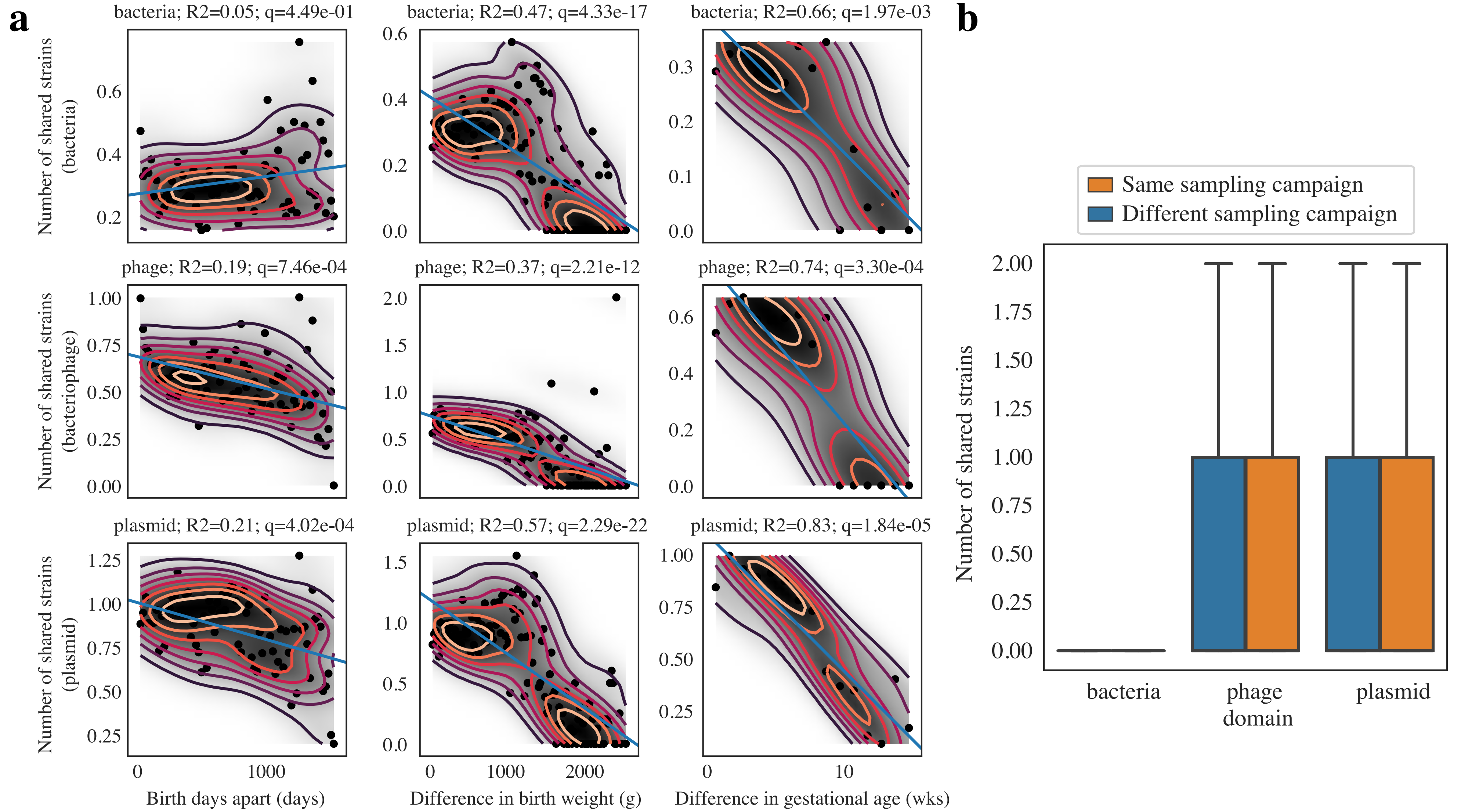

### Supplemental Figure S5

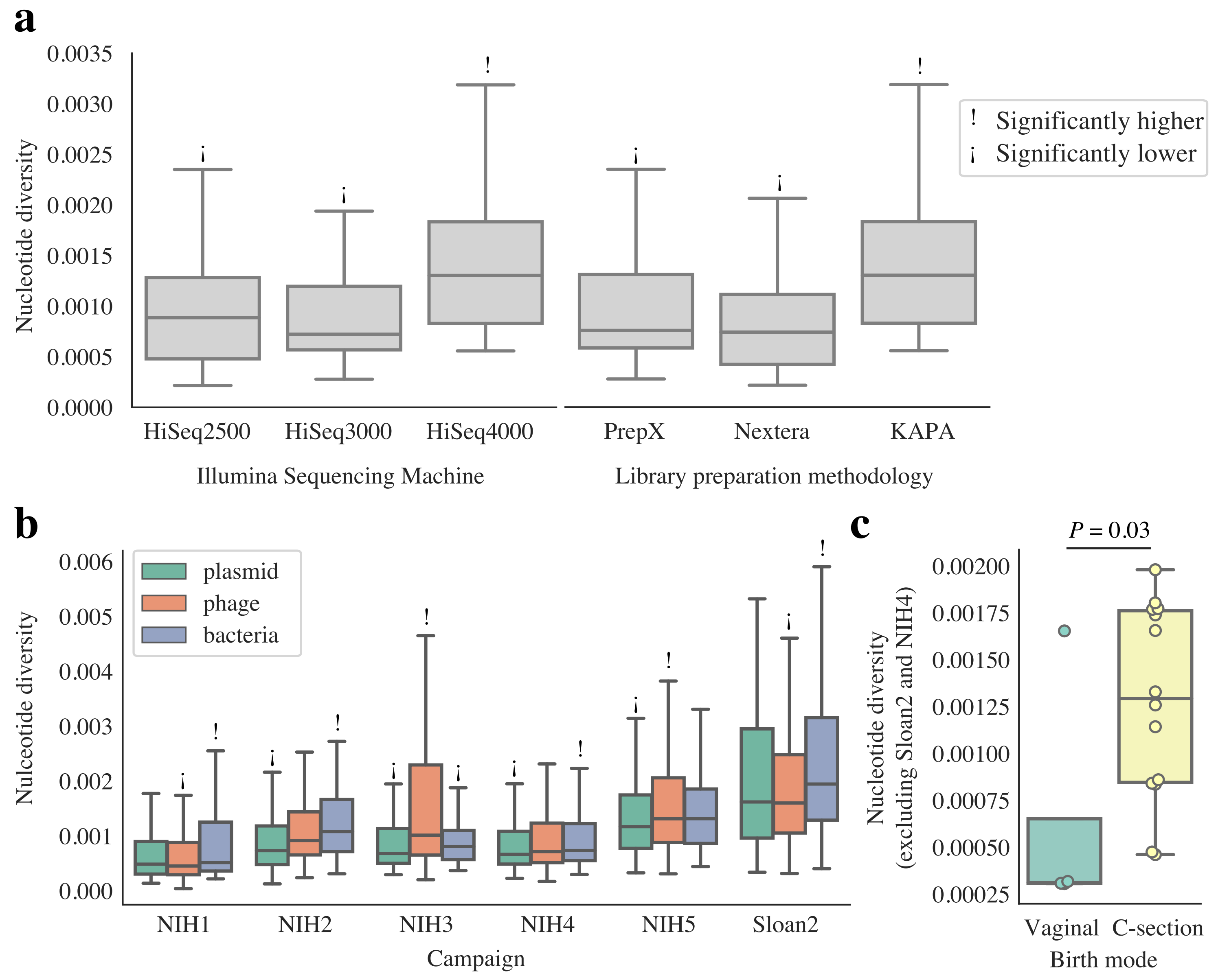
