## Supplemental Document S1 for "InStrain enables population genomic analysis from metagenomic data and rigorous detection of identical microbial strains"

### inStrain

***Release 1.0.0***

**Jan 21, 2020**

InStrain is a tool for analysis of co-occurring genome populations from metagenomes that allows highly accurate genome comparisons, analysis of coverage, microdiversity, and linkage, and sensitive SNP detection with gene localization and synonymous non-synonymous identification

Source code is available on GitHub.

See links to the left for *Installation and Quickstart* instructions

Comments and suggestions can be sent to Matt Olm and/or Alex Crits-Christoph

Bugs reports and feature requests can be submitted through GitHub.

CHAPTER 1

---

Contents

---

### 1.1 Installation and Quickstart

#### 1.1.1 Installation

##### Using pip

inStrain is written in python. To install inStrain using the PyPi python repository, simply run

```
$ pip install instrain
```

Or to install from GitHub run

```
$ git clone https://github.com/MrOlm/instrain.git

$ cd instrain

$ pip install .
```

That's it!

Pip is a great package with many options to change the installation parameters in various ways. For details, see pip documentation

##### Dependencies

inStrain requires a few other programs to run. Not all dependencies are needed for all operations. There are a number of python package dependencies, but those should install automatically when inStrain is installed using pip

**Essential**

- samtools This is needed for pysam

**Optional**

- coverM This is needed for the quick_profile operation
- Prodigal This is needed to profile on a gene by gene level

##### 1.1.2 Quick Start

The functionality of inStrain is broken up into several core modules. For more details on these modules, see module_descriptions.:

```
$ inStrain -h

              ...::: inStrain v1.0.0 :::...

Matt Olm and Alex Crits-Christoph. MIT License. Banfield Lab, UC Berkeley. 2019

Choose one of the operations below for more detailed help. See https://instrain.
↪readthedocs.io for documentation.
Example: inStrain profile -h

  profile         -> Create an inStrain profile (microdiversity analysis) from a␣
↪mapping.
  compare         -> Compare multiple inStrain profiles (popANI, coverage_overlap,␣
↪etc.)
  profile_genes   -> Calculate gene-level metrics on an inStrain profile
  genome_wide     -> Calculate genome-level metrics on an inStrain profile
  quick_profile   -> Quickly calculate coverage and breadth of a mapping using coverM
  filter_reads    -> Commands related to filtering reads from .bam files
  plot            -> Make figures from the results of "profile" or "compare"
  other           -> Other miscellaneous operations
```

Below is a list of brief descriptions of each of the modules. For more information see module_descriptions, for help understanding the output, see *Example output and explanations*, and to change the parameters see choosing_parameters

**See also:**

**module_descriptions** for more information on the modules

***Example output and explanations*** to view example output

**choosing_parameters** for guidance changing parameters

**preparing_input** for information on how to prepare data for inStrain

###### profile

inStrain profile is the main method of the program. It takes a *.fasta* file and a *.bam* file (consisting of reads mapping to the *.fasta* file) and runs a series of steps to characterize the microdiversity, SNPs, linkage, etc. Details on how to generate the mapping, how the profiling is done, explanations of the output, how to choose the parameters can be found at preparing_input and module_descriptions

To run inStrain on a mapping run the following command:

```
$ inStrain profile .bam_file .fasta_file -o IS_output_name
```

##### compare

inStrain is able to compare multiple read mappings to the same .fasta file. Each mapping file must first be make into an inStrain profile using the above command. The coverage overlap and popANI between all pairs is calculated:

```
$ inStrain compare -i IS_output_1 IS_output_2 IS_output_3
```

##### profile_genes

Once you've run *inStrain profile*, you can also calculate gene-wise microdiversity, coverage, and SNP functions using this command. It relies on having gene calls in the *.fna* format from the program prodigal:

```
$ inStrain profile_genes -i IS_output -g called_genes.fna
```

##### genome_wide

This module is able to translate scaffold-level results to genome-level results. If the *.fasta* file you mapped to consists of a single genome, running this module on its own will average the results among all scaffolds. If the *.fasta* file you mapped to consists of several genomes, by providing a *scaffold to bin file* or a list of the individual *.fasta* files making up the combined *.fasta* file, you can get summary results for each individual genome. Running this module is also required before generating plots.

> $ inStrain genome_wide -i IS_output -s genome1.fasta genome2.fasta genome3.fasta

##### quick_profile

This auxiliary module is merely a quick way to calculate the coverage and breadth using the blazingly fast program coverM. This can be useful for quickly figuring out which scaffolds have any coverage, and then generating a list of these scaffolds to profile with inStrain profile, making it run faster:

```
$ inStrain quick_profile -b .bam_file -f .fasta_file -s scaffold_to_bin_file -o␣
↪output_name
```

##### filter_reads

This auxiliary module lets you do various tasks to filter and/or characterize a mapping file, and then generate a new mapping file with those filters applied:

```
$ inStrain filter_reads .bam_file .fasta_file -g new_sam_file_location
```

##### plot

This method makes a number of plots from an inStrain object. It is required that you run *genome_wide* first before running this module:

```
$ inStrain plot -i IS_output
```

##### other

This module lets you do random small things, like convert IS_profile objects that are in an old format to the newest format.

#### 1.2 Overview and FAQ

##### 1.2.1 Overview

When you sequence any microbial genome(s), you sequence a population of cells. This population may be a nearly clonal population grown up from an isolate in a culture flask, or a highly heterogenous population in the real world, but there is always real biological genetic hetereogeneity within that population - every cell does not have the same genotype at every single position.

**InStrain is a program for measuring, comparing, and interrogating the genetic heterogeneity of microbial populations in and between metagenomic samples.** We refer to these intraspecific differences as "microdiversity"

##### 1.2.2 FAQ (Frequently asked questions)

###### How does inStrain compare to other bioinformatics tools for strains analysis?

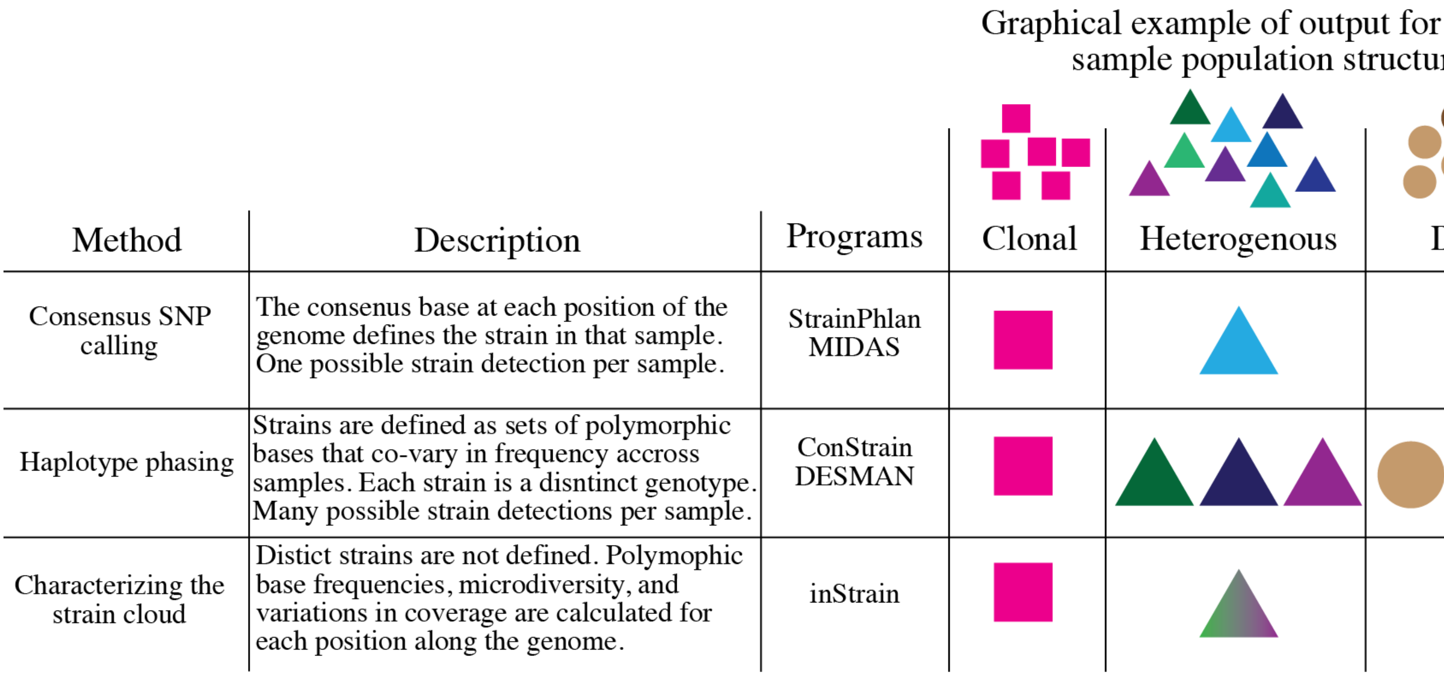

###### What can inStrain do?

inStrain includes calculation of nucleotide diversity, calling SNPs (including non-synonymous and synonymous variants), reporting accurate coverage / breadth, and calculating linkage disequilibrium in the contexts of genomes, contigs, and individual genes.

inStrain also includes comparing the frequencies of fixed and segregating variants between sequenced populations with extremely high accuracy, out-performing other popular strain-resolved metagenomics programs.

The typical use-case is to generate a *.bam* file by mapping metagenomic reads to a bacterial genome that is present in the metagenomic sample, and using inStrain to characterize the microdiversity present.

Another common use-case is detailed strain comparisons that involves comparing the genetic diversity of two populations and calculating the extent to which they overlap. This allows for the calculation of population ANI values for extremely similar genomic populations (>99.999% average nucleotide identity).

**See also:**

*Installation and Quickstart* To get started using the program

**module_descriptions** For descriptions of what the modules can do

*Example output and explanations* To view example output

**preparing_input** For information on how to prepare data for inStrain

**choosing_parameters** For detailed information on how to make sure inStrain is running correctly

##### When should I use inStrain?

inStrain is intended to be used as a genome-resolved metagenomics approach. Genome-resolved metagenomics involves sequencing and de novo assembly of the actual microbial genomes present in the sample(s) of interest. It is these microbial genomes, and not microbial genomes derived from reference databases, that we will then use as scaffolds on which to map reads from the sample.

We don't recommend using reference genomes for strain-resolved inferences in metagenomes. This is because reference databases have usually poorly sampled the true extent of microbial diversity below the species level across many environments. Using even partially inaccurate references can lead to inaccurate conclusions about the genetic variation within your samples.

inStrain can be run on individual microbial genomes assembled and binned from a metagenome, sets of de-replicated microbial genomes, or entire metagenomic assemblies at once.

##### When should I probably not use inStrain?

When you have not assembled genomes from the metagenomic samples you are interrogating; when breadth and coverage of the consensus genome are low; when you wish to compare populations that are <95% ANI with each other; when you are interested in species-level community composition, not intra-population diversity.

##### How does inStrain work?

The reasoning behind inStrain is that every sequencing read is derived from a single DNA molecule (and thus a single cell) in the original population of a given microbial species. During assembly, the consensus of these reads are assembled into contigs and these contigs are binned into genomes - but by returning to assess the variation in the reads that assembled into the contigs, we can characterize the genetic diversity of the population that contributed to the contigs and genomes.

The basic steps:

1. Map reads to a *.fasta* file to create a *.bam* file
2. Stringently filter mapped reads and calculate coverage and breadth
3. Calculate nucleotide diversity and SNPs
4. Calculate SNP linkage
5. Optional: calculate gene statistics and SNP function

6. Optional: compare SNPs between samples.

###### What is unique about the way that inStrain compares strains?

Most strain-resolved pipelines compare the dominant allele at each position. If you have two closely related strains A and B in sample 1, with B being at higher abundance, and two closely related strains A and C in sample 2, with C being at higher abundance, most strain comparison pipelines will in actuality compare strain B and C. This is because they work on the principle of finding the dominant strain in each sample and then comparing the dominant strains. inStrain, on the other hand, is able to identify the fact that A is present in both samples. This is because it doesn't just compare the dominant alleles, but compares all alleles in the two populations. See doc:*module_descriptions* and choosing_parameters for more information.

###### What is a population?

To characterize intra-population genetic diversity, it stands to reason that you first require an adequate definition of "population". inStrain relies mainly on population definitions that are largely technically limited, but also coincide conveniently with possibly biological real microbial population constraints (link1. **'link2<https://www.nature.com/articles/s41467-018-07641-9>'_**.). Often, we dereplicate genomes from an environment at average nucleotide identities (ANI) from 95% to 99%, depending on the hetereogeneity expected within each sample - lower ANIs might be preferred with more complex samples. We then assign reads to each genome's population by stringently requiring that combined read pairs for SNP calling be properly mapped pairs with an similarity to the consensus of at least 95% by default, so that the cell that the read pair came from was at least 95% similar to the average consensus genotype at that position. Within environment, inStrain makes it possible to adjust these parameters as needed and builds plots which can be used to estimate the best cutoffs for each project.

##### 1.2.3 Glossary of terms used in inStrain

**Community** The collection of taxa in a metagenome, i.e. the species diversity of a microbiome.

**Population** The collection of cells for each taxa in a metagenome, i.e. the genetic diversity of each species or subspecies in a microbiome.

---

**Note:** inStrain is for characterizing metagenomes at the population level, not at the community level.

---

**SNP** A SNP is a Single Nucleotide Polymorphism, a genetic variant of a single nucleotide change that some percentage of the cells that comprise a species population. We identify and call SNPs using a simple model to distinguish them from errors, and more importantly in our experience, careful read mapping and filtering of paired reads to be assured that the variants (and the reads that contain them) are truly from the species being profiled, and not from another species in the metagenome (we call it 'mismapping' when this happens). Note that a SNP refers to genetic variation *within a read set*.

**SNV** Single nucleotide variant - in inStrain used interchangeably with SNP

**Microdiversity** We use the term microdiversity to refer to intraspecific genetic variation, i.e. the genetic variation between cells within a microbial species. To measure this, we calculate a per-site nucleotide diversity of all reads - thus this metric is slightly influenced by sequencing error, but within study error rates should be consistent, and this effect is extremely minor compared to the extent of biological variation observed within samples. The metric of nucleotide diversity (often referred to as 'pi' in the population genetics world) is from Nei and Li 1979, calculated per site and then averaged across all sites.

**refSNP** A genetic difference between the consensus of a read set and a reference genome. This is in contrast to SNPs, which are variants within a population being studied - reference SNPs are differences between the population you are studying (your reads) and the genome that you are mapping to. If you are mapping to a genome that was assembled from that sample, there will be very few to no refSNPs, because the consensus of that genome was built from the consensus of the reads in that sample. However, refSNPs are useful to track and understand cross-mapping, and we also use the percentage of refSNPs per read pair to filter read mappings.

**popANI** Calculated by *inStrain compare* function between two different inStrain profiles.

**N SNP** A polymorphic variant that changes the amino acid code of the protein encoded by the gene in which it resides; non-synonymous.

**S SNP** A polymoprhic variant that does not change the amino acid code of the protein encoded by the gene in which it resides; synonymous.

**ANI** Average nucleotide identity. The average nucleotide distance between two genomes or .fasta files. If two genomes have a difference every 100 base-pairs, the ANI would be 99%

**fasta file** A file containing a DNA sequence. Details on this file format can be found on wikipedia

**bam file** A file containing metagenomic reads mapped to a DNA sequence. Very similar to a *.sam* file. Details can be found online

#### 1.3 Program documentation

##### 1.3.1 Preparing input

There are two simple inputs to *inStrain*: a *.fasta* file and a mapping file in *.bam* format. A third, a prodigal *.faa* file, can be used in later steps Here we go over some considerations involved in choosing these inputs.

###### Preparing the .fasta file

A *.fasta* file contains the DNA sequences of the contigs that you map your reads to. Choosing what *.fasta* you will use (consensus / reference genomes) is extremely important and will affect the interpretation of your *inStrain* results. Below we describe the three most common strategies.

Please note that the *.fasta* file must always be the same as, or a subset of, the *.fasta* file used to create the *.bam* file, i.e. the *.fasta* file that reads were mapped to.

###### Using a collection of genomes (recommended)

The recommended workflow for running *inStrain*:

1. Assemble reads into contigs for each sample collected from the environment. Recommended software: IDBA_UD, MEGAHIT, metaSPADES.
2. Bin genomes out of each assembly using differential coverage binning. Recommended software: Bowtie2 (for mapping), MetaBAT, CONCOCT, DasTOOL (for binning).
3. Dereplicate the entire set of genomes across all samples from the environment at 97-99% identity, filter genome set draft quality genomes. Recommended software: dRep, checkM.
4. Create a bowtie2 index of the representative genomes from this dereplicated set and map reads to this set from each sample: Recommended software: Bowtie2
5. Profile the resulting mapping *.bam* files using inStrain.
6. Use *inStrain genome_wide* to calculate genome-level microdiveristy metrics for each originally binned genome.

The most important aspect of this workflow is to **map to many genomes at once**. Mapping to just one genome at a time is highly discouraged, because this encourages mismapped reads from other genomes to be recruited by this genome. By including many (dereplicated) genomes in your bowtie2 index, you will be able to far more accurately filter out mismapped reads and reduce false positive SNPs.

For more information on this, see choosing_parameters

###### Using a single genome .FASTA file

If your .fasta file is a single genome, the main consideration is that it should be a good representitive genome for some organism in your sample. Ideally, it was assembled directly from that sample, isolated from that sample, or you have some other evidence that this genome is highly representation of a species in that sample. Regardless, you should check your *inStrain plot* output and *scaffold_info.tsv* output file to be sure that your inStrain run had decent coverage and breadth of coverage of the genome that you use before attempting to interpret the results.

Remember, your *.fasta* file can be a subset of the *.fasta* file that was used to create the *.bam* file. You can create a BAM with all dereplicated genomes from your environment, but then just pass a *.fasta* file for only the genomes of particular interest. This approach is recommended as opposed to creating a BAM for just each genome, as it reduces mismapping.

###### Using a metagenomic assembly

You can also pass *inStrain* an entire metagenomic assembly from a sample, including either binned or unbinned contigs. In this case, the output inStrain profile will include population information for each contig in the set. To then break it down by microbial genome / species, You can use `inStrain genome_wide` including a scaffold to bin file to generate results by genome.

###### Preparing the .bam file

*inStrain* requires paired-end Illumina read sequencing. We recommend using Bowtie2 to map your reads to your genome.

Bowtie2 default parameters are what we use for mapping, but it may be worth playing around with them to see how different settings perform on your data. It is important to note that the *-X* flag (capital X) is the expected insert length and is by default *500*. In many cases (e.g., 2x250 bp or simply datasets with longer inserts) it may be worthwhile to increase this value up to *-X 1000* for passing to bowtie2.

###### Preparing the prodigal *.fna* genes file for gene-level profiling

You can run prodigal on your *.fasta* file to generate the *.fna* file with the gene-level information that *inStrain profile_genes* requires.

Example:

> $ prodigal -i assembly.fasta -d genes.fna

##### 1.3.2 Module descriptions

The functionality of inStrain is broken up into modules. To see a list of available modules, check the help:

```
$ inStrain -h

             ...::: inStrain v1.0.0 :::...

Matt Olm and Alex Crits-Christoph. MIT License. Banfield Lab, UC Berkeley. 2019

Choose one of the operations below for more detailed help. See https://instrain.
↪readthedocs.io for documentation.
Example: inStrain profile -h

 profile         -> Create an inStrain profile (microdiversity analysis) from a␣
↪mapping.
 compare         -> Compare multiple inStrain profiles (popANI, coverage_overlap, etc.
↪)
 profile_genes   -> Calculate gene-level metrics on an inStrain profile
 genome_wide     -> Calculate genome-level metrics on an inStrain profile
 quick_profile   -> Quickly calculate coverage and breadth of a mapping using coverM
 filter_reads    -> Commands related to filtering reads from .bam files
 plot            -> Make figures from the results of "profile" or "compare"
 other           -> Other miscellaneous operations
```

##### IS_profile

An IS_profile (inStrain profile) is created by running the *inStrain profile* command. It contains all of the program's internal workings, cached data, and output is stored. Additional modules can then be run on an IS_profile (to analyze genes, compare profiles, etc.), and there is lots of nice cached data stored in it that can be accessed using python.

***Example output and explanations*** For help finding where the output from your run is located in the IS_profile

***Advanced use*** For access to the raw internal data (which can be very useful)

##### profile

The most complex part of inStrain, and must be run before any other modules can be. The functionality of *profile* is broken into several steps.

First, all reads in the .bam file are filtered to only keep those that map with sufficient quality. Reads must be paired (all non-paired reads will be filtered out) and an additional set of filters are applied to the read pair (not the individual reads). Command line parameters can be adjusted to change the specifics, but in general:

- Pairs must be mapped in the proper orientation with an expected insert size. The minimum insert distance can be set with a command line parameter. The maximum insert distance is a multiple of the median insert distance. So if pairs have a median insert size of 500bp, by default all pairs with insert sizes over 1500bp will be excluded.
- Pairs must have a minimum mapQ score. MapQ scores are confusing and how they're calculated varies based on the mapping algorithm being used, but are meant to represent both the number of mismatches in the mapping and how unique that mapping is. With bowtie2, if the read maps equally well to two positions on the genome, its mapQ score will be set to 2. The read in the pair with the higher mapQ is used for the pair.
- Pairs must be above some minimum nucleotide identity (ANI) value. For example if reads in a pair are 100bp each, and each read has a single mismatch, the ANI of that pair would be 0.99

Next, using only read pairs that pass filters, a number of microdiveristy metrics are calculated on a scaffold-by-scaffold basis. This includes:

- Calculate the coverage at each position along the scaffold

- Calculate the clonality at each position along the scaffold in which the coverage is greater than the min_cov argument. The formula for calculating clonality is the sum of the frequency of each base squared - [(frequency of A)^2 + (frequency of C)^2 + (frequency of G)^2 + (frequency of T)^2 ]. This clonality definition is nice because it is not effected by coverage
- Identify SNPs. The criteria for being called a SNP are 1) More than min_cov number of bases at that position, 2) More than min_freq percentage of reads that are a variant base, 3) The number of reads with the variant base is more than the null model for that coverage. The null model describes the probability that the number of true reads that support a variant base could be due to random mutation error, assuming Q30 score. The default false discovery rate with the null model is 1e-6 (one in a million)
- Calculate linkage between SNPs on the same read pair. For each pair harboring a SNP, calculate the linkage of that SNP with other SNPs within that same pair. This is only done for pairs of SNPs that are both on at least MIN_SNP reads
- Calculate scaffold-level properties. These include things like the overall coverage, breadth of coverage, average nucleotide identity (ANI) between the reads and the reference genome, and the expected breadth of coverage based on that true coverage.

Finally, this information is stored as an IS_profile object. This includes the locations of SNPs, the number of read pairs that passed filters (and other information) for each scaffold, the linkage between SNV pairs, ect.

**See also:**

*Example output and explanations* For help interpreting the output

*Advanced use* For access to the raw internal data (which can be very useful)

**choosing_parameters** For information about the pitfalls and other things to consider when running inStrain

To see the command-line options, check the help:

```
$ inStrain profile -h
usage: inStrain profile [-o OUTPUT] [-p PROCESSES] [-d] [-h]
                        [-l FILTER_CUTOFF] [--min_mapq MIN_MAPQ]
                        [--max_insert_relative MAX_INSERT_RELATIVE]
                        [--min_insert MIN_INSERT] [-c MIN_COV] [-f MIN_FREQ]
                        [-fdr FDR] [-s MIN_SNP]
                        [--min_fasta_reads MIN_FASTA_READS]
                        [--store_everything] [--skip_mm_profiling]
                        [--scaffolds_to_profile SCAFFOLDS_TO_PROFILE]
                        bam fasta

REQUIRED:
  bam                   Sorted .bam file
  fasta                 Fasta file the bam is mapped to

I/O PARAMETERS:
  -o OUTPUT, --output OUTPUT
                        Output prefix (default: inStrain)

SYSTEM PARAMETERS:
  -p PROCESSES, --processes PROCESSES
                        Number of processes to use (default: 6)
  -d, --debug           Make extra debugging output (default: False)
  -h, --help            show this help message and exit

READ FILTERING OPTIONS:
  -l FILTER_CUTOFF, --filter_cutoff FILTER_CUTOFF
                        Minimum percent identity of read pairs to consensus to
```

(continues on next page)

(continued from previous page)

```
                        use the reads. Must be >, not >= (default: 0.95)
  --min_mapq MIN_MAPQ   Minimum mapq score of EITHER read in a pair to use
                        that pair. Must be >, not >= (default: -1)
  --max_insert_relative MAX_INSERT_RELATIVE
                        Multiplier to determine maximum insert size between
                        two reads - default is to use 3x median insert size.
                        Must be >, not >= (default: 3)
  --min_insert MIN_INSERT
                        Minimum insert size between two reads - default is 50
                        bp. If two reads are 50bp each and overlap completely,
                        their insert will be 50. Must be >, not >= (default:
                        50)

VARIANT CALLING OPTIONS:
  -c MIN_COV, --min_cov MIN_COV
                        Minimum coverage to call an variant (default: 5)
  -f MIN_FREQ, --min_freq MIN_FREQ
                        Minimum SNP frequency to confirm a SNV (both this AND
                        the FDR snp count cutoff must be true to call a SNP).
                        (default: 0.05)
  -fdr FDR, --fdr FDR   SNP false discovery rate- based on simulation data
                        with a 0.1 percent error rate (Q30) (default: 1e-06)

OTHER OPTIONS:
  -s MIN_SNP, --min_snp MIN_SNP
                        Absolute minimum number of reads connecting two SNPs
                        to calculate LD between them. (default: 20)
  --min_fasta_reads MIN_FASTA_READS
                        Minimum number of reads mapping to a scaffold to
                        proceed with profiling it (default: 0)
  --store_everything    Store intermediate dictionaries in the pickle file;
                        will result in significantly more RAM and disk usage
                        (default: False)
  --skip_mm_profiling   Dont perform analysis on an mm level; saves RAM and
                        time (default: False)
  --scaffolds_to_profile SCAFFOLDS_TO_PROFILE
                        Path to a file containing a list of scaffolds to
                        profile- if provided will ONLY profile those scaffolds
                        (default: None)
```

##### compare

Compare provides the ability to compare two *IS_profile* folders (created by running *inStrain profile*). Both *IS_profile* objects must created based on mapping to the same *.bam* file for *compare* to work.

*inStrain compare* compares a set of different *IS_profile* folders (created by running *inStrain profile*). These *IS_profile* folders represent sets of different sample reads mapped to the same *.fasta* file. To use, we recommend assembly and binning of each sample, and then dereplication of genomes using the software dRep (https://drep.readthedocs.io/) at a high percent ANI, e.g. 96%-99%. Samples which contain multiple populations of the same dRep cluster (members of similar species or sub-species) can then be mapped back to the best genome from this dRep cluster, and then inStrain should be run on these dRep cluster genomes.

**Note:** *inStrain* can only compare read profiles that have been mapped to the same .fasta file

Compare does pair-wise comparisons between each input *IS_profile*. For each pair, a series of steps are undertaken.

1. All positions in which both *IS_profile* objects have at least *min_cov* coverage (5x by default) are identified. This information can be stored in the output by using the flag *–store_coverage_overlap*, but due to it's size, it's not stored by default
2. Each position identified in step 1 is compared. If the flag *–compare_consensus_bases* is used, the consensus base at each position is compared. That means that if the position is 60% A 40% G in sample 1, and 40% A 60% G in sample 2, they will considered different. By default, however, this position would be considered the same. The way that is compares positions is by testing whether the consensus base in sample 1 is detected at all in sample 2 and vice-verse. Detection of an allele in a sample is based on that allele being above the set *-min_freq* and *-fdr*. All detected differences between each pair of samples can be reported if the flag *–store_mismatch_locations* is set.
3. The coverage overlap and the average nucleotide identify for each scaffold is reported. For details on how this is done, see *Example output and explanations*

To see the command-line options, check the help:

```
$ inStrain compare -h
usage: inStrain compare -i [INPUT [INPUT ...]] [-o OUTPUT] [-p PROCESSES] [-d]
                        [-h] [-c MIN_COV] [-f MIN_FREQ] [-fdr FDR]
                        [-s SCAFFOLDS] [--store_coverage_overlap]
                        [--store_mismatch_locations]
                        [--compare_consensus_bases]
                        [--include_self_comparisons] [--greedy_clustering]
                        [--g_ani G_ANI] [--g_cov G_COV] [--g_mm G_MM]

REQUIRED:
  -i [INPUT [INPUT ...]], --input [INPUT [INPUT ...]]
                        A list of inStrain objects, all mapped to the same
                        .fasta file (default: None)
  -o OUTPUT, --output OUTPUT
                        Output prefix (default: instrainComparer)

SYSTEM PARAMETERS:
  -p PROCESSES, --processes PROCESSES
                        Number of processes to use (default: 6)
  -d, --debug           Make extra debugging output (default: False)
  -h, --help            show this help message and exit

VARIANT CALLING OPTIONS:
  -c MIN_COV, --min_cov MIN_COV
                        Minimum coverage to call an variant (default: 5)
  -f MIN_FREQ, --min_freq MIN_FREQ
                        Minimum SNP frequency to confirm a SNV (both this AND
                        the FDR snp count cutoff must be true to call a SNP).
                        (default: 0.05)
  -fdr FDR, --fdr FDR   SNP false discovery rate- based on simulation data
                        with a 0.1 percent error rate (Q30) (default: 1e-06)

OTHER OPTIONS:
  -s SCAFFOLDS, --scaffolds SCAFFOLDS
                        Location to a list of scaffolds to compare. You can
                        also make this a .fasta file and it will load the
                        scaffold names (default: None)
  --store_coverage_overlap
                        Also store coverage overlap on an mm level (default:
                        False)
```

(continues on next page)

(continued from previous page)

```
  --store_mismatch_locations
                        Store the locations of SNPs (default: False)
  --compare_consensus_bases
                        Only compare consensus bases; dont look for lower
                        frequency SNPs when calculating ANI (default: False)
  --include_self_comparisons
                        Also compare IS profiles against themself (default:
                        False)

GREEDY CLUSTERING OPTIONS [THIS SECTION IS EXPERIMENTAL!]:
  --greedy_clustering   Dont do pair-wise comparisons, do greedy clustering to
                        only find the number of clsuters. If this is set, use
                        the parameters below as well (default: False)
  --g_ani G_ANI         ANI threshold for greedy clustering- put the fraction
                        not the percentage (e.g. 0.99, not 99) (default: 0.99)
  --g_cov G_COV         Alignment coverage for greedy clustering- put the
                        fraction not the percentage (e.g. 0.5, not 10)
                        (default: 0.99)
  --g_mm G_MM           Maximum read mismatch level (default: 100)
```

##### profile_genes

After running *inStrain profile* on a sample, you can calculate the coverage, microdiveristy, and SNP type for each gene. You do this by providing a file of gene calls. See doc:*example_output* for example results, and doc:*preparing_input* for information about creating the input file.

To see the command-line options, check the help:

```
$ inStrain profile_genes -h
usage: inStrain profile_genes -i IS -g GENE_FILE [-p PROCESSES] [-d] [-h]

REQUIRED:
-i IS, --IS IS         an inStrain profile object (default: None)
-g GENE_FILE, --gene_file GENE_FILE
                       Path to prodigal .fna genes file. (default: None)

SYSTEM PARAMETERS:
-p PROCESSES, --processes PROCESSES
                       Number of processes to use (default: 6)
-d, --debug            Make extra debugging output (default: False)
-h, --help             show this help message and exit
```

##### genome_wide

After running *inStrain profile*, most results are presented on a scaffold-by-scaffold basis. To have the results summarized in a genome-by-genome way instead, you can use the module *inStrain genome_wide*. It is also required to run this module before making plots.

There are a number of ways of telling *inStrain* which scaffold belongs to which genome

1. Individual .fasta files. As recommended in preparing_input, if you want to run *inStrain* on multiple genomes in the same sample, you should first concatenate all of the individual genomes into a single *.fasta* file and map to that. To view the results of the individual genomes used to create the concatenated .fasta file, you can pass a list of the individual *.fasta files to *inStrain genome_wide*. (e.g. inStrain genome_wide -i inStrain_folder -s genome1.fasta genome2.fasta genome3.fasta)

2. Scaffold to bin file. This text file consists of two columns, with one column listing the scaffold name, and the second column listing the genome bin name. Columns should be separated by tabs.
3. Nothing. If all of your scaffolds belong to the same genome, by running *inStrain genome_wide* without any *-s* options it will summarize the results of all scaffolds together.

The flag *–mm_level* produces output for each mm. You probably don't want this. For information on what I mean by mm_level see *Advanced use*, for information on the output see *Example output and explanations*

To see the command-line options, check the help:

```
$ inStrain genome_wide -h
usage: inStrain genome_wide -i IS [-s [STB [STB ...]]] [--mm_level]
                          [-p PROCESSES] [-d] [-h]

REQUIRED:
-i IS, --IS IS        an inStrain profile object (default: None)
-s [STB [STB ...]], --stb [STB [STB ...]]
                      Scaffold to bin. This can be a file with each line
                      listing a scaffold and a bin name, tab-seperated. This
                      can also be a space-seperated list of .fasta files,
                      with one genome per .fasta file. If nothing is
                      provided, all scaffolds will be treated as belonging
                      to the same genome (default: [])
--mm_level            Create files on the mm level (see documentation for
                      info) (default: False)

SYSTEM PARAMETERS:
-p PROCESSES, --processes PROCESSES
                      Number of processes to use (default: 6)
-d, --debug           Make extra debugging output (default: False)
-h, --help            show this help message and exit
```

##### quick_profile

This is a quirky module that is not really related to any of the others. It is used to quickly profile a *.bam* file to pull out scaffolds from genomes that are at a sufficient breadth.

To use it you must provide a *.bam* file, the *.fasta* file that you mapped to to generate the *.bam* file, and a *scaffold to bin* file (see above section for details). The *stringent_breadth_cutoff* removed scaffolds entirely which have less breath than this (used to make the program run faster and produce smaller output). All scaffolds from genomes with at least the *breadth_cutoff* are then written to a file. In this way, you can then choose to run inStrain profile only on scaffolds from genomes that known to be of sufficient breadth, speeding up the run and reducing RAM usage (though not by much).

To see the command-line options, check the help:

```
$ inStrain quick_profile -h
usage: inStrain quick_profile -b BAM -f FASTA -s STB [-o OUTPUT]
                            [-p PROCESSES] [-d] [-h]
                            [--breadth_cutoff BREADTH_CUTOFF]
                            [--stringent_breadth_cutoff STRINGENT_BREADTH_CUTOFF]

REQUIRED:
-b BAM, --bam BAM     A bam file to profile (default: None)
-f FASTA, --fasta FASTA
                      The .fasta file to profile (default: None)
```

(continues on next page)

(continued from previous page)

```
-s STB, --stb STB     Scaffold to bin file for genome-wide coverage and
                      breadth (default: None)
-o OUTPUT, --output OUTPUT
                      Output prefix (default: None)

SYSTEM PARAMETERS:
-p PROCESSES, --processes PROCESSES
                      Number of processes to use (default: 6)
-d, --debug           Make extra debugging output (default: False)
-h, --help            show this help message and exit

OTHER OPTIONS:
--breadth_cutoff BREADTH_CUTOFF
                      Minimum breadth to pull scaffolds (default: 0.5)
--stringent_breadth_cutoff STRINGENT_BREADTH_CUTOFF
                      Minimum breadth to let scaffold into coverm raw
                      results (default: 0.01)
```

##### plot

This module produces plots based on the results of *inStrain profile* and *inStrain compare*. In both cases, before plots can be made, *inStrain genome_wide* must be run on the output folder first. In order to make plots 8 and 9, *inStrain profile_genes* must be run first as well.

The recommended way of running this module is with the default *-pl a*. It will just try and make all of the plots that it can, and will tell you about any plots that it fails to make.

See *Example output and explanations* for an example of the plots it can make.

To see the command-line options, check the help:

```
$ inStrain plot -h
usage: inStrain plot -i IS [-pl [PLOTS [PLOTS ...]]] [-p PROCESSES] [-d] [-h]

REQUIRED:
  -i IS, --IS IS        an inStrain profile object (default: None)
  -pl [PLOTS [PLOTS ...]], --plots [PLOTS [PLOTS ...]]
                        Plots. Input 'all' or 'a' to plot all
                        1) Coverage and breadth vs. read mismatches
                        2) Genome-wide microdiversity metrics
                        3) Read-level ANI distribution
                        4) Major allele frequencies
                        5) Linkage decay
                        6) Read filtering plots
                        7) Scaffold inspection plot (large)
                        8) Linkage with SNP type (GENES REQUIRED)
                        9) Gene histograms (GENES REQUIRED)
                        10) Compare dendrograms (RUN ON COMPARE; NOT PROFILE)
                         (default: a)

SYSTEM PARAMETERS:
  -p PROCESSES, --processes PROCESSES
                        Number of processes to use (default: 6)
  -d, --debug           Make extra debugging output (default: False)
  -h, --help            show this help message and exit
```

##### other

This module holds odds and ends functionalities. As of version 1.0.0, all it can do is convert old *IS_profile* objects (>v0.3.0) to newer versions (v0.8.0). As the code base around *inStrain* matures, we expect more functionalities to be included here.

To see the command-line options, check the help:

```
$ inStrain other -h
usage: inStrain other [-p PROCESSES] [-d] [-h] [--old_IS OLD_IS]

SYSTEM PARAMETERS:
  -p PROCESSES, --processes PROCESSES
                        Number of processes to use (default: 6)
  -d, --debug           Make extra debugging output (default: False)
  -h, --help            show this help message and exit

OTHER OPTIONS:
  --old_IS OLD_IS       Convert an old inStrain version object to the newer
                        version. (default: None)
```

#### 1.4 Example output and explanations

InStrain produces a variety of output in the IS folder depending on which operations are run. Generally, output that is meant for human eyes to be easily interpretable is located in the `output` folder.

##### 1.4.1 inStrain profile

A typical run of inStrain will yield the following files in the output folder:

###### scaffold_info.tsv

This gives basic information about the scaffolds in your sample at the highest allowed level of read identity.

Table 1: scaffold_info.tsv

| scaffold | length | breadth | coverage | median_cov | std_cov | [illegible] | [illegible] | [illegible] | [illegible] | [illegible] | [illegible] Breadth | [illegible] | SNPs | Reference_SNPs | BiAllelic_SNPs | MultiAllelic_SNPs | consensus_SNPs | population_SNPs | conANI | popANI |
|---|---|---|---|---|---|---|---|---|---|---|---|---|---|---|---|---|---|---|---|---|
| S3_[illegible] | [illegible] | [illegible] | [illegible] | [illegible] | [illegible] | [illegible] | [illegible] | [illegible] | [illegible] | [illegible] | [illegible] | [illegible] | [illegible] | [illegible] | [illegible] | 0 | 0 | 0 | 1.0 | 1.0 |
| S3_[illegible] | [illegible] | [illegible] | [illegible] | [illegible] | [illegible] | [illegible] | [illegible] | [illegible] | | | 0.0 | [illegible] | [illegible] | [illegible] | [illegible] | 0 | 0 | 0 | 0.0 | 0.0 |
| S3_[illegible] | [illegible] | [illegible] | [illegible] | [illegible] | [illegible] | [illegible] | [illegible] | [illegible] | | | 0.0 | [illegible] | [illegible] | [illegible] | [illegible] | 0 | 0 | 0 | 0.0 | 0.0 |
| S3_[illegible] | [illegible] | [illegible] | [illegible] | [illegible] | [illegible] | [illegible] | [illegible] | [illegible] | | | 0.0 | [illegible] | [illegible] | [illegible] | [illegible] | 0 | 0 | 0 | 0.0 | 0.0 |
| S3_[illegible] | [illegible] | [illegible] | [illegible] | [illegible] | [illegible] | [illegible] | [illegible] | [illegible] | 0.0 | 0.0 | [illegible] | [illegible] | [illegible] | [illegible] | [illegible] | 0 | 0 | 0 | 1.0 | 1.0 |

**scaffold** The name of the scaffold in the input .fasta file

**length** Full length of the scaffold in the input .fasta file

**breadth** The percentage of bases in the scaffold that are covered by at least a single read. A breadth of 1 means that all bases in the scaffold have at least one read covering them

**coverage** The average depth of coverage on the scaffold. If half the bases in a scaffold have 5 reads on them, and the other half have 10 reads, the coverage of the scaffold will be 7.5

**median_cov** The median coverage value of all bases in the scaffold, included bases with 0 coverage

**std_cov** The standard deviation of all coverage values

**bases_w_0_coverage** The number of bases with 0 coverage

**mean_clonality** The mean clonality value of all bases in the scaffold that have a clonality value calculated. So if only 1 base on the scaffold meats the minimum coverage to calculate clonality, the mean_clonality of the scaffold will be the clonality of that base

**median_clonality** The median clonality value of all bases in the scaffold that have a clonality value calculated

**mean_microdiversity** The mean mean_microdiversity value of all bases in the scaffold that have a mean_microdiversity value calculated (microdiveristy = 1 - clonality)

**median_microdiversity** The median microdiversity value of all bases in the scaffold that have a microdiversity value calculated

**unmaskedBreadth** The percentage of bases in the scaffold that have at least the min_cov number of bases. This value multiplied by the length of the scaffold gives the percentage of bases for which clonality is calculated and on which SNPs can be called

**SNPs** The total number of SNPs called on this scaffold

**expected_breadth** This tells you the breadth that you should expect if reads are evenly distributed along the genome, given the reported coverage value. Based on the function breadth = -1.000 * e^(0.883 * coverage) + 1.000. This is useful to establish whether or not the scaffold is actually in the reads, or just a fraction of the scaffold. If your coverage is 10x, the expected breadth will be ~1. If your actual breadth is significantly lower then the expected breadth, this means that reads are mapping only to a specific region of your scaffold (transposon, etc.)

**SNPs** The total number of SNPs called on this scaffold

**Referece_SNPs** The number of SNPs called on this scaffold with allele_count = 1. This means that the only allele detected in the reads is different from the reference base

**BiAllelic_SNPs** The number of SNPs called on this scaffold with allele_count = 2. This means that there are two possible alleles at this position

**MultiAllelic_SNPs** The number of SNPs called on this scaffold with allele_count > 2. This means that there are more than two possible alleles at this position

**consensus_SNPs** The number of SNPs called on this scaffold with allele_count > 0 **and** where consensus base is not the reference base. This should be the same as Reference_SNPs under almost all circumstances

**population_SNPs** These are SNPs where the reference base isn't detected at all, regardless of the allele count.

**conANI** The average nucleotide identity between the reads in the sample and the .fasta file based on consensus SNPs. Calculated using the formula ANI = (unmaskedBreadth * length) - consensus_SNPs)/ (unmaskedBreadth * length))

**popANI** The average nucleotide identity between the reads in the sample and the .fasta file based on consensus SNPs. Calculated using the formula ANI = (unmaskedBreadth * length) - population_SNPs)/ (unmaskedBreadth * length))

##### read_report.tsv

This provides an overview of the number of reads that map to each scaffold, and some basic metrics about their quality.

Table 2: read_report.tsv

| scaffold | unfiltered_reads | unfiltered_pairs | pass_filter_cutoff | pass_max_insert | pass_min_insert | pass_min_mapq | filtered_pairs | mean_mistmaches | mean_insert_distance | mean_mapq_score | mean_pair_length | median_insert | mean_PID |
|---|---|---|---|---|---|---|---|---|---|---|---|---|---|
| all_scaffolds | 3802370 | 1790817 | 1674511 | 1784011 | 1790699 | 1790817 | 1766849 | [illegible] | [illegible] | [illegible] | [illegible] | [illegible] | [illegible] |
| S3_002_000X1_scaffold_1162 | 12 | 6 | 6 | 6 | 6 | 6 | 6 | 1.0 | 281.1666[illegible] | [illegible] | [illegible] | 287.0 | 0.9966666666666668 |
| S3_002_000X1_scaffold_1005 | 10 | 5 | 5 | 5 | 5 | 5 | 5 | 0.2 | 318.0 | 33.2 | 299.8 | 208.0 | 0.9993333333333332 |
| S3_002_006X1_scaffold_1151 | 6 | 3 | 3 | 3 | 3 | 3 | 3 | 5.6666[illegible] | [illegible] | [illegible] | [illegible] | [illegible] | 0.9811111111111112 |
| S3_002_000X1_scaffold_1004 | [illegible] | 6 | 6 | 6 | 6 | 6 | 6 | 0.5 | 295.5 | 16.6666[illegible] | [illegible] | [illegible] | 0.9983333333333334 |

The following metrics are provided for all individual scaffolds, and for all scaffolds together (scaffold "all_scaffolds"). For the max insert cutoff, the median_insert for all_scaffolds is used

**header line** The header line (starting with #; not shown in the above table) describes the parameters that were used to filter the reads

**scaffold** The name of the scaffold in the input .fasta file

**unfiltered_reads** The raw number of reads that map to this scaffold

**unfiltered_pairs** The raw number of pairs of reads that map to this scaffold. Only paired reads are used by inStrain

**pass_filter_cutoff** The number of pairs of reads mapping to this scaffold that pass the ANI filter cutoff (specified in the header as "filter_cutoff")

**pass_max_insert** The number of pairs of reads mapping to this scaffold that pass the maximum insert size cutoff- that is, their insert size is less than 3x the median insert size of all_scaffolds. Note that the insert size is measured from the start of the first read to the end of the second read (2 perfectly overlapping 50bp reads will have an insert size of 50bp)

**pass_min_insert** The number of pairs of reads mapping to this scaffold that pass the minimum insert size cutoff

**pass_min_mapq** The number of pairs of reads mapping to this scaffold that pass the minimum mapQ score cutoff

**filtered_pairs** The number of pairs of reads that pass all cutoffs

**mean_mistmaches** Among all pairs of reads mapping to this scaffold, the mean number of mismatches

**mean_insert_distance** Among all pairs of reads mapping to this scaffold, the mean insert distance. Note that the insert size is measured from the start of the first read to the end of the second read (2 perfectly overlapping 50bp reads will have an insert size of 50bp)

**mean_mapq_score** Among all pairs of reads mapping to this scaffold, the average mapQ score

**mean_pair_length** Among all pairs of reads mapping to this scaffold, the average length of both reads in the pair summed together

**median_insert** Among all pairs of reads mapping to this scaffold, the median insert distance.

**mean_PID** Among all pairs of reads mapping to this scaffold, the average percentage ID of both reads in the pair to the reference .fasta file

#### SNVs.tsv

This describes the SNPs that are detected in this mapping.

Table 3: SNVs.tsv

| scaffold | position | refBase | A | C | T | G | conBase | varBase | allele_count | cryptic | baseCoverage | varFreq | refFreq |
|---|---|---|---|---|---|---|---|---|---|---|---|---|---|
| S3_003_000X1_scaffold_[illegible] | 838 | [illegible] | 2 | 7 | 0 | 0 | C | A | 2 | False | 9 | 0.2222222222[illegible] | [illegible]7777777778 |
| S3_003_000X1_scaffold_[illegible] | 99 | [illegible] | 0 | 0 | 5 | 0 | T | A | 1 | False | 5 | 0.0 | 1.0 |
| S3_003_000X1_scaffold_[illegible] | 123 | [illegible] | 0 | 0 | 0 | 11 | G | A | 1 | False | 11 | 0.0 | 1.0 |
| S3_003_000X1_scaffold_[illegible] | 261 | [illegible] | 19 | 0 | 0 | 0 | A | A | 1 | False | 19 | 1.0 | 1.0 |
| S3_003_000X1_scaffold_[illegible] | 291 | [illegible] | 0 | 16 | 2 | 0 | C | T | 2 | False | 18 | 0.1111111111[illegible] | [illegible]8888888888 |

See the module_descriptions for what constitutes a SNP (what makes it into this table)

**scaffold** The scaffold that the SNP is on

**position** The genomic position of the SNP

**refBase** The reference base in the .fasta file at that position

**A, C, T, and G** The number of mapped reads encoding each of the bases

**conBase** The consensus base; the base that is supported by the most reads

**varBase** Variant base; the base with the second most reads

**morphia** The number of bases that are detected above background levels. In order to be detected above background levels, you must pass an fdr filter. See module descriptions for a description of how that works. A morphia of 0 means no bases are supported by the reads, a morphia of 1 means that only 1 base is supported by the reads, a morphia of 2 means two bases are supported by the reads, etc.

**cryptic** If a SNP is cryptic, it means that it is detected when using a lower read mismatch threshold, but becomes undetected when you move to a higher read mismatch level. See "dealing with mm" in the advanced_use section for more details on what this means.

**baseCoverage** The total number of reads at this position

**varFreq** The fraction of reads supporting the varBase

**refFreq** The fraction of reds supporting the refBase

**conFreq** The fraction of reds supporting the conBase

#### linkage.tsv

This describes the linkage between pairs of SNPs in the mapping that are found on the same read pair at least min_snp times.

Table 4: linkage.tsv

| r2 | d_prime | r2_normalized | d_prime_normalized | total | countAB | countAb | countaB | countab | allele_A | allele_a | allele_B | allele_b | distance | position_A | position_B | scaffold |
|---|---|---|---|---|---|---|---|---|---|---|---|---|---|---|---|---|
| 1.0 | 1.0 | 1.0 | 1.0 | 27 | 0 | 14 | 13 | 0 | G | A | T | C | 45 | 191425 | 191470 | S3_003_000X1_scaffol[illegible] |
| 0.1074[illegible] | [illegible] | [illegible] | [illegible] | [illegible] | [illegible] | 0 | 9 | 2 | G | A | C | A | 80 | 191425 | 191505 | S3_003_000X1_scaffol[illegible] |
| 0.0833[illegible] | [illegible] | [illegible] | [illegible] | [illegible] | [illegible] | 2 | 13 | 0 | T | C | C | A | 35 | 191470 | 191505 | S3_003_000X1_scaffol[illegible] |
| 1.0000[illegible] | [illegible] | [illegible] | 1.0 | 30 | 22 | 0 | 0 | 8 | C | T | T | C | 12 | 99342 | 99354 | S3_003_000X1_scaffol[illegible] |
| 1.0000[illegible] | [illegible] | [illegible] | 1.0 | 22 | 17 | 0 | 0 | 5 | C | T | T | A | 60 | 99342 | 99402 | S3_003_000X1_scaffol[illegible] |

Linkage is used primarily to determine if organisms are undergoing horizontal gene transfer or not. It's calculated for pairs of SNPs that can be connected by at least `min_snp` reads. It's based on the assumption that each SNP as two alleles (for example, a A and b B). This all gets a bit confusing and has a large amount of literature around each of these terms, but I'll do my best to briefly explain what's going on

**scaffold** The scaffold that both SNPs are on

**position_A** The position of the first SNP on this scaffold

**position_B** The position of the second SNP on this scaffold

**distance** The distance between the two SNPs

**allele_A** One of the two bases at position_A

**allele_a** The other of the two bases at position_A

**allele_B** One of the bases at position_B

**allele_b** The other of the two bases at position_B

**countAB** The number of read-pairs that have allele_A and allele_B

**countAb** The number of read-pairs that have allele_A and allele_b

**countaB** The number of read-pairs that have allele_a and allele_B

**countab** The number of read-pairs that have allele_a and allele_b

**total** The total number of read-pairs that have have information for both position_A and position_B

**r2** This is the r-squared linkage metric. See below for how it's calculated

**d_prime** This is the d-prime linkage metric. See below for how it's calculated

**r2_normalized, d_prime_normalized** These are calculated by rarefying to `min_snp` number of read pairs. See below for how it's calculated

Python code for the calculation of these metrics:

```
freq_AB = float(countAB) / total
freq_Ab = float(countAb) / total
freq_aB = float(countaB) / total
freq_ab = float(countab) / total

freq_A = freq_AB + freq_Ab
freq_a = freq_ab + freq_aB
freq_B = freq_AB + freq_aB
freq_b = freq_ab + freq_Ab

linkD = freq_AB - freq_A * freq_B

if freq_a == 0 or freq_A == 0 or freq_B == 0 or freq_b == 0:
    r2 = np.nan
else:
    r2 = linkD*linkD / (freq_A * freq_a * freq_B * freq_b)

linkd = freq_ab - freq_a * freq_b

# calc D-prime
d_prime = np.nan
if (linkd < 0):
    denom = max([(-freq_A*freq_B),(-freq_a*freq_b)])
    d_prime = linkd / denom
```

(continues on next page)

(continued from previous page)

```
elif (linkD > 0):
    denom = min([(freq_A*freq_b), (freq_a*freq_B)])
    d_prime = linkd / denom

################
# calc rarefied

rareify = np.random.choice(['AB','Ab','aB','ab'], replace=True, p=[freq_AB,freq_Ab,
↪freq_aB,freq_ab], size=min_snp)
freq_AB = float(collections.Counter(rareify)['AB']) / min_snp
freq_Ab = float(collections.Counter(rareify)['Ab']) / min_snp
freq_aB = float(collections.Counter(rareify)['aB']) / min_snp
freq_ab = float(collections.Counter(rareify)['ab']) / min_snp

freq_A = freq_AB + freq_Ab
freq_a = freq_ab + freq_aB
freq_B = freq_AB + freq_aB
freq_b = freq_ab + freq_Ab

linkd_norm = freq_ab - freq_a * freq_b

if freq_a == 0 or freq_A == 0 or freq_B == 0 or freq_b == 0:
    r2_normalized = np.nan
else:
    r2_normalized = linkd_norm*linkd_norm / (freq_A * freq_a * freq_B * freq_b)

# calc D-prime
d_prime_normalized = np.nan
if (linkd_norm < 0):
    denom = max([(-freq_A*freq_B),(-freq_a*freq_b)])
    d_prime_normalized = linkd_norm / denom

elif (linkd_norm > 0):
    denom = min([(freq_A*freq_b), (freq_a*freq_B)])
    d_prime_normalized = linkd_norm / denom

rt_dict = {}
for att in ['r2', 'd_prime', 'r2_normalized', 'd_prime_normalized', 'total', 'countAB
↪', \
            'countAb', 'countaB', 'countab', 'allele_A', 'allele_a', \
            'allele_B', 'allele_b']:
    rt_dict[att] = eval(att)
```

##### 1.4.2 inStrain compare

A typical run of inStrain will yield the following files in the output folder:

Table 5: comparisonsTable.tsv

| scaffold | name1 | name2 | coverage_overlap | compared_bases_count | percent_genome_compared | length | consensus_SNPs | population_SNPs | conANI | popANI |
|---|---|---|---|---|---|---|---|---|---|---|
| S3_016_000X1_scaffold_1420 | Sloan3AlfGenomeInventory.fasta-vs-S3_003_000X1.sorted.bam | Sloan3AlfGenomeInventory.fasta-vs-S3_016_000X1.sorted.bam | [illegible] | [illegible] | [illegible] | [illegible] | [illegible] | 0 | [illegible] | [illegible] |
| S3_016_000X1_scaffold_9493 | Sloan3AlfGenomeInventory.fasta-vs-S3_003_000X1.sorted.bam | Sloan3AlfGenomeInventory.fasta-vs-S3_016_000X1.sorted.bam | [illegible] | [illegible] | [illegible] | [illegible] | [illegible] | 0 | [illegible] | [illegible] |
| S3_016_000X1_scaffold_12686 | Sloan3AlfGenomeInventory.fasta-vs-S3_003_000X1.sorted.bam | Sloan3AlfGenomeInventory.fasta-vs-S3_016_000X1.sorted.bam | [illegible] | [illegible] | [illegible] | [illegible] | [illegible] | 0 | [illegible] | [illegible] |
| S3_016_000X1_scaffold_11829 | Sloan3AlfGenomeInventory.fasta-vs-S3_003_000X1.sorted.bam | Sloan3AlfGenomeInventory.fasta-vs-S3_016_000X1.sorted.bam | [illegible] | [illegible] | [illegible] | [illegible] | [illegible] | 0 | [illegible] | [illegible] |
| S3_016_000X1_scaffold_889 | Sloan3AlfGenomeInventory.fasta-vs-S3_003_000X1.sorted.bam | Sloan3AlfGenomeInventory.fasta-vs-S3_016_000X1.sorted.bam | [illegible] | [illegible] | [illegible] | [illegible] | [illegible] | 0 | [illegible] | [illegible] |

**scaffold** The scaffold being compared

**name1** The name of the first *inStrain profile* being compared

**name2** The name of the second *inStrain profile* being compared

**coverage_overlap** The percentage of bases that are either covered or not covered in both of the profiles (covered = the base is present at at least min_snp coverage). The formula is length(coveredInBoth) / length(coveredInEither). If both scaffolds have 0 coverage, this will be 0.

**compared_bases_count** The number of considered bases; that is, the number of bases with at least min_snp coverage in both profiles. Formula is length([x for x in overlap if x == True]).

**percent_genome_compared** The percentage of bases in the scaffolds that are covered by both. The formula is length([x for x in overlap if x == True])/length(overlap). When ANI is np.nan, this must be 0. If both scaffolds have 0 coverage, this will be 0.

**length** The total length of the scaffold

**consensus_SNPs** The number of locations along the genome where both samples have the base at >= 5x coverage, and the consensus allele in each sample is different

**population_SNPs** The number of locations along the genome where both samples have the base at >= 5x coverage, and no alleles are shared between either sample. See inStrain manuscript for more details.

**popANI** The average nucleotide identity among compared bases between the two scaffolds, based on population_SNPs. Calculated using the formula popANI = (compared_bases_count - population_SNPs) / compared_bases_count

**conANI** The average nucleotide identity among compared bases between the two scaffolds, based on consensus_SNPs. Calculated using the formula conANI = (compared_bases_count - consensus_SNPs) / compared_bases_count

##### 1.4.3 inStrain profile_genes

A typical run of inStrain profile_genes will yield the following additional files in the output folder:

##### gene_info.tsv

This describes some basic information about the genes being profiled

Table 6: gene_info.tsv

| gene | scaffold | direction | partial | start | end | coverage | breadth | clonality | microdiversity | masked_breadth | SNPs_per_bp | min_ANI |
|---|---|---|---|---|---|---|---|---|---|---|---|---|
| S3_002_028G1_scaffold_0_1 | S3_002_028G1_scaffold_0 | -1 | False | 957 | 2219 | | | | | | | 0 |
| S3_002_028G1_scaffold_0_2 | S3_002_028G1_scaffold_0 | -1 | False | 2189 | 3136 | | | | | | | 0 |
| S3_002_028G1_scaffold_0_3 | S3_002_028G1_scaffold_0 | 1 | False | 3274 | 5013 | | | | | | | 0 |
| S3_002_028G1_scaffold_0_4 | S3_002_028G1_scaffold_0 | -1 | False | 5018 | 5746 | | | | | | | 0 |
| S3_002_028G1_scaffold_0_5 | S3_002_028G1_scaffold_0 | 1 | False | 5888 | 6862 | | | | | | | 0 |

**gene** Name of the gene being profiled

**scaffold** Scaffold that the gene is on

**direction** Direction of the gene (based on prodigal call). If -1, means the gene is not coded in the direction expressed by the .fasta file

**partial** If True this is a partial gene; based on not having *partial=00* in the record description provided by Prodigal

**start** Start of the gene (position on scaffold; 0-indexed)

**end** End of the gene (position on scaffold; 0-indexed)

**coverage** The mean coverage across the length of the gene

**breadth** The number of bases in the gene that have at least 1x coverage

**microdiversity** The mean nucleotide diversity (pi) among positions on the gene with at least 5x coverage

**clonality** 1 - microdiversity

**masked_breadth** The percentage of positions in the gene with at least 5x coverage

**SNPs_per_bp** The number of positions on the gene where a SNP is called

**min_ANI** The minimum read ANI level when profile_genes was run (0 means the value is whatever was set with Profile was originally run)

##### SNP_mutation_types.tsv

This describes whether SNPs are synonymous, nonsynonymous, or intergenic

Table 7: SNP_mutation_types.tsv

| scaffold | position | refBase | A | C | T | G | conBase | varBase | allele_count | baseCoverage | varFreq | refFreq | mutation_type | mutation | gene |
|---|---|---|---|---|---|---|---|---|---|---|---|---|---|---|---|
| S3_002_056W1_scaffold_121 | 2[illegible]34 | C | 0 | 3 | 2 | 0 | C | T | 2 | 5 | 0.4 | 0.6 | N | N:H936Y | S3_002_056W1_scaffold_12 |
| S3_002_056W1_scaffold_121 | 8509 | G | 7 | 0 | 0 | 0 | A | A | 1 | 7 | 1.0 | 1.0 | N | N:G459R | S3_002_056W1_scaffold_12 |
| S3_002_056W1_scaffold_121 | 8510 | G | 7 | 0 | 0 | 0 | A | A | 1 | 7 | 1.0 | 1.0 | N | N:G460E | S3_002_056W1_scaffold_12 |
| S3_002_056W1_scaffold_121 | 16899 | G | 0 | 2 | 0 | 5 | G | C | 2 | 7 | 0.2857142857142857 | [illegible] | [illegible] | N:G1068R | S3_002_056W1_scaffold_12 |
| S3_002_056W1_scaffold_121 | 24347 | C | 0 | 9 | 2 | 0 | C | T | 2 | 11 | 0.18181818181818182 | [illegible] | [illegible] | N:Q294* | S3_002_056W1_scaffold_12 |

All genes with an allele_count of 1 or 2 make it into this table; see the above description of SNVs.tsv for details on what most of these columns mean

**mutation_type** What type of mutation this is. N = nonsynonymous, S = synonymous, I = intergenic, M = there are multiple genes with this base so you cant tell

**mutation** Short-hand code for the amino acid switch. If synonymous, this will be S: + the position. If nonsynonymous, this will be N: + the old amino acid + the position + the new amino acid.

**gene** The gene this SNP is in

#### 1.4.4 inStrain genome_wide

A typical run of inStrain genome_wide will yield the following additional files in the output folder:

##### genomeWide_scaffold_info.tsv

This is a genome-wide version of the scaffold report described above. See above for column descriptions.

Table 8: genomeWide_scaffold_info.tsv

| genome | detected_scaffolds | true_scaffolds | length | SNPs | Referece_SNPs | BiAllelic_SNPs | MultiAllelic_SNPs | consensus_SNPs | population_SNPs | breadth | coverage | std_cov | mean_clonality | conANI | popANI | unmaskedBreadth | expected_breadth |
|---|---|---|---|---|---|---|---|---|---|---|---|---|---|---|---|---|---|
| [illegible] | [illegible] | [illegible] | [illegible] | [illegible] | [illegible] | [illegible] | [illegible] | [illegible] | [illegible] | [illegible] | [illegible] | [illegible] | [illegible] | [illegible] | [illegible] | [illegible] | [illegible] |
| [illegible] | [illegible] | [illegible] | [illegible] | [illegible] | [illegible] | [illegible] | [illegible] | [illegible] | [illegible] | [illegible] | [illegible] | [illegible] | [illegible] | [illegible] | [illegible] | 0.0 | 0.08543195678460236 |
| [illegible] | [illegible] | [illegible] | [illegible] | [illegible] | [illegible] | [illegible] | [illegible] | [illegible] | [illegible] | [illegible] | [illegible] | [illegible] | [illegible] | [illegible] | [illegible] | [illegible] | [illegible] |
| [illegible] | [illegible] | [illegible] | [illegible] | [illegible] | [illegible] | [illegible] | [illegible] | [illegible] | [illegible] | [illegible] | [illegible] | [illegible] | [illegible] | [illegible] | [illegible] | [illegible] | [illegible] |
| [illegible] | [illegible] | [illegible] | [illegible] | [illegible] | [illegible] | [illegible] | [illegible] | [illegible] | [illegible] | [illegible] | [illegible] | [illegible] | [illegible] | [illegible] | [illegible] | [illegible] | [illegible] |

##### genomeWide_read_report.tsv

This is a genome-wide version of the read report described above. See above for column descriptions.

Table 9: genomeWide_read_report.tsv

| genome | scaffolds | unfiltered_reads | unfiltered_pairs | pass_filter_cutoff | pass_max_insert | pass_min_insert | pass_min_mapq | filtered_pairs | mean_mistmaches | mean_insert_distance | mean_mapq_score | mean_pair_length | median_insert | mean_PID |
|---|---|---|---|---|---|---|---|---|---|---|---|---|---|---|
| [illegible] | [illegible] | [illegible] | [illegible] | [illegible] | [illegible] | 5062 | 5062 | 5048 | [illegible] | [illegible] | [illegible] | [illegible] | [illegible] | [illegible] |
| [illegible] | [illegible] | [illegible] | [illegible] | [illegible] | [illegible] | [illegible] | [illegible] | [illegible] | [illegible] | [illegible] | [illegible] | [illegible] | [illegible] | [illegible] |

#### 1.4.5 inStrain plot

This is what the results of inStrain plot look like.

##### 1) Coverage and breadth vs. read mismatches

Breadth of coverage (blue line), coverage depth (red line), and expected breadth of coverage given the depth of coverage (dotted blue line) versus the minimum ANI of mapped reads. Coverage depth continues to increase while breadth of plateaus, suggesting that all regions of the reference genome are not present in the reads being mapped.

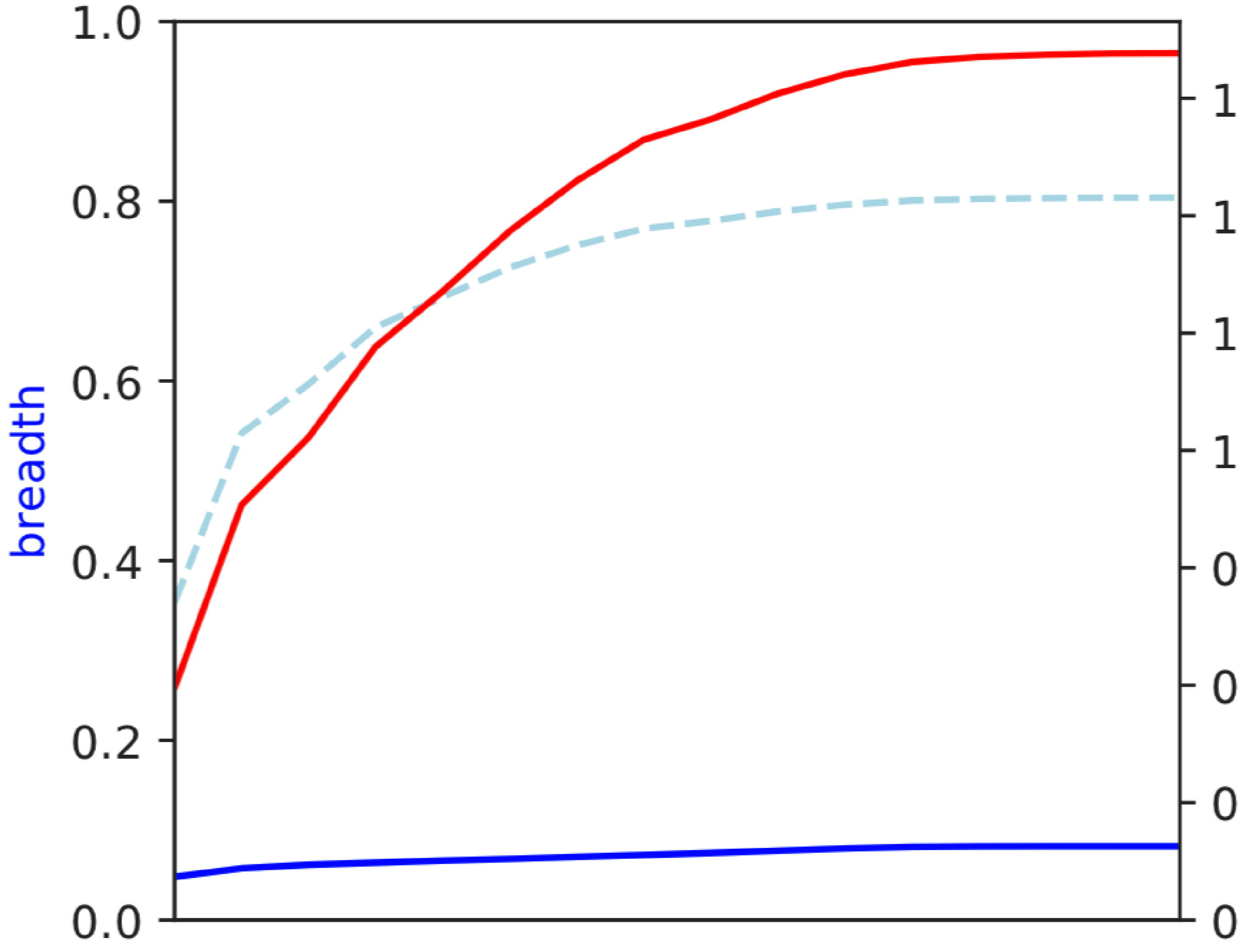
1.0
0.8
0.6
0.4
0.2
0.0
breadth
1
1
1
1
0
0
0
0

##### 2) Genome-wide microdiversity metrics

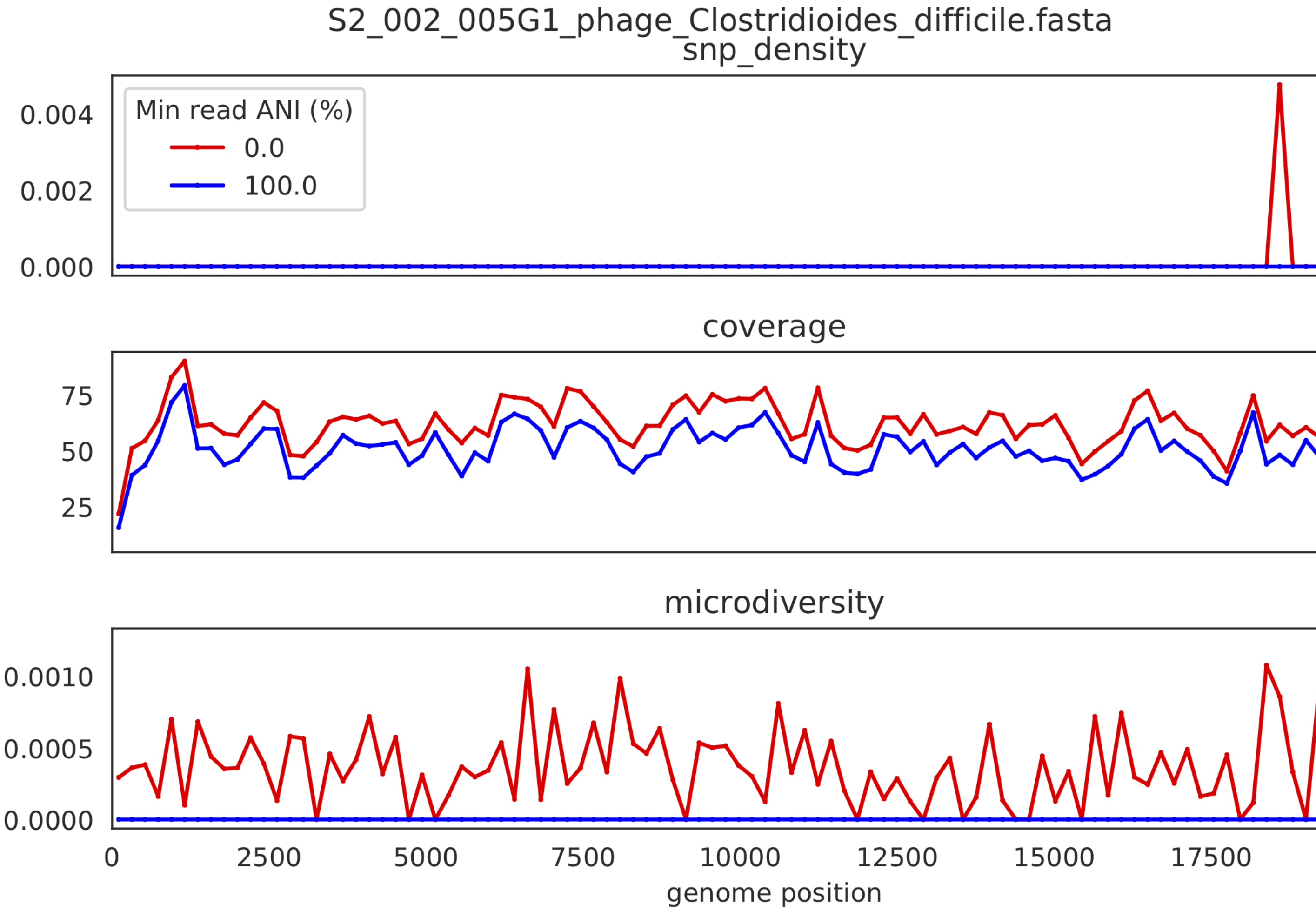

SNV density, coverage, and nucleotide diversity. Spikes in nucleotide diversity and SNV density do not correspond with increased coverage, indicating that the signals are not due to read mis-mapping. Positions with nucleotide diversity and no SNV-density are those where diversity exists but is not high enough to call a SNV

##### 3) Read-level ANI distribution

Distribution of read pair ANI levels when mapped to a reference genome; this plot suggests that the reference genome is >1% different than the mapped reads

##### 4) Major allele frequencies

Distribution of the major allele frequencies of bi-allelic SNVs (the Site Frequency Spectrum). Alleles with major frequencies below 50% are the result of multiallelic sites. The lack of distinct puncta suggest that more than a few distinct strains are present.

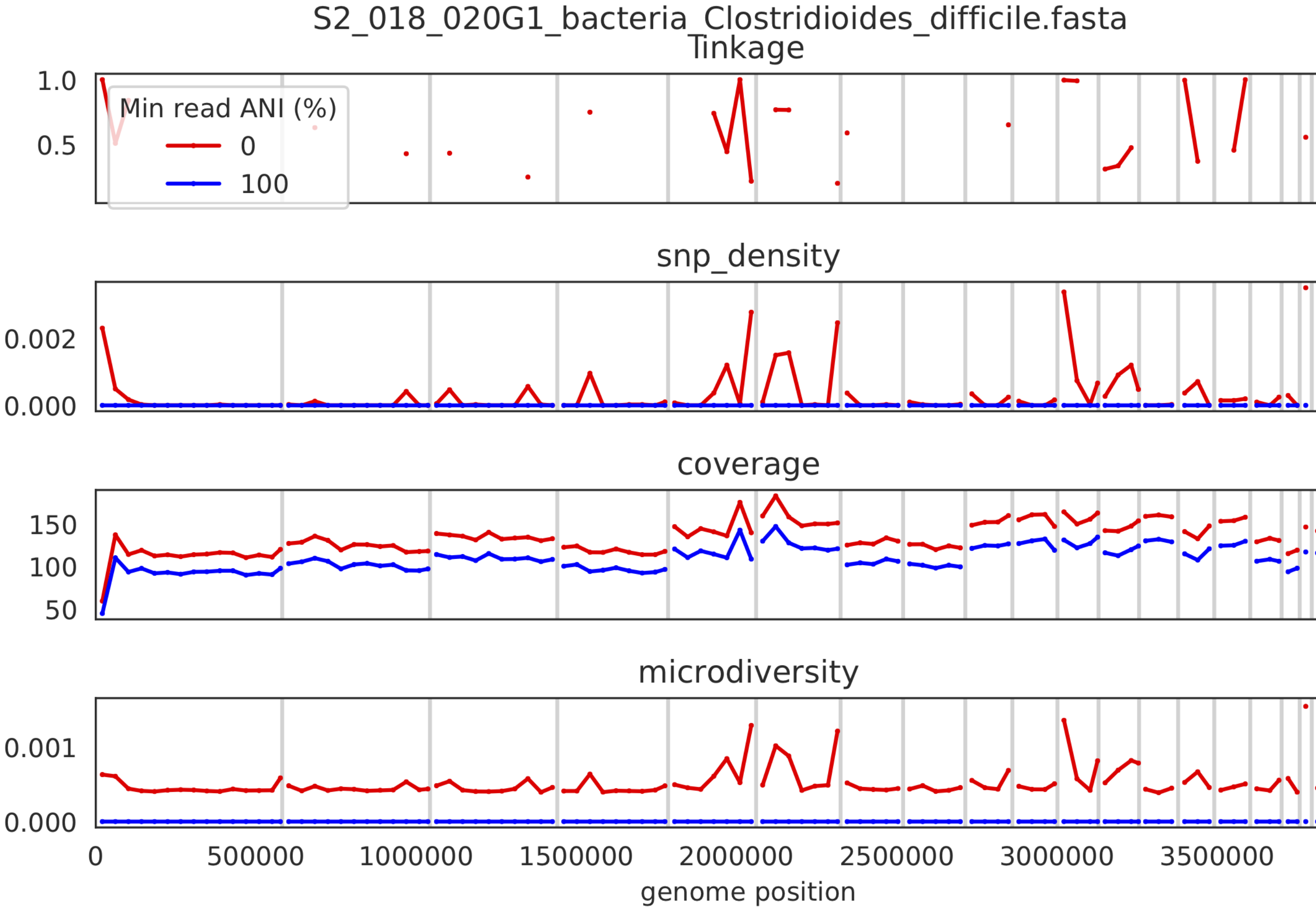
S2_018_020G1_bacteria_Clostridioides_difficile.fasta
linkage
1.0
0.5
Min read ANI (%)
0
100
snp_density
0.002
0.000
coverage
150
100
50
microdiversity
0.001
0.000
0
500000
1000000
1500000
2000000
2500000
3000000
3500000
genome position

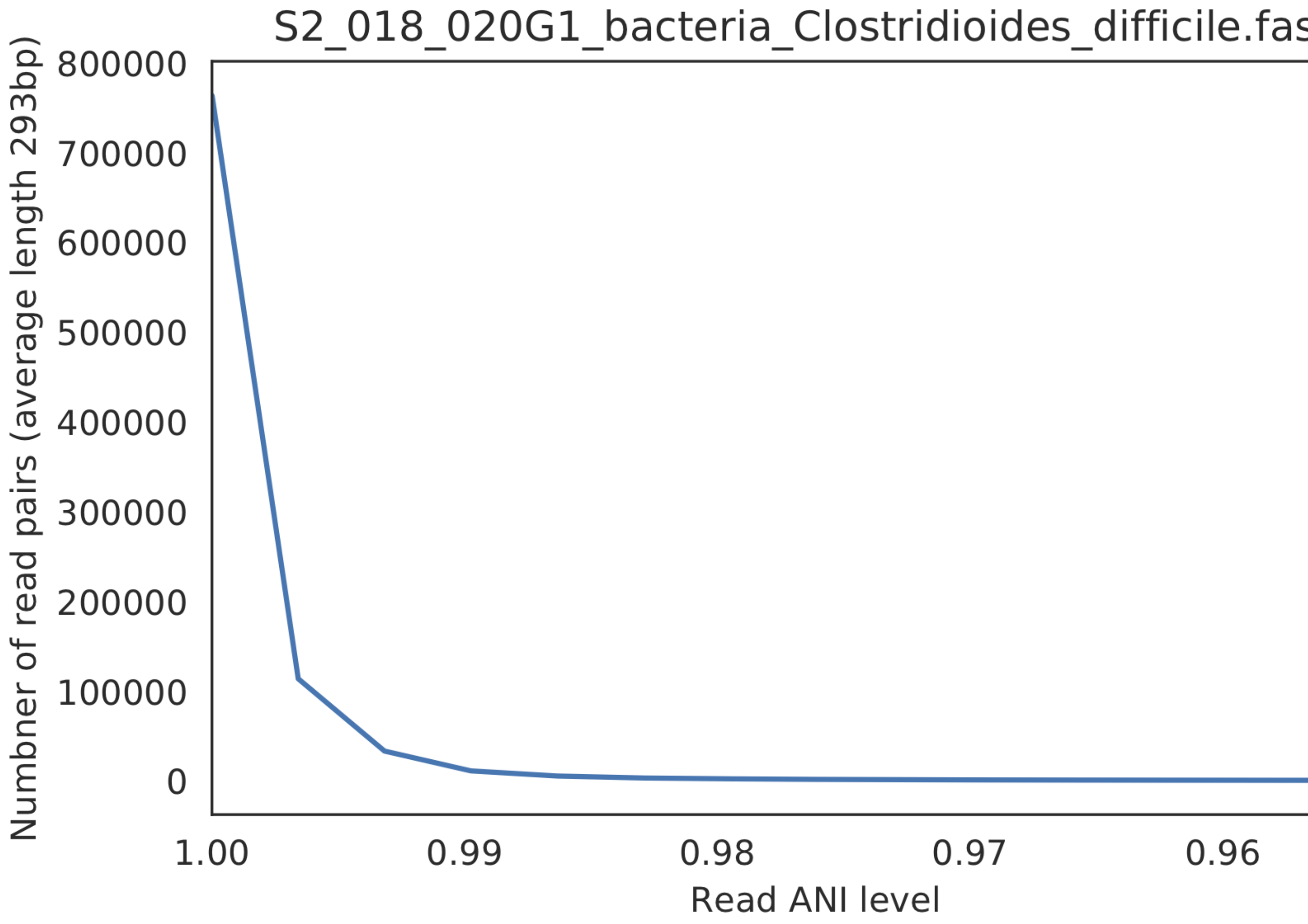
S2_018_020G1_bacteria_Clostridioides_difficile.fas
Numbner of read pairs (average length 293bp)
800000
700000
600000
500000
400000
300000
200000
100000
0
1.00
0.99
0.98
0.97
0.96
Read ANI level

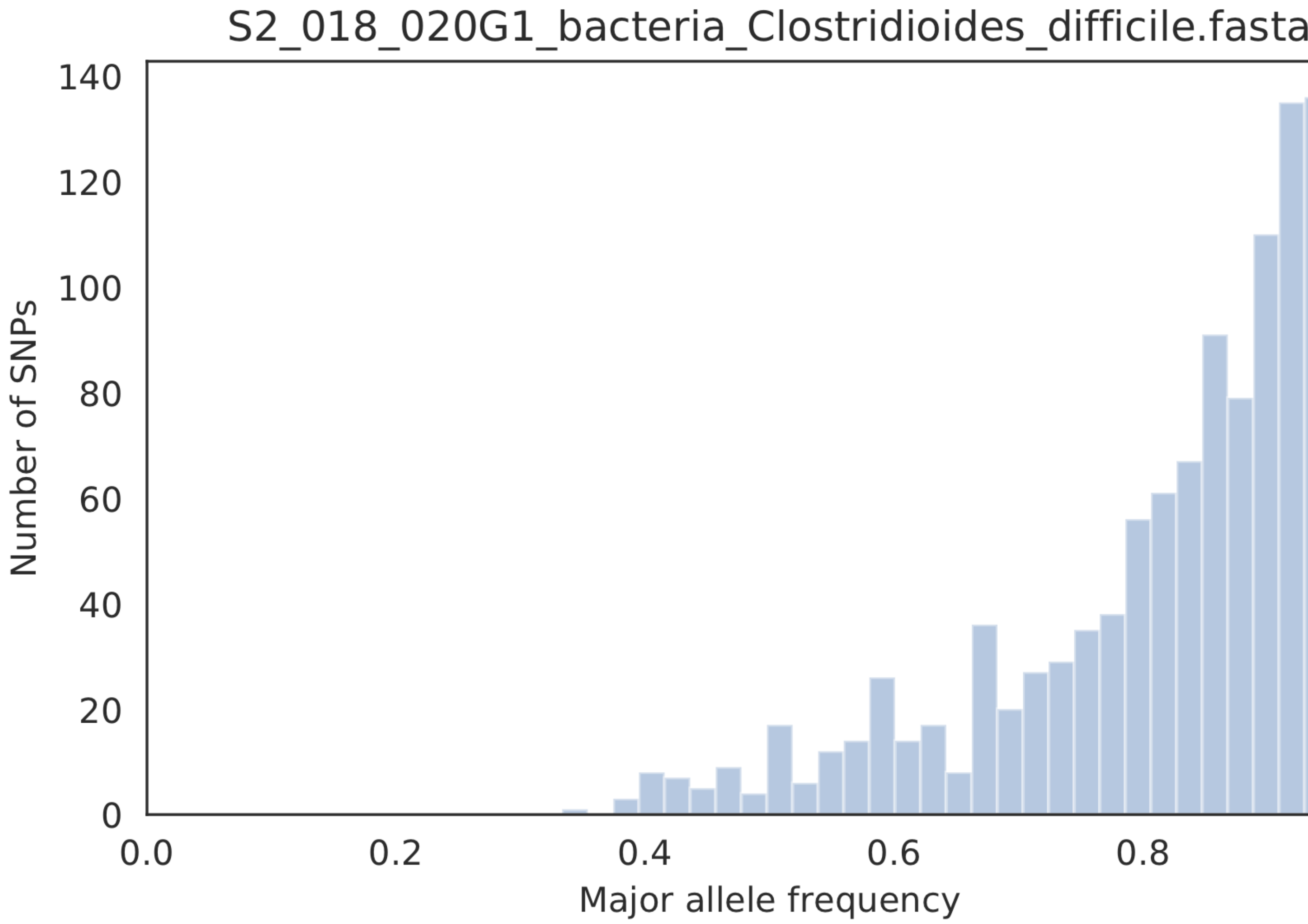
S2_018_020G1_bacteria_Clostridioides_difficile.fasta
Number of SNPs
0
20
40
60
80
100
120
140
0.0
0.2
0.4
0.6
0.8
Major allele frequency

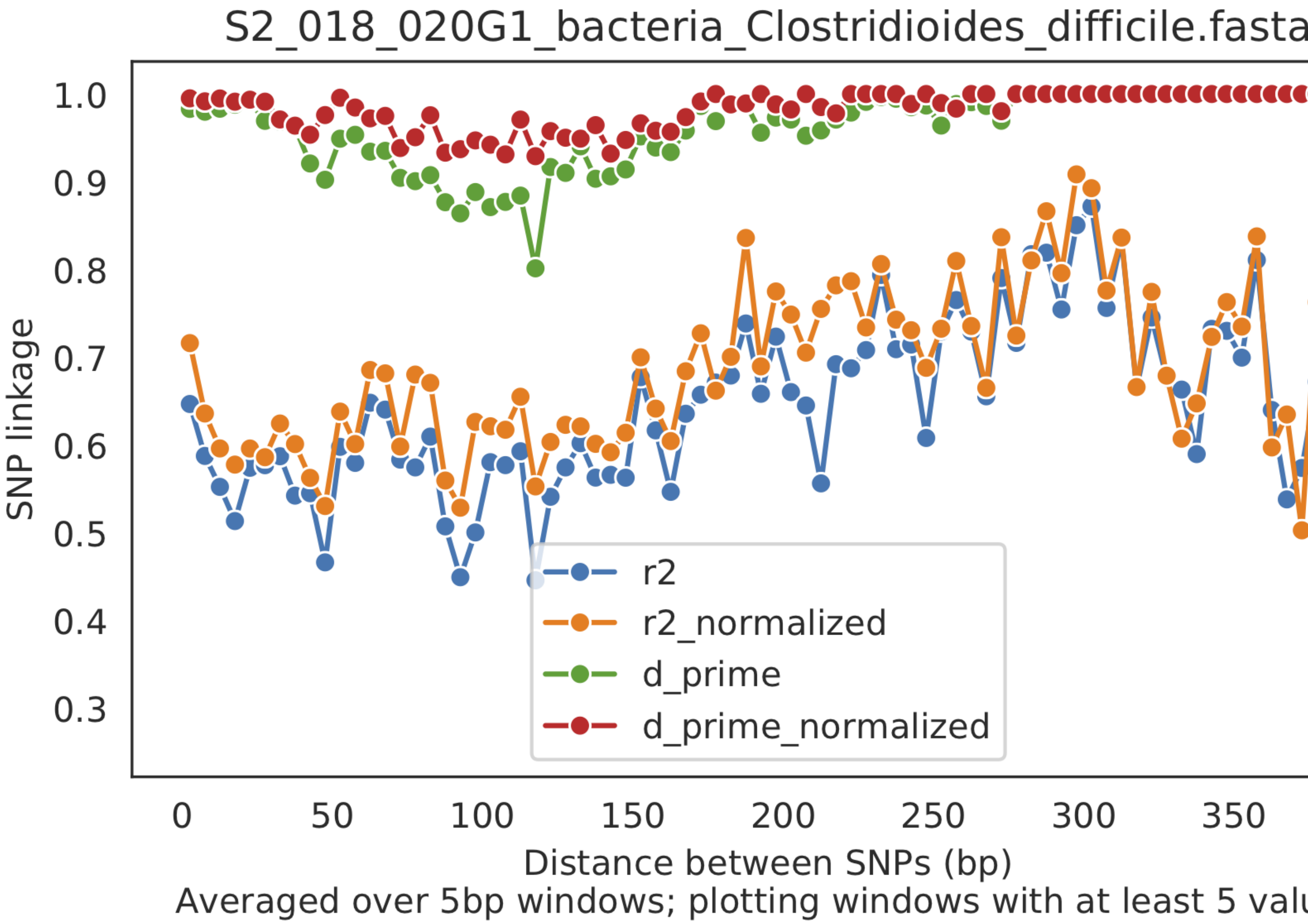
S2_018_020G1_bacteria_Clostridioides_difficile.fasta
SNP linkage
1.0
0.9
0.8
0.7
0.6
0.5
0.4
0.3
0
50
100
150
200
250
300
350
Distance between SNPs (bp)
Averaged over 5bp windows; plotting windows with at least 5 valu
r2
r2_normalized
d_prime
d_prime_normalized

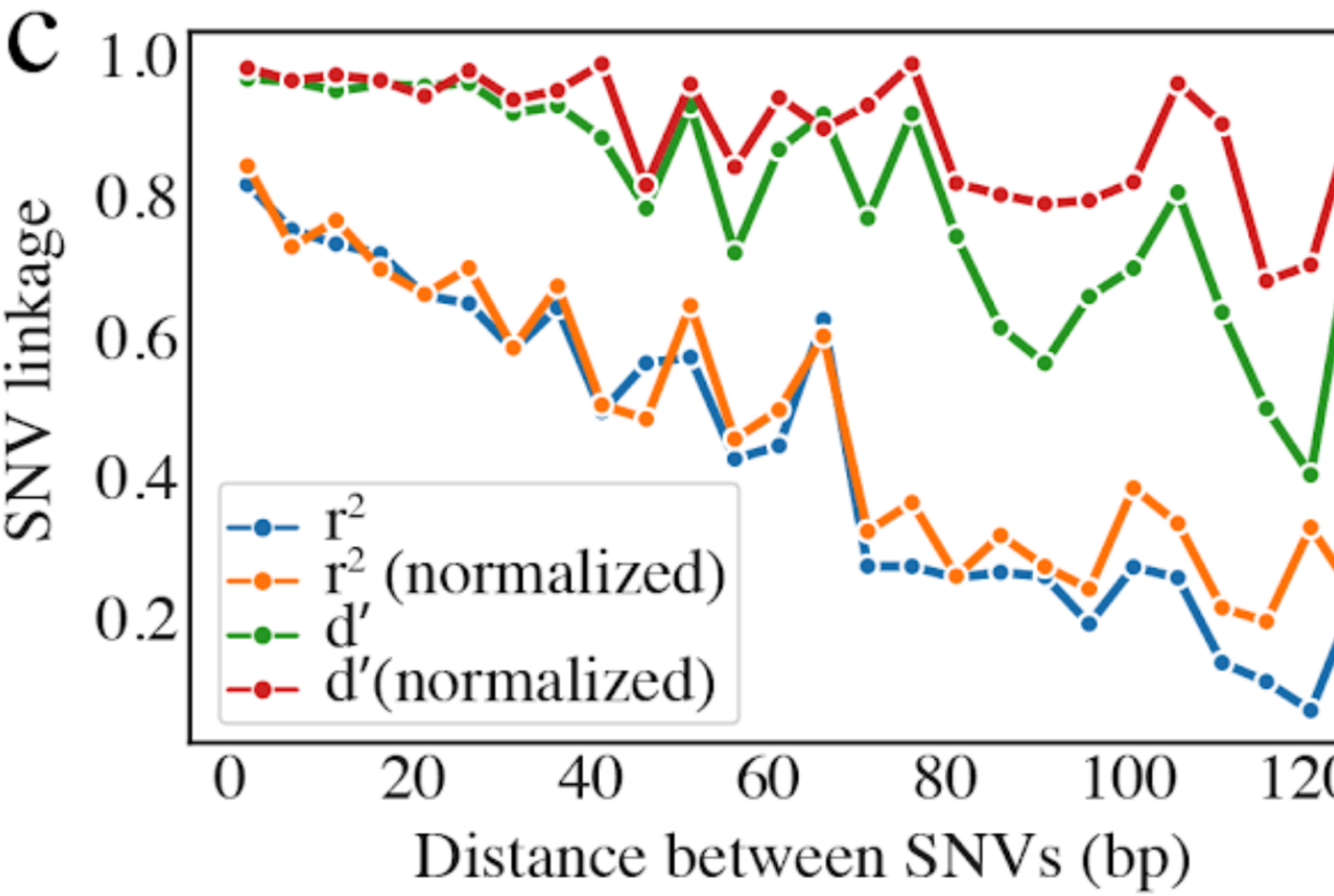
C
SNV linkage
1.0
0.8
0.6
0.4
0.2
r2
r2 (normalized)
d′
d′(normalized)
0
20
40
60
80
100
Distance between SNVs (bp)

###### 5) Linkage decay

Metrics of SNV linkage vs. distance between SNVs; linkage decay (shown in one plot and not the other) is a common signal of recombination.

###### 6) Read filtering plots

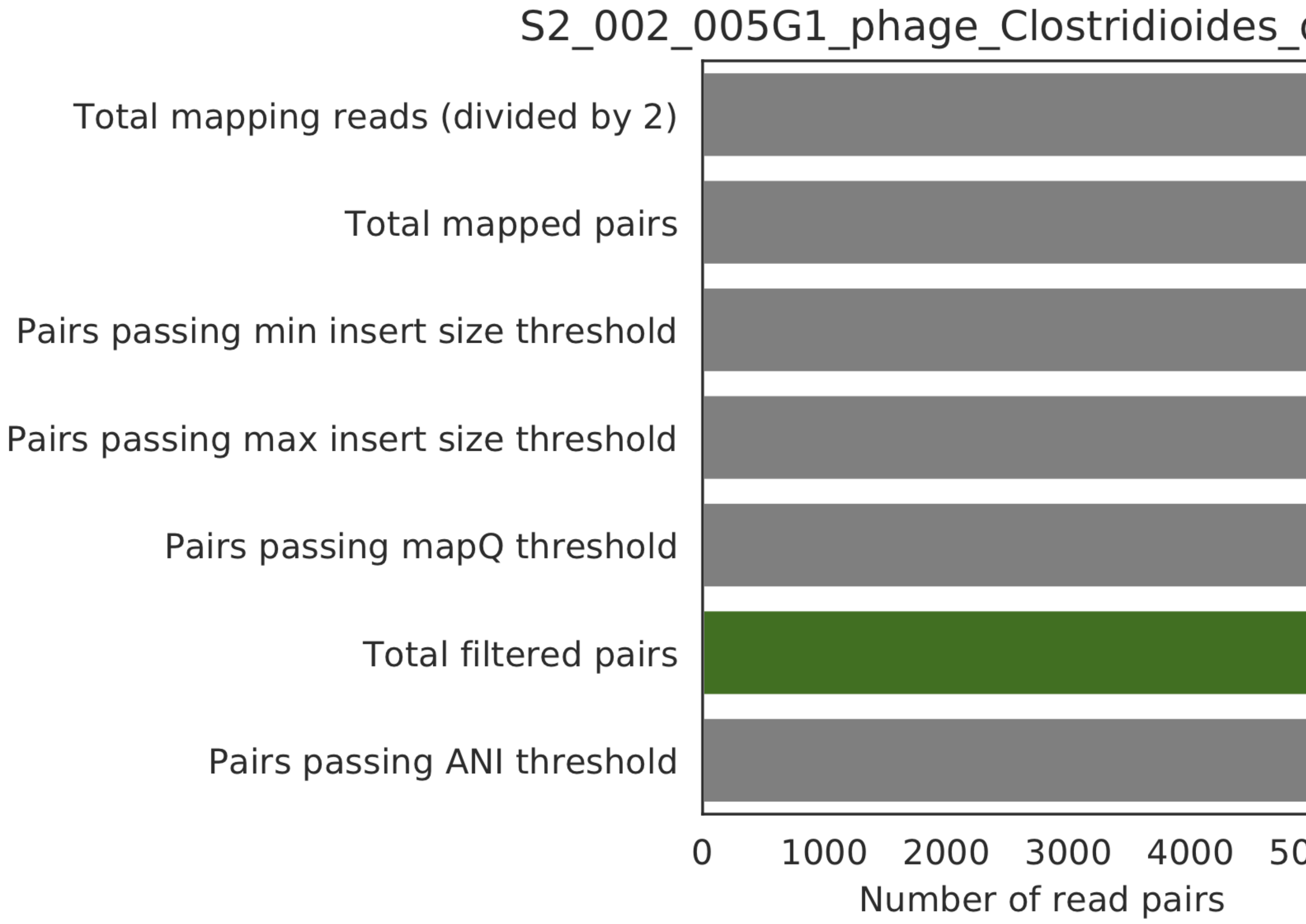

Bar plots showing how many reads got filtered out during filtering. All percentages are based on the number of paired reads; for an idea of how many reads were filtered out for being non-paired, compare the top bar and the second to top bar.

###### 7) Scaffold inspection plot (large)

This is an elongated version of the genome-wide microdiversity metrics that is long enough for you to read scaffold names on the y-axis

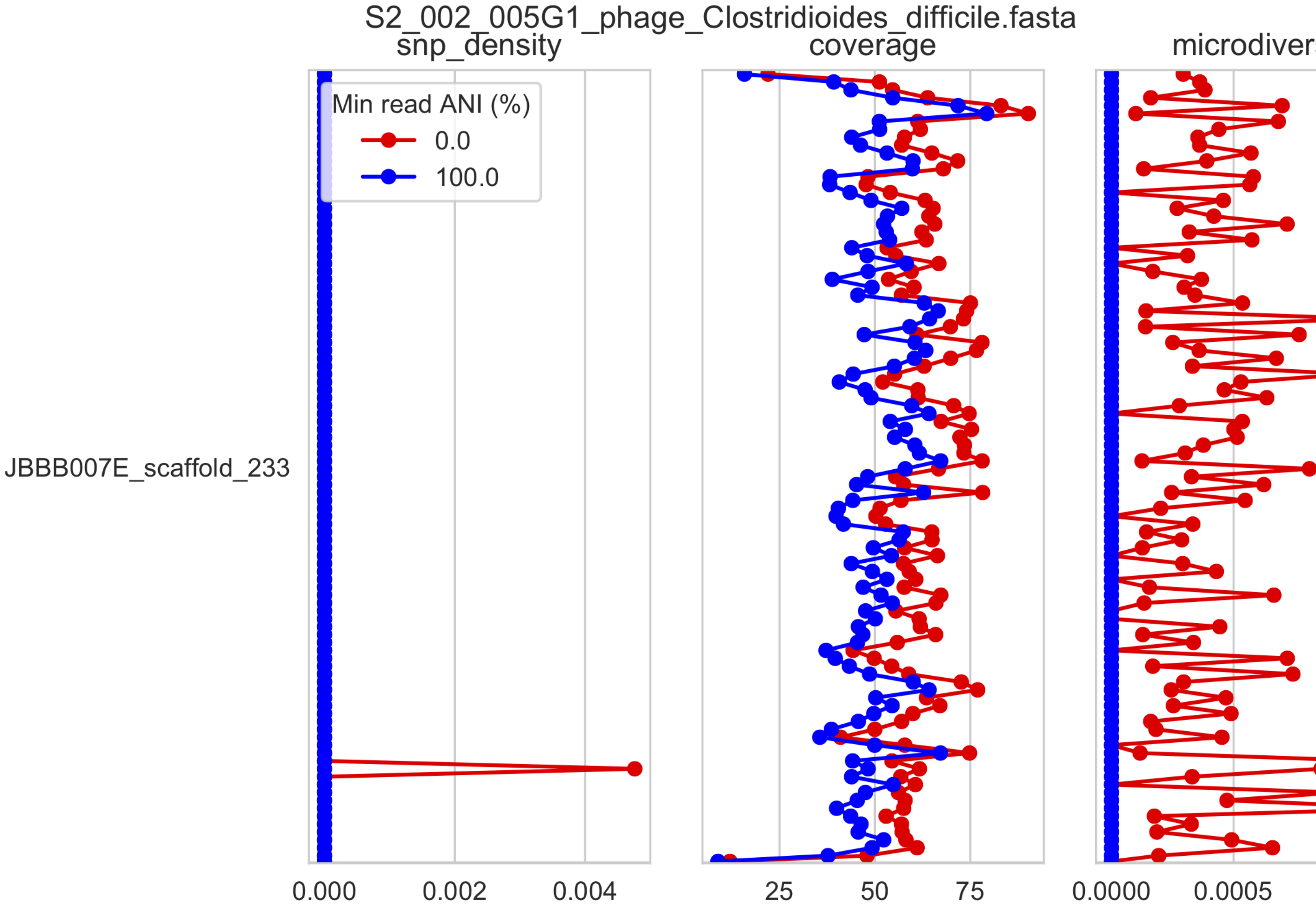
S2_002_005G1_phage_Clostridioides_difficile.fasta
snp_density
coverage
microdivers
Min read ANI (%)
0.0
100.0
JBBB007E_scaffold_233
0.000
0.002
0.004
25
50
75
0.0000
0.0005

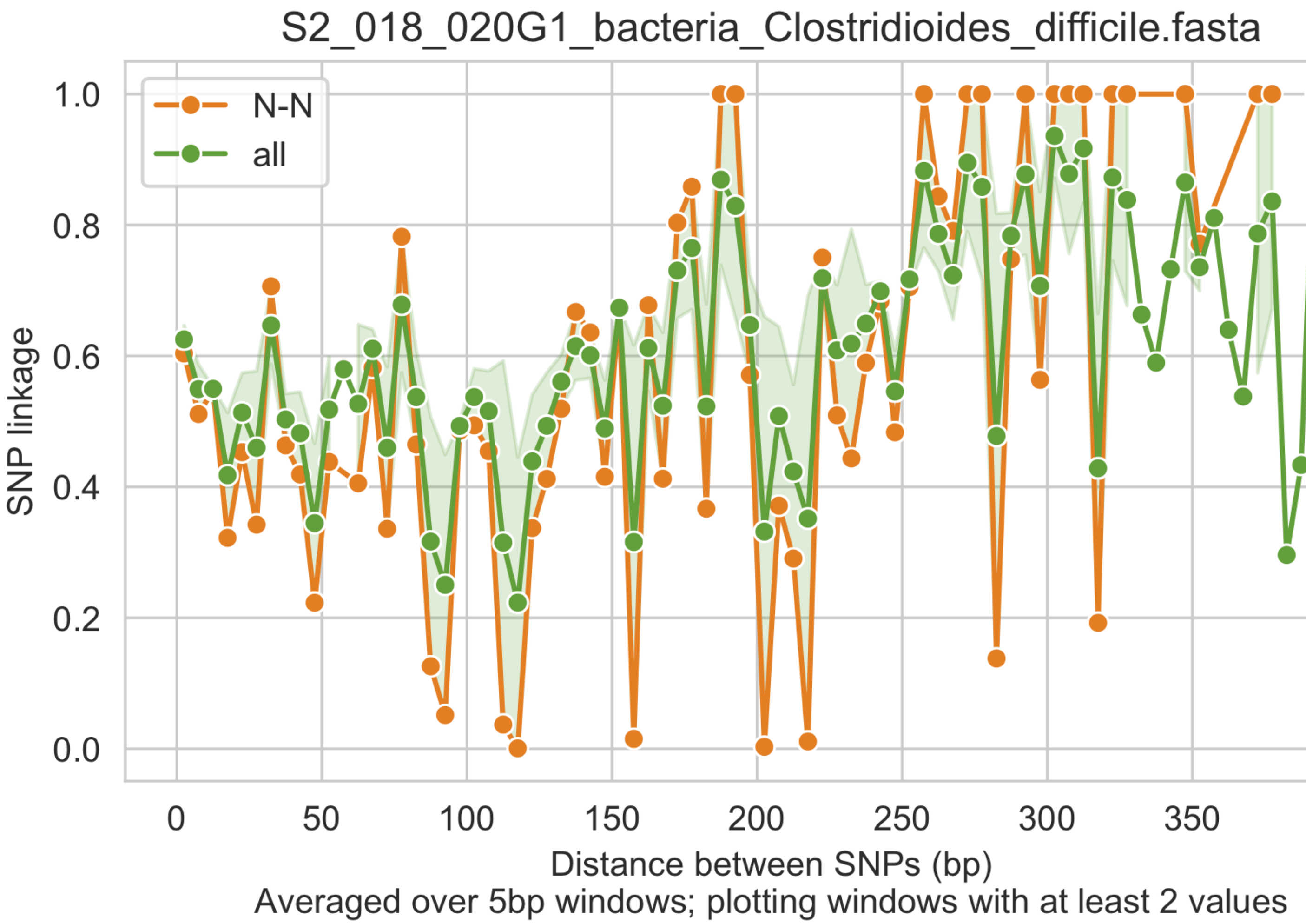
S2_018_020G1_bacteria_Clostridioides_difficile.fasta
N-N
all
SNP linkage
1.0
0.8
0.6
0.4
0.2
0.0
0
50
100
150
200
250
300
350
Distance between SNPs (bp)
Averaged over 5bp windows; plotting windows with at least 2 values

###### 8) Linkage with SNP type (GENES REQUIRED)

Linkage plot for pairs of non-synonymous SNPs and all pairs of SNPs

###### 9) Gene histograms (GENES REQUIRED)

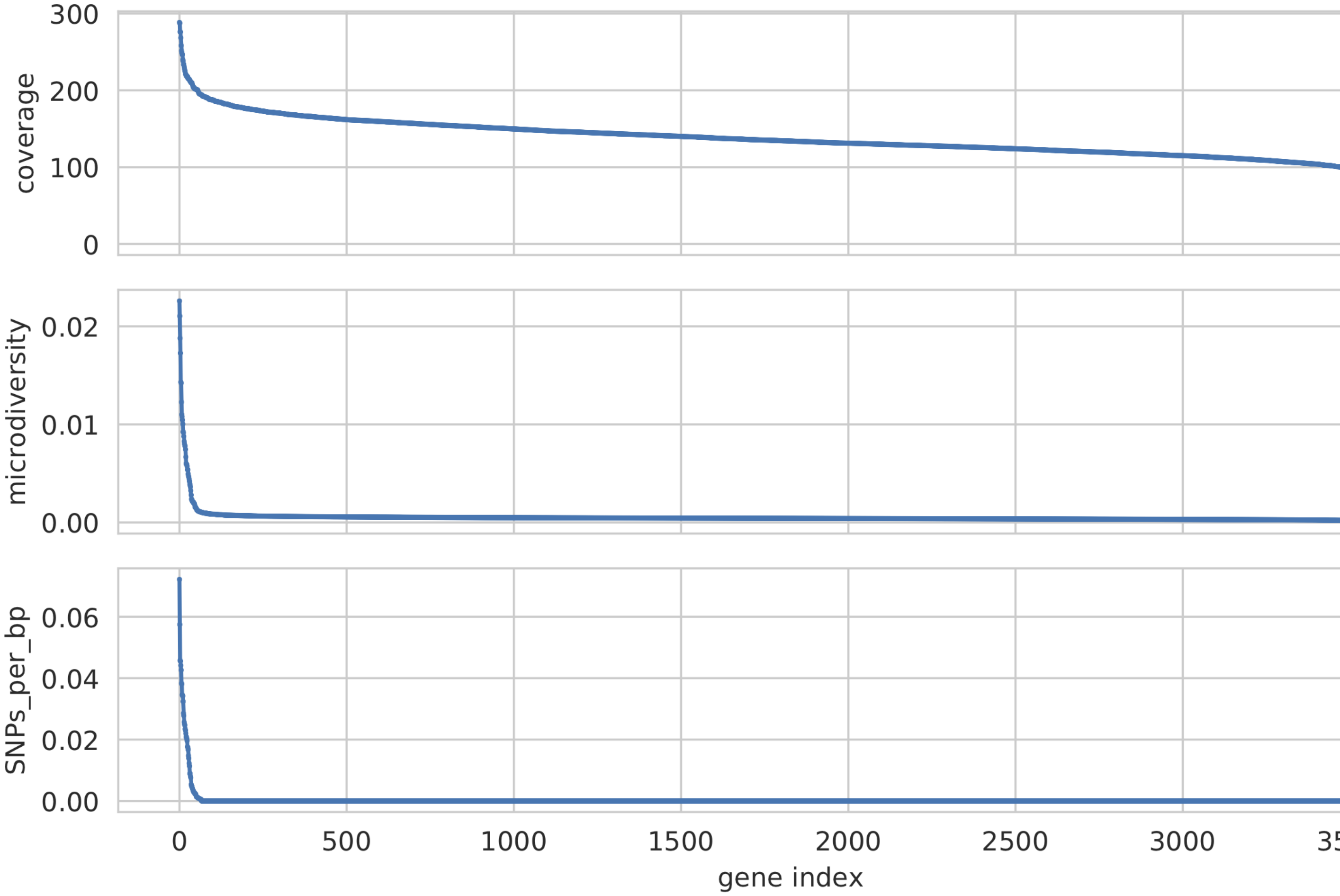

Histogram of values for all genes profiled

###### 10) Compare dendrograms (RUN ON COMPARE; NOT PROFILE)

A dendrogram comparing all samples based on popANI and based on shared_bases.

#### 1.5 Tutorial

The following tutorial goes through an example run of inStrain. You can follow along with your own data, or download the files that are used in this tutorial from the following link. Note that the reads files are several Gbp, and you really only need one set of reads (forward, which end in .1.fastq.gz, and reverse, which end in .2.fastq.gz). These read come from fecal samples of premature infants.

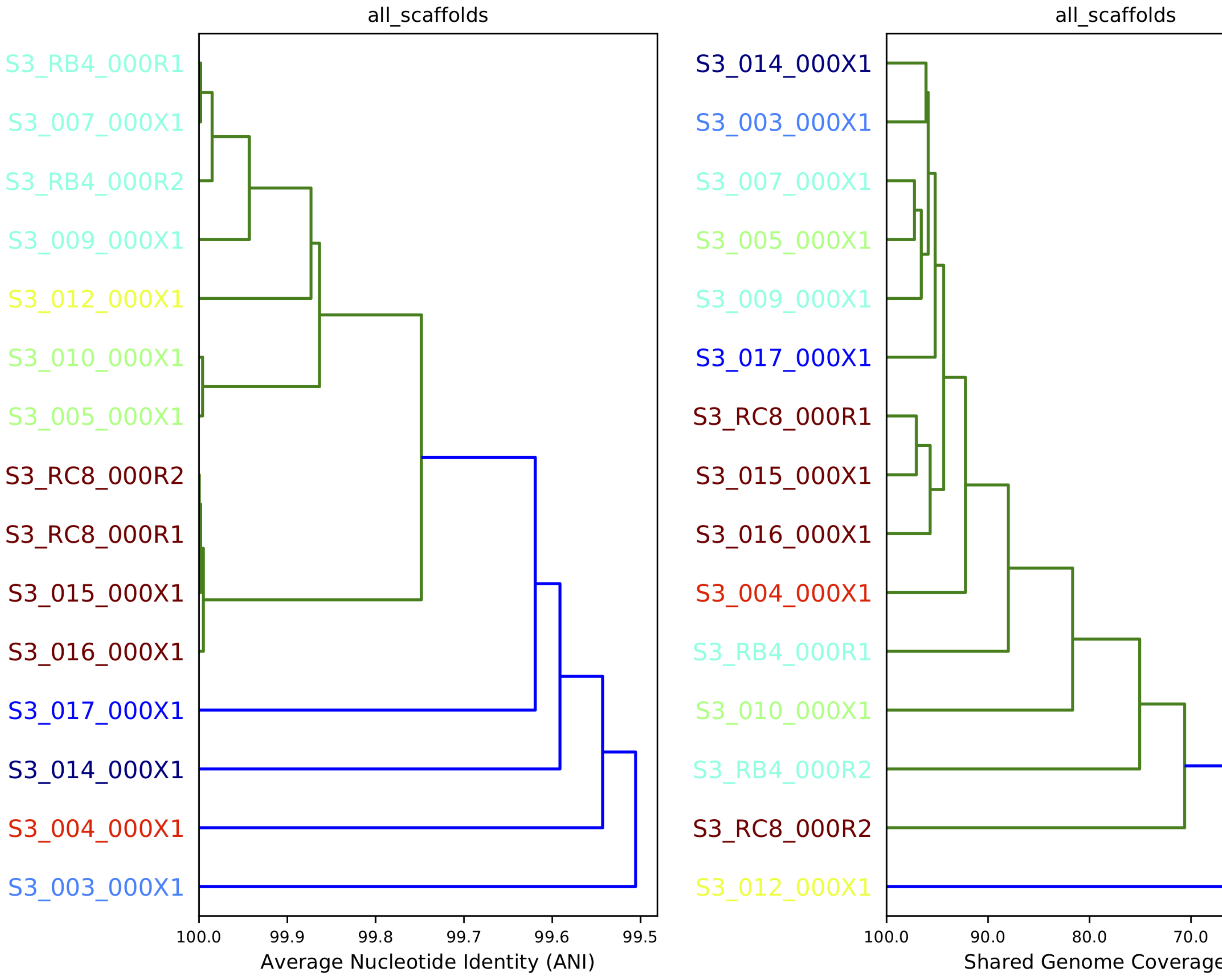
all_scaffolds
S3_RB4_000R1
S3_007_000X1
S3_RB4_000R2
S3_009_000X1
S3_012_000X1
S3_010_000X1
S3_005_000X1
S3_RC8_000R2
S3_RC8_000R1
S3_015_000X1
S3_016_000X1
S3_017_000X1
S3_014_000X1
S3_004_000X1
S3_003_000X1
100.0 99.9 99.8 99.7 99.6 99.5
Average Nucleotide Identity (ANI)
all_scaffolds
S3_014_000X1
S3_003_000X1
S3_007_000X1
S3_005_000X1
S3_009_000X1
S3_017_000X1
S3_RC8_000R1
S3_015_000X1
S3_016_000X1
S3_004_000X1
S3_RB4_000R1
S3_010_000X1
S3_RB4_000R2
S3_RC8_000R2
S3_012_000X1
100.0 90.0 80.0 70.0
Shared Genome Coverage

**See also:**

**quickstart** To get started using the program

*Program documentation* For descriptions of what the modules can do and information on how to prepare data for inStrain

*Example output and explanations* To view example output

*Advanced use* For detailed information on how to rationally adjust inStrain parameters

##### 1.5.1 Preparing .bam and .fasta files

After downloading the genome file that you would like to profile (.fasta file) and at least one set of paired reads, the first thing to do is to map the reads to the .fasta file in order to generate a .bam file.

When this mapping is performed it is important that you map to all genomes simutaneously, so the first thing to do is to combine all of the genomes that you'd like to map into a single .fasta file:

```
$  cat raw_data/S2_002_005G1_phage_Clostridioides_difficile.fasta raw_data/S2_018_
↪020G1_bacteria_Clostridioides_difficile.fasta > allGenomes_v1.fasta
```

Next we must map all of the reads to this .fasta file in order to create .bam files. In this tutorial we will use the mapper Bowtie 2, and also the program shrinksam to make the resulting .sam files smaller. In this case we will just make a .sam file and let inStrain handle conversion to .bam format

```
$ mkdir bt2

$ bowtie2-build allGenomes_v1.fasta bt2/allGenomes_v1.fasta.fa

$ bowtie2 -p 6 -x bt2/allGenomes_v1.fasta.fa -1 raw_data/N4_005_026G1.r1.fastq.gz -2␣
↪raw_data/N4_005_026G1.r2.fastq.gz 2> N4_005_026G1_mapped.log | shrinksam >␣
↪allGenomes_v1.fasta-vs-N4_005_026G1.sam
```

##### 1.5.2 Running inStrain profile

Now that we've made a mapping file we can run inStrain to profile the microdiversity within this genome population.

**Note:** It is possible to run make of the next steps simultaneously with this one, including profile_genes, genome_wide, and plot, just by including more parameters to this initial profile step, but in this tutorial we're going to run these steps separately

To run inStrain profile you just need to provide the .sam or .bam file, the .fasta file that the reads are mapped to, and the name of the output folder:

```
$ inStrain profile allGenomes_v1.fasta-vs-N4_005_026G1.sam allGenomes_v1.fasta -o␣
↪allGenomes_v1.fasta-vs-N4_005_026G1.IS
```

##### 1.5.3 Running inStrain genome_wide

Now that we've profiled all scaffolds, its time to average the results for scaffolds that belong to the same genome. This is done using the genome_wide command

```
$ inStrain genome_wide -i allGenomes_v1.fasta-vs-N4_005_026G1.IS -s raw_data/*.fasta
```

##### 1.5.4 Running profile_genes

In order to get gene level information, we need to provide inStrain with a list of the genes to profile. We can call these genes using the program Prodigal:

```
$ prodigal -i raw_data/S2_002_005G1_phage_Clostridioides_difficile.fasta -d S2_002_
↪005G1_phage_Clostridioides_difficile.fasta.genes.fna

$ prodigal -i raw_data/S2_018_020G1_bacteria_Clostridioides_difficile.fasta -d S2_018_
↪020G1_bacteria_Clostridioides_difficile.fasta.genes.fna

$ cat S2_002_005G1_phage_Clostridioides_difficile.fasta.genes.fna S2_018_020G1_
↪bacteria_Clostridioides_difficile.fasta.genes.fna > allGenomes_v1.genes.fna
```

Once we have all the genes to profile in .fna format, we can tell inStrain to profile them

$ inStrain profile_genes -i allGenomes_v1.fasta-vs-N4_005_026G1.IS/ -g allGenomes_v1.genes.fna

##### 1.5.5 Plotting

To make all of the plots that you can given the current inStrain profile object, just run the plot command:

```
inStrain plot -i allGenomes_v1.fasta-vs-N4_005_026G1.IS
```

##### 1.5.6 inStrain compare

To run inStrain compare, you first need to profile another .bam file from another set of reads based on mapping to the same .fasta file. Once that is done, you can compare them using the command:

```
inStrain compare -i allGenomes_v1.fasta-vs-N4_005_026G1.IS allGenomes_v1.fasta-vs-N5_
↪215_032G1.IS -o allGenomes_v1.fasta.RC
```

##### 1.5.7 Interpreting the output

For help interpreting the output, see *Example output and explanations*

#### 1.6 Advanced use

##### 1.6.1 Adjusting parameters

There are a number of important considerations when running inStrain. Here is some theory and data about how to make inStrain work best

##### Reference genome selection

inStrain relies on mapping reads from a sample to a reference genome. How similar the reference genome is to the reads, and the minimum read ANI threshold that you set, are very important and will determine much of what you get out of inStrain.

Below are a series of plots made by introducing a known number of mutations into an E. coli genome, simulating reads from these mutated genomes (at 20x coverage) with known ANI differences from the original reference genome, mapping the synthetic reads back to the original reference genome, and running inStrain.

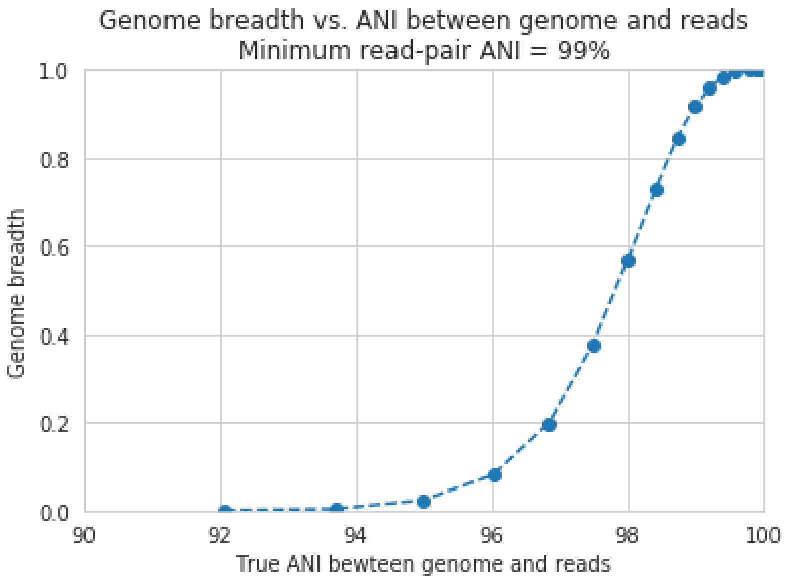

In the above plot, inStrain was run with a minimum read ANI of 0.99 (inStrain profile parameter *-l* or *–filter_cutoff*). The reported genome breadth is reported on the y-axis. At 20x coverage, you should see 100% genome breadth (meaning that every base of the reference genome is covered by at least one read). However, when the reference genome is sufficiently different from the reads, the breadth is much lower. This is because when the read pair differs from the reference base by more than 99% ANI, it gets filtered out, and no longer maps to the genome. This can be exemplified a bit better by showing a variety of read filtering thresholds simultaneously:

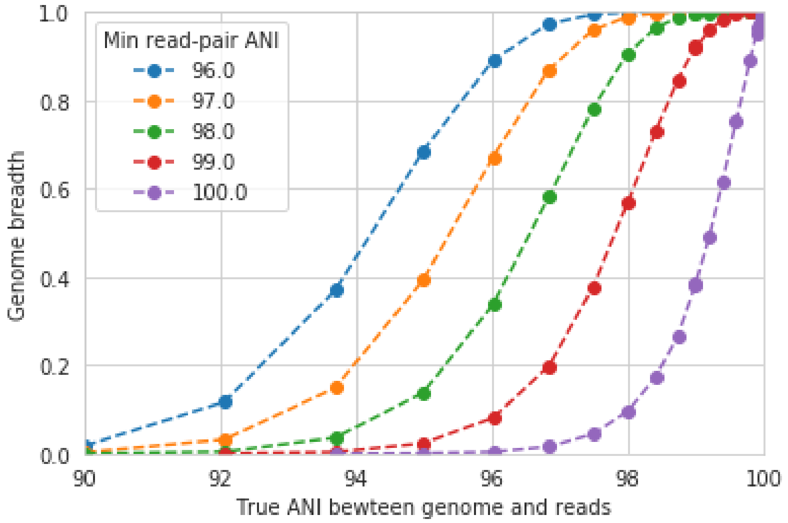

The line drawn in the first figure is now in red on this second figure. As you can see, the more you relax the minimum read ANI, the more you can align reads to more distantly related reference genomes.

> **Warning:** You don't want your minimum read pair ANI to be too relaxed, because then you risk mapping reads that don't actually belong to the population represented by your reference genome ("non-specific" mapping). You can also avoid non-specific mapping by increasing the size of your reference genome dataset (more on that below)

An important takeaway from the above figure is that the minimum read ANI should be at least 3% lower than the expected differences between your reads and the reference genome. If you look at the genome that's 96% ANI from the reads, for example, you see that none of the minimum read ANI levels get the correct breadth of 1. If you look at the genome that's 98% ANI from the reads, you can see that having a minimum read ANI of 96% is the only one that's actually near 100% breadth. This can also be visualized by looking at the distribution of ANI values of read pairs mapping to the 98% genome:

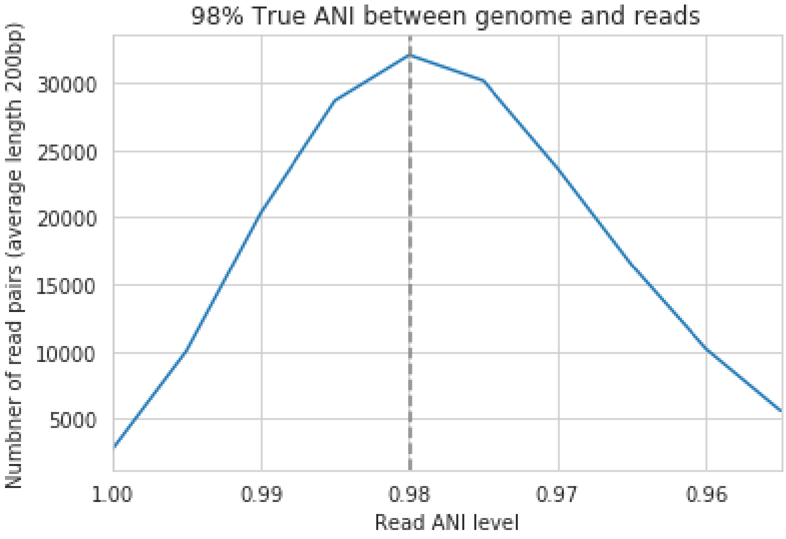

Most read pairs have 98%, as expected, but there is a wide distribution of read ANI values. This is because SNPs are not evenly spread along the genome, a fact that is even more true when you consider that real genomes likely have even more heterogeneity in where SNPs occur than this synthetic example.

The fact that the reads fail to map to heterogenous areas of the genome is also more problematic than it originally seems. It means that the area of the genome that are most similar to the sample reads will recruit reads during read mapping, but the (potentially interesting) areas with more SNPs will not. This is exemplified in the figure below:

The y-axis in this figure shows the inStrain calculated ANI; that is, the number of identified SNPs divided by the number of bases with at least 5x coverage. If you look at red line, where only reads with at least 99% ANI are mapped, the ANI of reads mapping to the genome is almost always overestimated. This is because reads are only mapping to a small fraction of the genome (see the breadth in the second figure), and the small fraction of the genome that the reads are mapping to are the regions with a small number of SNPs.

By staring at this figure like I have, you'll notice that the correct ANI is identified when the minimum read pair ANI is 2-3% lower than the actual difference between the reads and the genome. 96% minimum ANI reads correctly identify the ANI of the 98% genome, for example.

Finally, in case you're wondering what the maximum read ANI is that bowtie2 is table to map, the answer is that it's complicated:

When mapping to a genome that is 90% ANI to the reads, you no longer see a peak at 90% as you do in the 98% example. This is because bowtie2 doesn't have a string ANI cutoff, it just maps what it can. This likely depends on

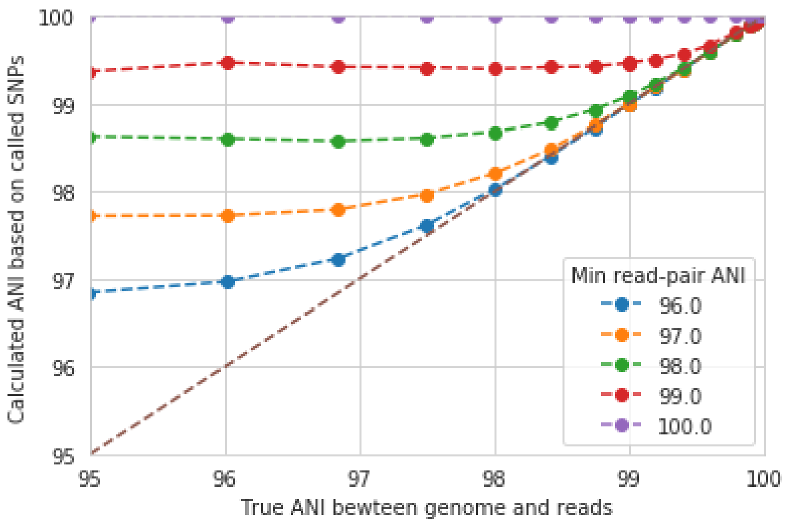
Calculated ANI based on called SNPs
True ANI bewteen genome and reads
Min read-pair ANI
96.0
97.0
98.0
99.0
100.0

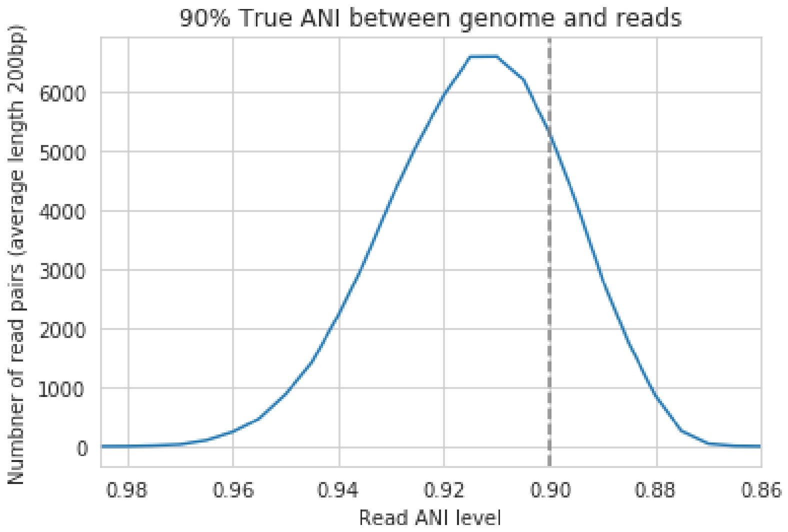
90% True ANI between genome and reads
Numbner of read pairs (average length 200bp)
Read ANI level

where the SNPs are along the read, whether they're in the seed sequence that bowtie2 uses, etc. While bowtie2 can map reads that are up to 86% ANI with the reference genome, I wouldn't push it past 92% based on this graph.

**Note:** In conclusion, you want your reference genome to be as similar to your reads as possible, and to set your minimum read-pair ANI to at least ~3% lower than the expected different from the reads and the reference genome. The inStrain default is 95% minimum read pair ANI, which is probably ideal in the case that you've assembled your reference genome from the sample itself. If you plan on using inStrain to map reads to a genome that you downloaded from a reference database, you may want to lower the minimum read-pair ANI to as low as ~92%, and ensure that the genome your mapping to is at least the same species as the organism in your reads (as genomes of the same species share ~95% ANI)

##### Mapping to multiple reference genomes

###### Mapping to multiple genomes simultaneously to avoid mis-mapping

There are a number of ways to avoid mis-mapped reads (reads from a different population mapping to your reference genome). One method is to filter out distantly related reads, including by using the minimum read-pair ANI threshold (*-l*, *–filter_cutoff*) or by using the mapQ score cutoff (more on that later). Another method is to include multiple reference genomes in the *.fasta* file that you map to, which gives the mapping software a chance to better place your reads.

When bowtie2 maps reads, by default, it only maps reads to a single location. That means that if a read maps at 98% ANI to one scaffold, and 99% ANI to another scaffold, it will place the read at the position with 99% ANI. If the read only maps to one scaffold at 98% ANI, however, bowtie2 will place the read there. Thus, by including more reference genome sequences when performing the mapping, reads will end up mapping more accurately overall.

**Based on the above information, if you'd like to run inStrain on multiple reference genomes for the same set of reads, you should concatenate the genomes first and map to the concatenated genome set. You can then use inStrain genome_wide to get information on each genome individually.**

**Note:** You can get can an idea of the extent of mis-mapping going on in your sample by looking at the variation in coverage across the genome. If you see a region of the genome with much higher coverage than the rest, it is likely that that region is recruiting reads from another population. Looking at these wavy coverage patterns can be confusing, however. Here is a link for more information on this phenomenon.

**Warning:** It is possible to include too many genomes in your reference .fasta file, however. You generally don't want to have genomes that are over 98% ANI to each other in your reference genome set, because then the genomes can steal reads from each other. More on that below.

###### Read stealing due to including closely related genomes in the reference .fasta file

If bowtie2 finds a read that maps equally well to multiple different positions in your .fasta file, it will randomly choose one of the two positions to place the read at. Because of this, you really don't want to have multiple positions in your .fasta file that are identical. At these positions it is impossible for the alignment algorithm to known which reference sequence the read should actually map to. You can then end up with "read stealing", where closely related genomes will steal reads from the true reference genome.

In the below example, thousands of bacterial genomes were dereplicated at 99.8% ANI and combined into a single .fasta file. One genome was randomly chosen to profile, and reads from the sample from which that genome was assembled were mapped to this concatenation of all genomes together and to that one genome individually. We then profiled the difference in read mapping when mapping to the two different .fasta files. Specifically, we looked at reads that mapped to the genome of interest when mapping to that genome individually, and mapped elsewhere when mapping to all genomes concatenated together.

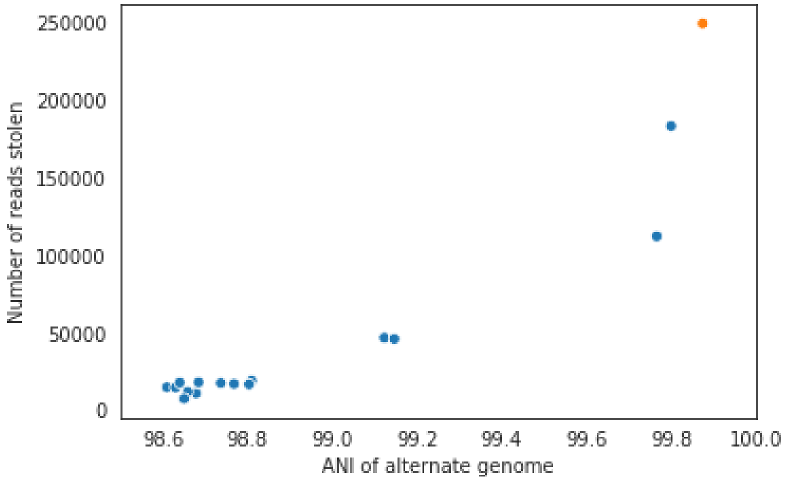

Each dot represents a genome in the concatenated genome set. The position on the x-axis indicates that genomes ANI to the genome of interest (orange dot), and the position on the y-axis indicates the number of reads that were stollen from the genome of interest. The number of reads that were stollen from the genome of interest is the number of reads that mapped to the genome of interest when it was mapped to as an individual .fasta file, but that now map to a different genome when reads were mapped to a concatenation of many genomes together.

As you can see, the more closely related an alternate genome is to a genome of interest, the more likely it is steal reads. This makes sense, because assuming that the genomes represented by blue dots are not actually present in the sample (likely true in this case), the only way these genomes have reads mapped to them is be having regions that are identical to the genome that is actually present in the sample. In fact, you can even calculate the probability of having an identical region as long as a pair of reads (190bp in this case) based on the genome ANI using the formula: Probability of identical 190bp fragment = (genome ANI) ^ 190. We can then overlay this onto the above plot:

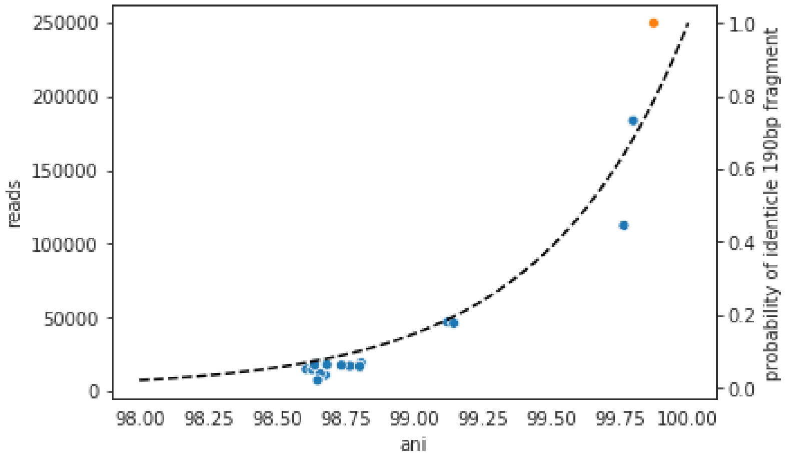

This simple formula fits the observed trend remarkably well, providing pretty good evidence that simple genome-ANI-based read stealing is what is going on.

**Note:** In the above example, read stealing approaches 0 at around 98% ANI. Thus, when dereplicating your genome set (using dRep for example), using a threshold of 98% or lower is a good idea.

As a final check, we can also filter reads by MapQ score. A MapQ is assigned to each read mapped by bowtie2, and is mean to signify how well the read mapped. MapQ scores are incredibly confusing (see the following link for more information), but MapQ scores of 0 and 1 have a special meaning. If a read maps equally well to multiple different locations on a .fasta file, it always gets a MapQ score of 0 or 1. Thus, by filtering out reads with MapQ scores < 2, we can see reads that map uniquely to one genome only.

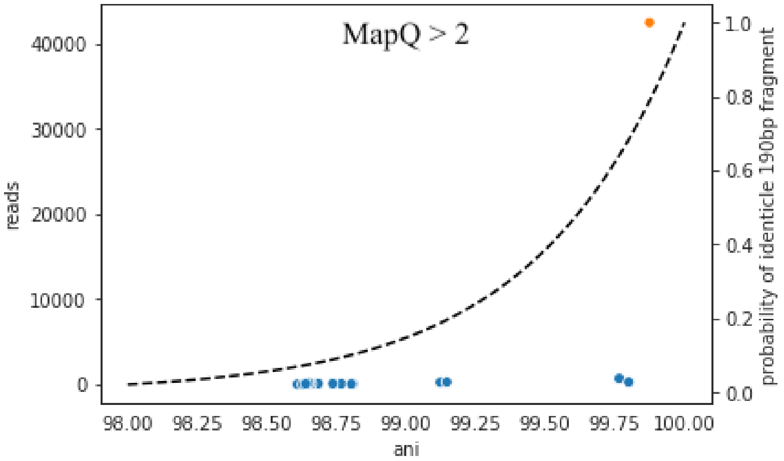

Just as we suspected, read no longer map to these alternate genomes at all. This provides near conclusive evidence that the organisms with these genomes are not truly in the sample, but are merely stealing reads from the genome of the organisms that is there by having regions of identical DNA. For this reason it can be smart to set a minimum MapQ score of 2 to avoid mis-mapping, but at the same time, look at the difference in the number of reads mapping to the correct genome when the MapQ filter is used- 85% of the reads are filtered out. Using MapQ filters is a matter of debate depending on your specific use-case.

##### Other considerations

A final aspect to consider is de novo genome assembly. When multiple closely related genomes are present in a sample, the assembly algorithm can break and you can fail to recover genomes from either organism. A solution to this problem is to assemble and bin genomes from each metagenomic sample individually, and dereplicate the genome set at the end. For more information on this, see the publication "dRep: a tool for fast and accurate genomic comparisons that enables improved genome recovery from metagenomes through de-replication"

Assuming you de-replicate your genomes at 98% before mapping to run inStrain, another matter to consider is how you define detection of a genome in a sample. The following figure shows the expected genome overlap between genomes of various ANI values from different environments (adapted from "Consistent metagenome-derived metrics verify and define bacterial species boundaries")

As you can see, genomes from that share >95% ANI tend to share ~75% of their genome content. Thus, using a breadth detection cutoff of somewhere around 50-75% seems to be reasonable.

**Note:** Based on the above information we recommend the following pipeline. 1) Assemble and bin genomes from all samples individually. 2) Dereplicate genomes based on 97-98% ANI. 3) Concatenate all dereplicated genomes into a single .fasta file, and map reads from all original samples to this concatenated .fasta file. 4) Use inStrain to profile the strain-level diversity of each microbial population (represented by a genome in your concatenated .fasta file)

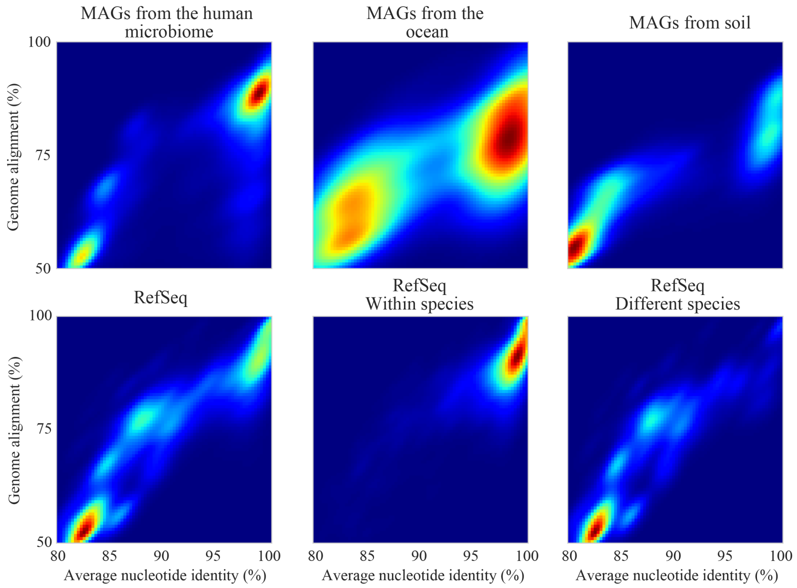

##### Detecting closely related organisms with inStrain compare

To compare strains with inStrain, one must first generate two inStrain profiles (using the command *inStrain profile*) based on mapping reads to the same .fasta file. *inStrain compare* then compares the reads mapped from both samples to the same .fasta file to calculate an extremely precise and accurate ANI value for the populations in the two samples. In order for this to work well, however, there are a number of things that you must keep in mind.

Same as *inStrain profile*, *inStrain compare* requires the user to think about the minimum read-pair ANI that should be considered. It will use the read-pair ANI selected during the *inStrain profile* commands by default, but the user can also access many other min read-pair ANI values using the ANI (see section *Dealing with "mm"* below for more information)

Below are a series of plots generated from synthetic data. In these plots, a reference genome was downloaded from NCBI and mutated to a series of known ANI values. Synthetic reads were generated from each of these mutated genomes, mapped back to the original genome, and then *inStrain profile* was run on the resulting .bam file. Synthetic reads were also generated from the original genome and mapped back to it as well. Finally, *inStrain compare* was run to compare the .bams resulting the mutated genomes to the original genome. This allows us to compare the (pop)ANI value reported by inStrain compare to the true ANI value (generated by introducing a known number of mutations).

---

**Note:** The ANI values reported from inStrain compare are referred to as popANI values

---

As you can see, the calculated popANI value is incorrect when the actual ANI different is large. This makes sense based on the section above. When mapping reads from an organism that is 90% ANI to the .fasta file that you're mapping to, many read-pairs will have an ANI of over 90%, and thus be thrown out when using a 95% read-pair ANI cutoff. This can also be exemplified by looking at the fraction of the genome that is compared when comparing genomes of increasing ANI.

As expected, when comparing genomes of low ANI values with a read-pair ANI threshold of 95%, only a small amount of the genome is actually being compared. This genome fraction represents the spaces of the genome that happen to be the most similar, and thus the inStrain calculated ANI value is overestimated. It's also worth noting that when comparing genomes 95% ANI away from each other, only 50% of the genome bases can be compared when you filter read-pairs at a minimum of 95% ANI. You can also visualize how a lack of genome breadth of coverage leads to errors in the ANI calculation in another way:

Now that we understand all of this, lets visualize lots of minimum read-pair ANI cutoffs simultaneously

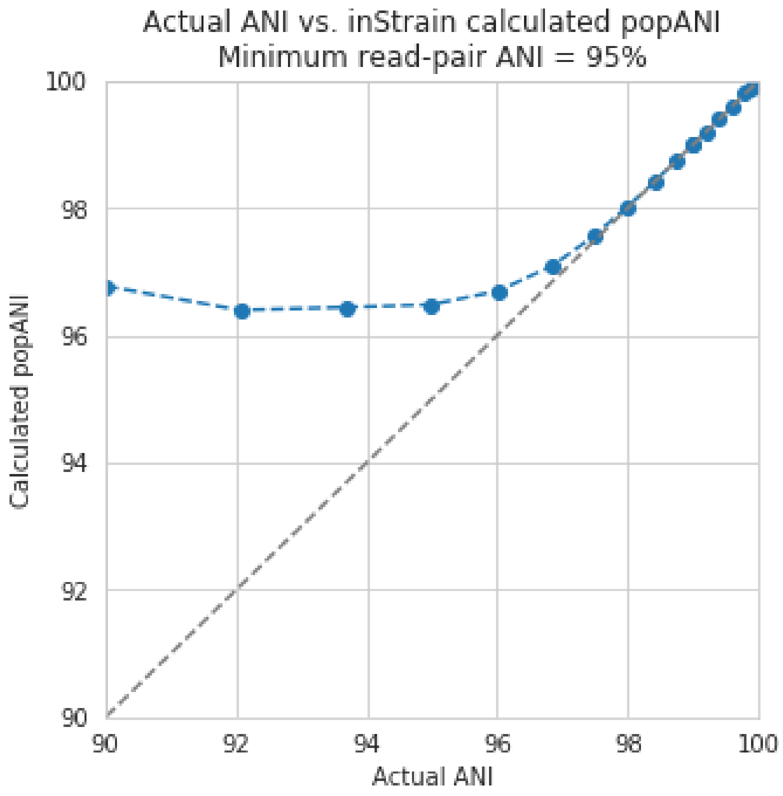
Actual ANI vs. inStrain calculated popANI
Minimum read-pair ANI = 95%
Calculated popANI
Actual ANI
100
98
96
94
92
90
90
92
94
96
98
100

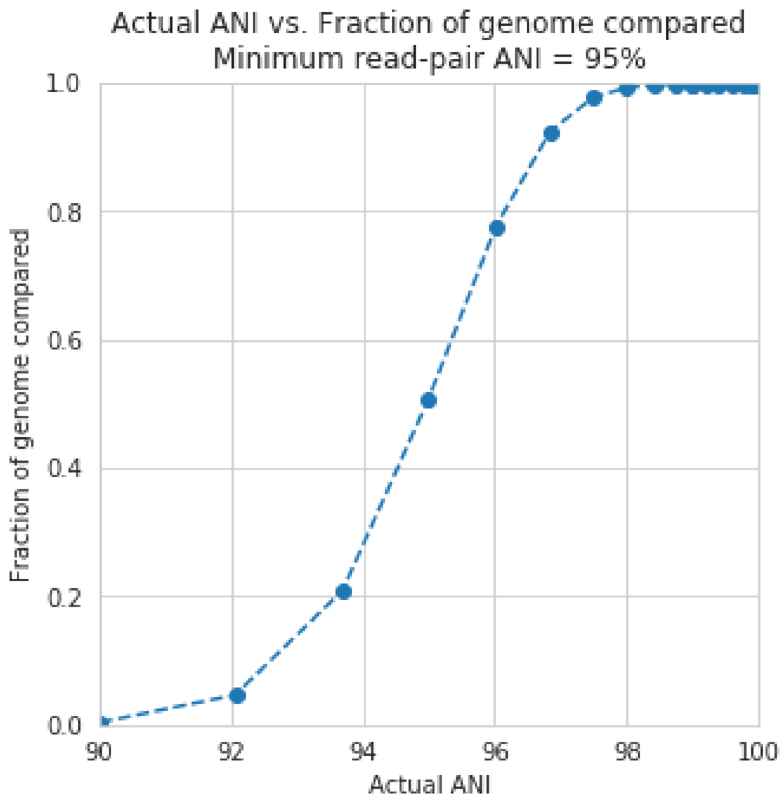
Actual ANI vs. Fraction of genome compared
Minimum read-pair ANI = 95%
Fraction of genome compared
Actual ANI
1.0
0.8
0.6
0.4
0.2
0.0
90
92
94
96
98
100

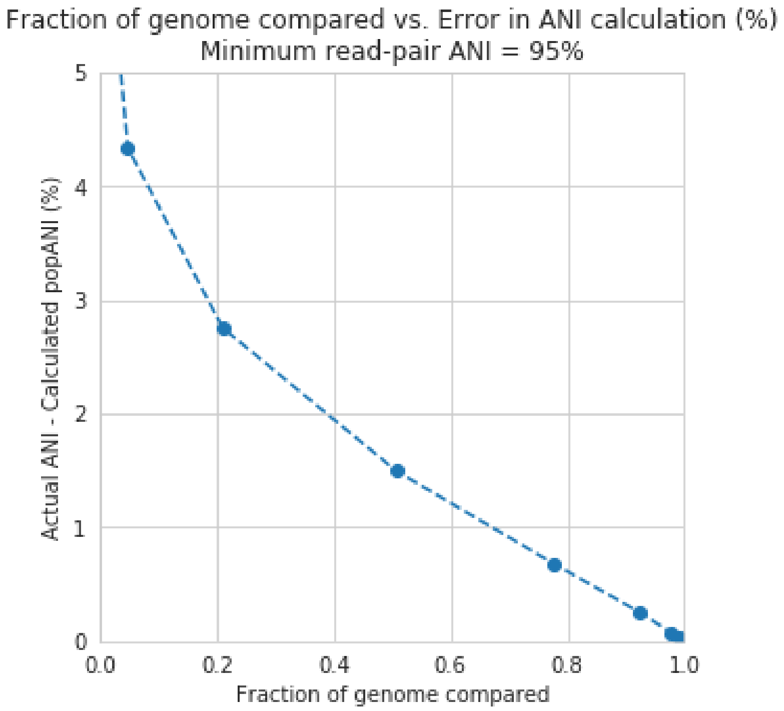
Fraction of genome compared vs. Error in ANI calculation (%)
Minimum read-pair ANI = 95%
Actual ANI - Calculated popANI (%)
Fraction of genome compared
0.0
0.2
0.4
0.6
0.8
1.0
0
1
2
3
4
5

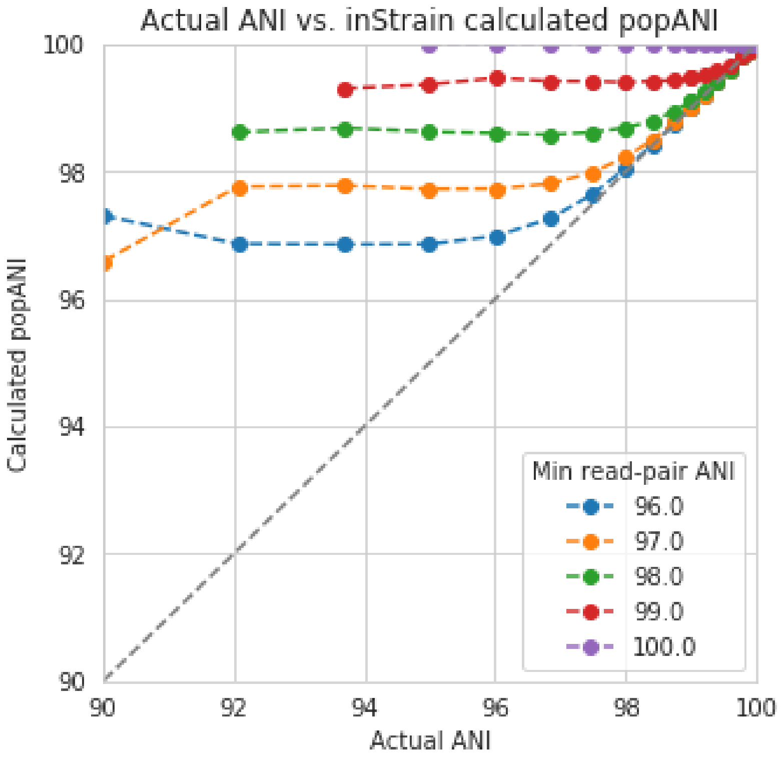
Actual ANI vs. inStrain calculated popANI
Calculated popANI
Actual ANI
Min read-pair ANI
96.0
97.0
98.0
99.0
100.0
90
92
94
96
98
100

There are a couple of things to point out here.

1) Having a lower minimum read-pair ANI cutoff lets you accurately detect more distant ANI values. This makes sense given the logic above.
2) There is a ceiling to how much the ANI is overestimated. If your minimum read-pair ANI is 96%, you think even very distantly related things have an ANI of ~96.5% ANI. If the minimum ANI threshold is 98%, you think distantly related things are ~98.5% ANI.
3) To get an accurate ANI value, you need to set your minimum read-pair ANI cutoff significantly below the ANI value that you wish to detect.

All of this begs the question, why would you ever set your minimum ANI threshold above 90% or so? If you're comparing clonal genomes, that would be a good idea. However, in most real scenarios, you want to set your minimum ANI threshold as high as possible to avoid mis-mapped reads, which will artificially increase your reported popANI.

Finally, this brings is to perhaps the most confusing yet import figure of this whole section. If I want to identify nearly identical genomes in two samples, what should I set my minimum ANI threshold to?

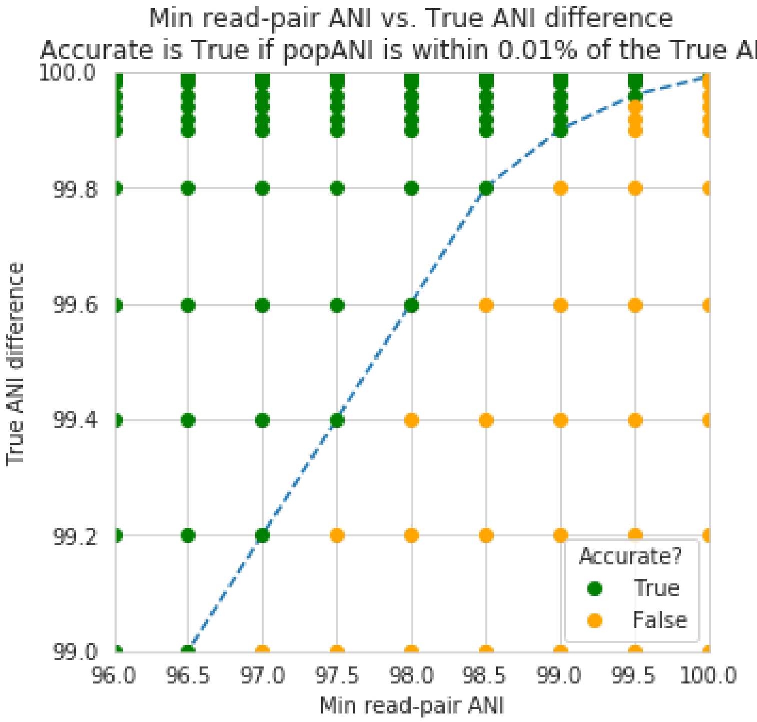

The above figure shows a range of minimum read-pair ANI thresholds on the x-axis, and a range of True ANI differences between genomes on the y-axis. Dots are colored green if the reported popANI is within 0.01% ANI of the True ANI, and colored yellow if they are not. As you can see, when you want to identify genomes that are extremely closely related (>99.9%), pretty much all minimum read-pair ANI thresholds values work. This is because if the genomes are that similar, there are going to be few reads that are thrown out due to have too many SNPs. This figure looks a bit more odd when you consider an "accurate" comparison to be one with 0.001% of the actual ANI

However, you also need to keep in mind that you want to have high breadth of coverage for each of the reads mapped to the reference genome. If the reference genome is not perfect, you need to relax your ANI threshold even more

**Note:** In conclusion: If you have a reference genome that closely represents the true organism, and you want to identify extremely similar genomes (>99.999% ANI), a minimum read-pair ANI threshold of 98% is probably good.

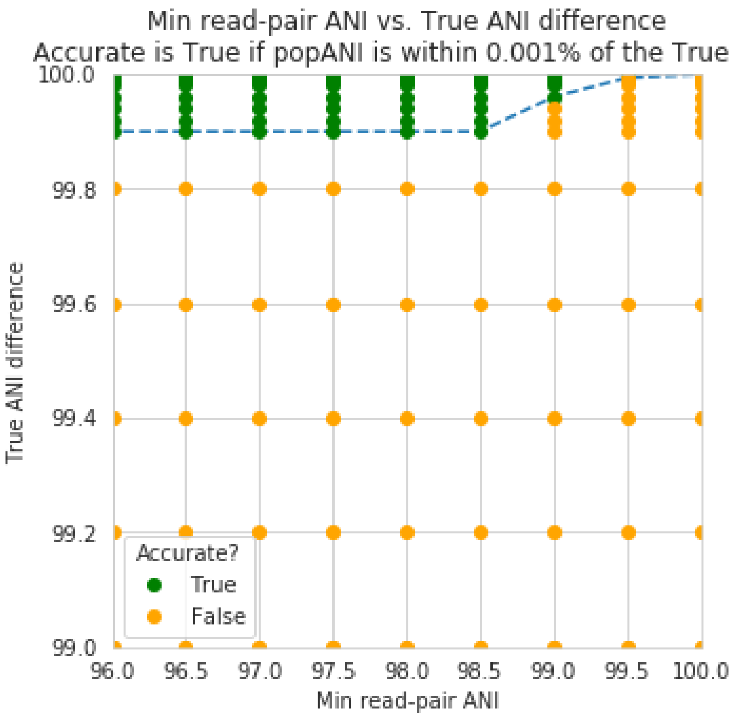

If you are working with a de-replicated set of genomes that you're mapping to, however (as recommended above), a minimum read-pair ANI threshold of 95% is probably better.

##### 1.6.2 Accessing raw data

inStrain stores much more data than is shown in the output folder. It is kept in the `raw_data` folder, and is mostly stored in compressed formats (see the section "Descriptions of raw data" for what kinds of data are available). This data can be easily accessed using python, as described below.

To access the data, you first make an SNVprofile object of the inStrain output profile, and then you access data from that object. For example, the following code accessed the raw SNP table

```
import inStrain
import inStain.SNVprofile

IS = inStain.SNVprofile.SNVprofile(``/home/mattolm/inStrainOutputTest/``)
raw_snps = IS.get('raw_snp_table')
```

You can use the example above (`IS.get()`) to access any of the raw data described in the following section. There are also another special things that are accessed in other ways, as described in the section "Accessing other data"

###### Basics of raw_data

A typical run of inStrain will yield a folder titled "raw_data", with lots of individual files in it. The specifics of what files are in there depend on how inStrain was run, and whether or not additional commands were run as well (like profile_genes).

There will always be a file titled "attributes.tsv". This describes some basic information about each item in the raw data. Here's an example:

```
name	value	type	description
location	/Users/mattolm/Programs/strains_analysis/test/test_data/N5_271_010G1_
↪scaffold_min1000.fa-vs-N5_271_010G2.sorted.bam.v6.IS	value	Location of␣
↪SNVprofile object
version	0.6.0	value	Version of inStrain
bam_loc	N5_271_010G1_scaffold_min1000.fa-vs-N5_271_010G2.sorted.bam	value	␣
↪Location of .bam file
scaffold_list	/home/mattolm/Bio_scripts/TestingHouse/N5_271_010G1_scaffold_min1000.fa-
↪vs-N5_271_010G2.sorted.bam.v6.IS/raw_data/scaffold_list.txt	list	1d list of␣
↪scaffolds, in same order as counts_table
counts_table	/home/mattolm/Bio_scripts/TestingHouse/N5_271_010G1_scaffold_min1000.fa-
↪vs-N5_271_010G2.sorted.bam.v6.IS/raw_data/counts_table.npz	numpy	1d numpy␣
↪array of 2D counts tables for each scaffold
scaffold2length	/home/mattolm/Bio_scripts/TestingHouse/N5_271_010G1_scaffold_
↪min1000.fa-vs-N5_271_010G2.sorted.bam.v6.IS/raw_data/scaffold2length.json	␣
↪dictionary	Dictionary of scaffold 2 length
window_table	/home/mattolm/Bio_scripts/TestingHouse/N5_271_010G1_scaffold_min1000.fa-
↪vs-N5_271_010G2.sorted.bam.v6.IS/raw_data/window_table.csv.gz	pandas	Windows␣
↪profiled over (not sure if really used right now)
raw_linkage_table	/home/mattolm/Bio_scripts/TestingHouse/N5_271_010G1_scaffold_
↪min1000.fa-vs-N5_271_010G2.sorted.bam.v6.IS/raw_data/raw_linkage_table.csv.gz	␣
↪pandas	Raw table of linkage information
raw_snp_table	/home/mattolm/Bio_scripts/TestingHouse/N5_271_010G1_scaffold_min1000.fa-
↪vs-N5_271_010G2.sorted.bam.v6.IS/raw_data/raw_snp_table.csv.gz	pandas	Contains␣
↪raw SNP information on a mm level
cumulative_scaffold_table	/home/mattolm/Bio_scripts/TestingHouse/N5_271_010G1_
↪scaffold_min1000.fa-vs-N5_271_010G2.sorted.bam.v6.IS/raw_data/cumulative_scaffold_
↪table.csv.gz	pandas	Cumulative coverage on mm level. Formerly scaffoldTable.
↪csv
cumulative_snv_table	/home/mattolm/Bio_scripts/TestingHouse/N5_271_010G1_scaffold_
↪min1000.fa-vs-N5_271_010G2.sorted.bam.v6.IS/raw_data/cumulative_snv_table.csv.gz	␣
↪pandas	Cumulative SNP on mm level. Formerly snpLocations.pickle
scaffold_2_mm_2_read_2_snvs	/home/mattolm/Bio_scripts/TestingHouse/N5_271_010G1_
↪scaffold_min1000.fa-vs-N5_271_010G2.sorted.bam.v6.IS/raw_data/scaffold_2_mm_2_read_
↪2_snvs.pickle	pickle	crazy nonsense needed for linkage
covT	/home/mattolm/Bio_scripts/TestingHouse/N5_271_010G1_scaffold_min1000.fa-vs-N5_
↪271_010G2.sorted.bam.v6.IS/raw_data/covT.hd5	special	Scaffold -> mm ->␣
↪position based coverage
snpsCounted	/home/mattolm/Bio_scripts/TestingHouse/N5_271_010G1_scaffold_min1000.fa-
↪vs-N5_271_010G2.sorted.bam.v6.IS/raw_data/snpsCounted.hd5	special	Scaffold ->␣
↪mm -> position based True/False on if a SNPs is there
clonT	/home/mattolm/Bio_scripts/TestingHouse/N5_271_010G1_scaffold_min1000.fa-vs-N5_
↪271_010G2.sorted.bam.v6.IS/raw_data/clonT.hd5	special	Scaffold -> mm ->␣
↪position based clonality
read_report	/home/mattolm/Bio_scripts/TestingHouse/N5_271_010G1_scaffold_min1000.fa-
↪vs-N5_271_010G2.sorted.bam.v6.IS/raw_data/read_report.csv.gz	pandas	Report on␣
↪reads
```

This is what the columns correspond to:

**name** The name of the data. This is the name that you put into `IS.get()` to have inStrain retrieve the data for you. See the section "Accessing raw data" for an example.

**value** This lists the path to where the data is located within the raw_data folder. If the type of data is a value, than this just lists the value

**type** This describes how the data is stored. Value = the data is whatever is listed under value; list = a python list;

numpy = a numpy array; dictionary = a python dictionary; pandas = a pandas dataframe; pickle = a piece of data that's stored as a python pickle object; special = a piece of data that is stored in a special way that inStrain knows how to de-compress

**description** A one-sentence description of what's in the data.

> **Warning:** Many of these pieces of raw data have the column "mm" in them, which means that things are calculated at every possible read mismatch level. This is often not what you want. See the section "Dealing with mm" for more information.

##### Accessing other data

In addition to the raw_data described above, there are a couple of other things that inStrain can make for you. You access these from methods that run on the IS object itself, instead of using the `get` method. For example:

```
import inStrain
import inStain.SNVprofile

IS = inStain.SNVprofile.SNVprofile(``/home/mattolm/inStrainOutputTest/``)
coverage_table = IS.get_raw_coverage_table()
```

The fellowing methods work like that:

**get_nonredundant_scaffold_table()** Get a scaffold table with just one line per scaffold, not multiple mms

**get_nonredundant_linkage_table()** Get a linkage table with just one line per scaffold, not multiple mms

**get_nonredundant_snv_table()** Get a SNP table with just one line per scaffold, not multiple mms

**get_clonality_table()** Get a raw clonality table, listing the clonality of each position. Pass *nonredundant=False* to keep multiple mms

##### Dealing with "mm"

Behind the scenes, inStrain actually calculates pretty much all metrics for every read pair mismatch level. That is, only including read pairs with 0 mis-match to the reference sequences, only including read pairs with >= 1 mis-match to the reference sequences, all the way up to the number of mismatches associated with the "PID" parameter.

For most of the output that inStrain makes in the output folder, it removes the "mm" column and just gives the results for the maximum number of mismatches. However, it's often helpful to explore other mismatches levels, to see how parameters vary with more or less stringent mappings. Much of the data stored in "read_data" is on the mismatch level. Here's an example of what the looks like (this is the cumulative_scaffold_table):

```
,scaffold,length,breadth,coverage,median_cov,std_cov,bases_w_0_coverage,mean_
↪clonality,median_clonality,unmaskedBreadth,SNPs,expected_breadth,ANI,mm
0,N5_271_010G1_scaffold_102,1144,0.9353146853146853,5.106643356643357,5,2.
↪932067325774674,74,1.0,1.0,0.6145104895104895,0,0.9889923642060382,1.0,0
1,N5_271_010G1_scaffold_102,1144,0.9353146853146853,6.421328671328672,6,4.
↪005996333777764,74,0.9992001028104149,1.0,0.6748251748251748,0,0.9965522492489882,1.
↪0,1
2,N5_271_010G1_scaffold_102,1144,0.9423076923076923,7.3627622377622375,7,4.
↪2747074564903285,66,0.9993874800638958,1.0,0.7928321678321678,0,0.998498542620078,1.
↪0,2
3,N5_271_010G1_scaffold_102,1144,0.9423076923076923,7.859265734265734,8,4.
↪748789115369562,66,0.9992251555869703,1.0,0.7928321678321678,0,0.9990314705263914,1.
↪0,3
```

(continues on next page)

(continued from previous page)

```
4,N5_271_010G1_scaffold_102,1144,0.9423076923076923,8.017482517482517,8,4.
↪952541407151938,66,0.9992251555869703,1.0,0.7928321678321678,0,0.9991577528529144,1.
↪0,4
5,N5_271_010G1_scaffold_102,1144,0.9458041958041958,8.271853146853147,8,4.
↪9911156795536105,62,0.9992512780077317,1.0,0.8024475524475524,0,0.9993271891539499,
↪1.0,7
```

As you can see, the same scaffold is shown multiple times, and the last column is `mm`. At the row with mm = 0, you can see what the stats are when only considering reads that perfectly map to the reference sequence. As the mm goes higher, so do stats like coverage and breadth, as you now allow reads with more mismatches to count in the generation of these stats. In order to convert this files to what is provided in the output folder, the following code is run:

```
import inStrain
import inStain.SNVprofile

IS = inStain.SNVprofile.SNVprofile(``/home/mattolm/inStrainOutputTest/``)
scdb = IS.get('cumulative_scaffold_table')
ScaffDb = scdb.sort_values('mm')\
            .drop_duplicates(subset=['scaffold'], keep='last')\
            .sort_index().drop(columns=['mm'])
```

The last line looks complicated, but it's very simple what is going on. First, you sort the database by `mm`, with the lowest mms at the top. Next, for each scaffold, you only keep the row with the lowest mm. That's done using the `drop_duplicates(subset=['scaffold'], keep='last')` command. Finally, you re-sort the DataFrame to the original order, and remove the `mm` column. In the above example, this would mean that the only row that would survive would be where mm = 7, because that's the bottom row for that scaffold.

You can of course subset to any level of mismatch by modifying the above code slightly. For example, to generate this table only using reads with <=5 mismatches, you could use the following code:

```
import inStrain
import inStain.SNVprofile

IS = inStain.SNVprofile.SNVprofile(``/home/mattolm/inStrainOutputTest/``)
scdb = IS.get('cumulative_scaffold_table')
scdb = scdb[scdb['mm'] <= 5]
ScaffDb = scdb.sort_values('mm')\
            .drop_duplicates(subset=['scaffold'], keep='last')\
            .sort_index().drop(columns=['mm'])
```

**Warning:** You usually do not want to subset these DataFrames using something like `scdb = scdb[scdb['mm'] == 5]`. That's because if there are no reads that have 5 mismatches, as in the case above, you'll end up with an empty DataFrame. By using the drop_duplicates technique described above you avoid this problem, because in the cases where you don't have 5 mismatches, you just get the next-highest mm level (which is usually what you want)

Performance issues +————

inStrain uses a lot of RAM. In the log file, it often reports how much RAM it's using and how much system RAM is available. To reduce RAM usage, you can try the following things:

- Use the `--skip_mm` flag. This won't profile things on the mm level (see the above section), and will treat every read pair as perfectly mapped
- Use `quick_profile` to figure out which scaffolds actually have reads mapping to them, and only run inStrain

on those

##### 1.6.3 A note for programmers

If you'd like to edit inStrain to add functionality for your data, don't hesitate to reach out to the authors of this program for help. Additionally, please consider submitting a pull request on GitHub so that others can use your changes as well.
